# Harbouring *Starships*: The accumulation of large Horizontal Gene Transfers in Domesticated and Pathogenic Fungi

**DOI:** 10.1101/2024.07.03.601904

**Authors:** Samuel O’Donnell, Gabriela Rezende, Jean-Philippe Vernadet, Alodie Snirc, Jeanne Ropars

**Affiliations:** Universite Paris-Saclay, CNRS, AgroParisTech, Ecologie Systématique et Evolution, 91190, Gif-sur-Yvette, France

**Keywords:** adaptation, transposable elements, moulds, filamentous fungi, fermented food, clinical, *Penicillium*, *Aspergillus*, human-made environments, accurate phylogeny, HGT

## Abstract

Human-related environments, including food and clinical settings, present microorganisms with atypical and challenging conditions that necessitate adaptation. Several cases of novel horizontally acquired genetic material associated with adaptive traits have been recently described, contained within giant transposons named *Starships.* While a handful of *Starships* have been identified in domesticated species, their abundance has not yet been systematically explored in human-associated fungi. Here, we investigated whether *Starships* have shaped the genomes of two major genera of fungi occurring in food and clinical environments, *Aspergillus* and *Penicillium.* Using seven independent domestication events, we found in all cases that the domesticated strains or species exhibited significantly greater *Starship* content compared with close relatives from non-human-related environments. We found a similar pattern in clinical contexts. Our findings have clear implications for agriculture, human health and the food industry as we implicate *Starships* as a widely recurrent mechanism of gene transfer aiding the rapid adaptation of fungi to novel environments.

## Introduction

A central question in evolutionary biology is to understand how species adapt to their environment. Indeed, identifying genomic footprints of adaptation is of interest for predicting how species can face global changes. Horizontal gene transfer (HGT) and transposable elements (TEs) are key contributors to adaptive evolution. First recognized as a major mechanism of adaptation in bacteria, it is now accepted that HGT also occurs in eukaryotes (Keeling 2024), facilitating, for example, the acquisition of both virulence genes in fungal plant pathogens (Friesen et al. 2006) and carotenoid production in aphids (Moran and Jarvik 2010). TEs are mobile genetic elements present in all domains of life, capable of moving within and between genomes through horizontal transfer events (Zhang et al. 2020). This movement generates genetic variability and genome plasticity, fostering the emergence of novel adaptive variants, particularly in species coping with new environments (Schrader and Schmitz 2019).

Fungi are models for studying adaptation in eukaryotes given their relatively small genomes and their experimental assets. Additionally, some have been domesticated by humans to ferment food (e.g. beer, bread, sake, tamari, cheese), for industrial purposes (enzyme and protein synthesis) and to produce antibiotics (penicillin). Considering domestication is the result of strong and recent selection by humans on some phenotypic traits, these domesticated fungi are prime models for looking at genomic footprints of adaptation as they should be still visible in the genomes. Horizontal gene transfers have been identified as a recurrent genomic mechanism involved in adaptation in domesticated fungi, in the budding yeast *Saccharomyces cerevisiae* (Novo et al. 2009; Legras et al. 2018), in the dry-cured meat *Penicillium* species *P. nalgiovense* and *P. salamii* (Lo et al. 2023) as well as in the cheese species *P. camemberti* and *P. roqueforti* (Cheeseman et al. 2014; Ropars et al. 2015). The very same HGTs have been identified in a variety of distantly related *Penicillium* fungi used for both cheese and meat production giving us cases of convergent adaptation (Cheeseman et al. 2014; Ropars et al. 2015; Lo et al. 2023).

Recently, these HGTs found in domesticated *Penicillium* fungi have been formally identified as elements from the superfamily of giant transposons called *Starships* (Gluck-Thaler et al. 2022; Lo et al. 2023). These large TEs, ranging from tens to hundreds of kilobases, contain a structurally conserved tyrosine recombinase with a DUF3435 domain, also called the Captain, and can house an extensive and diverse array of cargo genes not involved in transposition (Gluck-Thaler et al. 2022; Urquhart et al. 2023). Notably, these cargo genes have already been linked to, sometimes retroactively, resistance to formaldehyde (A.S. Urquhart et al. 2024), heavy metals (Urquhart et al. 2022; Urquhart et al. 2023), adaptation to food production (Cheeseman et al. 2014; Ropars et al. 2015; Lo et al. 2023) and plant pathogenicity (Peck et al. 2023; Bucknell et al. 2024), suggesting that *Starships* and their associated cargo may facilitate rapid adaptation to challenging environments. This may be particularly relevant to both newly emerging and treatment resistant pathogens within the *Aspergillus* genus.

Due to both their size and the frequent presence of repetitive material, *Starships* are often fragmented in short-read assembled genomes and the analysis of their structure and impact can be hindered (Cheeseman et al. 2014; Gluck-Thaler et al. 2022; Bucknell et al. 2024). Fortunately, long-read assembled genomes have enabled the discovery and description of complete *Starship* elements along with their surrounding genomic context (Vogan et al. 2021; Gluck-Thaler et al. 2022; Bucknell et al. 2024). Here, we focused on the fungal genera *Penicillium* and *Aspergillus*, each comprising various species of considerable significance to humans, to test whether human-related strains, compared to environmental, have accumulated a greater amount of accessory material from *Starships* insertions. These fungi play roles in the production of cheese (e.g. *Penicillium camemberti* for the production of soft cheeses and *P. roqueforti* for blue cheeses), cured sausages (e.g. *P. nalgiovense* and *P. salamii*) and both soy and rice based products such as soy sauce, miso and alcohols (e.g. *Aspergillus oryzae*, *A*. *sojae* and *A. luchuensis*). Other species included problematic human pathogens (e.g. *A. fumigatus*, *A. udagawae* and *A. felis*).

To do so, we first compiled a genome database containing more than 1,600 public and newly-assembled genomes, manually curated information on the isolation origins of each strain and generated phylogenies using >3000 single copy orthologs to ensure accurate species identification. We then used starfish (Gluck-Thaler and Vogan 2024) to identify tyrosine recombinase/Captain genes within all these genomes, using their abundance as a proxy for the number of *Starships* present. All strains or species associated with food production or clinical contexts exhibited significantly greater *Starship* content compared with closely related strains or species from environmental sources. Second, we developed a genome-graph based pipeline to extract both complete *Starships* and large *Starship-like* regions from long-read genomes. We found that *Starships* were mostly unique, suggesting independent acquisition events, but that same cargo genes were found in different *Starships.* Importantly, specific gene functions were enriched in *Starships* of domesticated fungi, relevant with adaptation to food. This includes several functionally relevant cargo clusters, present across multiple *Starships*, thus greatly enhancing the list of adaptive convergence cases between distinct lineages. Additionally, we unveiled a novel element present in a subset of *Starships*, a MYB/SANT gene, often located in *Starships* at the opposite edge to the Captain gene, therefore likely denoting the downstream border to these *Starships*. In summary, our findings corroborate the building evidence that *Starships* play a distinguished role in rapid adaptation of fungi to novel environments, including those with human significance.

## Results

### Accumulation of *Starships* in human-associated fungal isolates

Using 1516 (368 *Penicillium* and 1148 *Aspergillus*) publicly assembled genomes and 157 newly assembled *Penicillium* genomes, alongside manually curated isolation origins, we tested whether food and clinical environments have any discernible relationship to *Starship* content (**table S1**). Due to the limitations in using short-read generated assemblies, which make up the vast majority of publicly available genomes, we use solely the number of identified DUF3435-domain containing genes (hereafter named Captains) as a proxy for the number of *Starship* insertions considering their conserved and unique position in *Starships* (Gluck-Thaler et al. 2022; Gluck-Thaler and Vogan 2024).

To ensure the accuracy of the species identification for all genomes used in this study, we also constructed BUSCO-based phylogenetic trees for each genus (**Fig. 1**; **fig. S3**; **fig. S4**; **fig. S5**; **fig. S6**; **table S1**; **File S1**; **File S2**). Using these genus wide datasets, we see greater *Starship* content on average in strains isolated from food, saline and clinical contexts compared to environmental (p < 0.0001; **Fig. 1B**; **Fig. 1D**).

**Figure 1.**
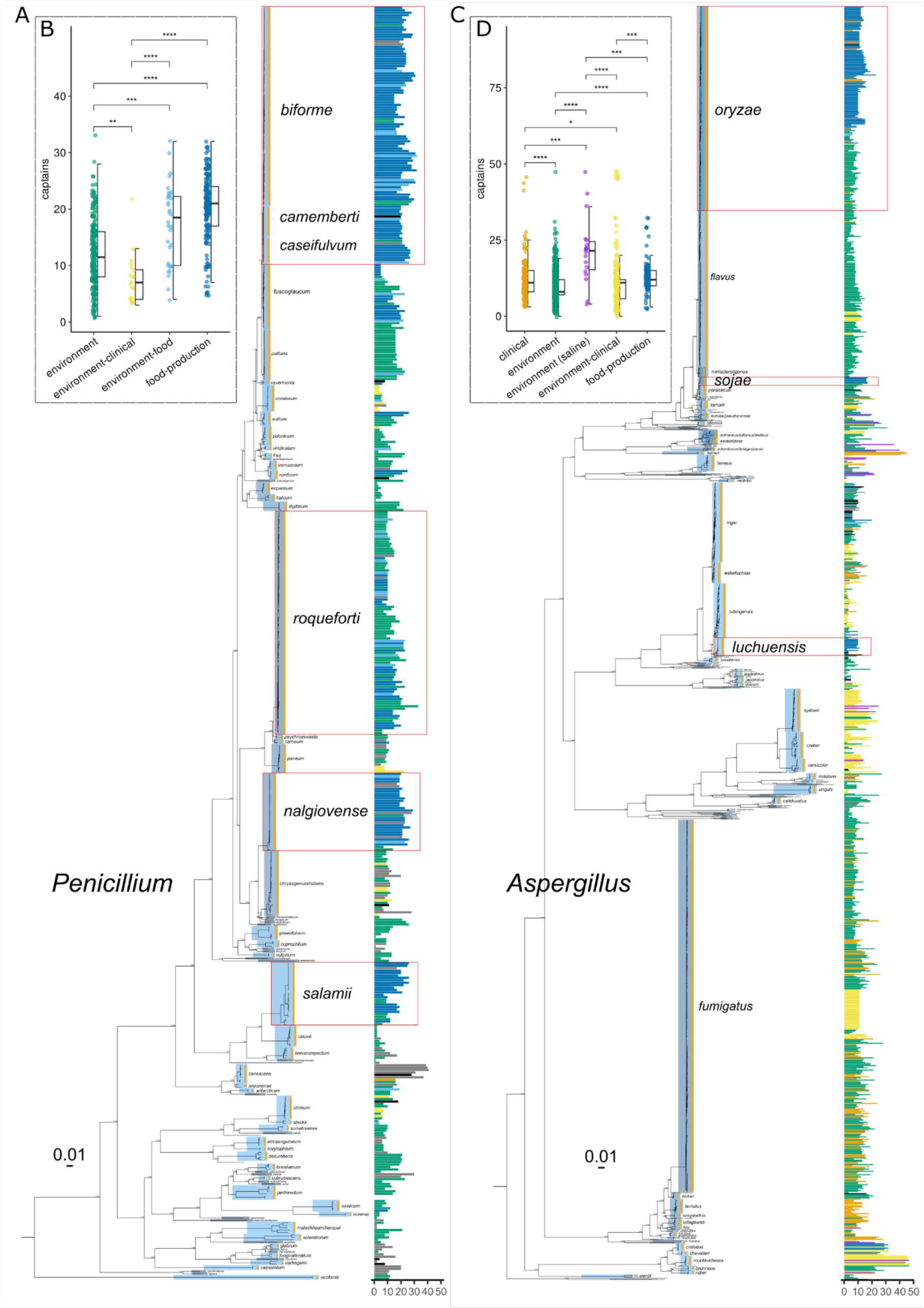
Greater S*tarship* content in clinical, food-related and saline-environmental strains with phylogenetic evidence of species identification. Captain count per strain and isolation category along the phylogeny of *Penicillium* and *Aspergillus* using 525 and 1148 genome assemblies respectively. (A) 3154 single copy BUSCO genes were extracted from each assembly and used to generate a Maximum-Likelihood tree with SH-like support. Tree was rooted using *Aspergillus* genomes which have been removed. (B) 3258 single copy BUSCO genes were extracted from each assembly and used to generate a Maximum-Likelihood tree with SH-like support. Tree was rooted using *Penicillium* genomes which have been removed. (A and B) Trees are labelled by species specific nodes and scale bars represent 0.01 substitutions per site for branch lengths. Red outlines encompass seven examples of species independently domesticated for food production. Each assembly forms a row that contains a bar on the right with the number of Captain genes detected, coloured by isolation category (clinical = orange; environment = green; environment (saline) = purple; environment-clinical = yellow; environment-food = light-blue; food-production=dark-blue; unknown=grey; NA=black). (B and C) Boxplots and dotplots of the number of Captains per assembly split by isolation category for their respective genus. P-values for the differences between groups were calculated using Wilcoxon rank-sum tests with Bonferroni adjusted p-values for multiple tests (* = p <= 0.05; ** = p <= 0.01; *** = p <= 0.001; **** = p <= 0.0001). Non-significant differences are not labelled.

Importantly, in order to confirm that we are indeed detecting, in some cases, horizontally transferred Captains located within a *Starship*, we looked at the discrepancies between the pairwise identity of Captain proteins and the average amino acid identity (AAI) of their respective genomes estimated using 50 BUSCO proteins. We only analysed Captains with a pairwise identity >=95% and generated a non-redundant dataset, accounting for species, identity and Captain length, comprising 9821 Captain proteins (**fig. S7**). We then visualised alignments between genomes with large differences between their AAI and Captain identity to show that Captains are associated with regions with similarity unexplained by vertical inheritance (**fig. S7**).

#### Accumulation of *Starships* in moulds domesticated for food-production

We have previously shown that three *Penicillium* species have been domesticated for cheesemaking, namely *P. roqueforti* used for blue cheeses worldwide such as Roquefort or Stilton (Dumas et al. 2020), *P. camemberti* used for soft cheeses such as Camembert (Ropars et al. 2020) and *P. biforme* present on many natural cheese rinds and also found on some dried sausages (Ropars et al. 2020; Lo et al. 2023), with *P. biforme* and *P. camemberti* being sister clades. Our analysis shows that *P. biforme* and the two varieties of *P. camemberti;* i.e. var. *camemberti* and var. *caseifulvum*, contain significantly more *Starships* than their wild sister species *P. fuscoglaucum* and *P. palitans* (p <= 0.026; **Fig. 2A**; **fig. S8**). *Penicillium roqueforti* has been domesticated independently at least twice (Dumas et al. 2020), with one population containing strains of Roquefort PDO cheeses (named Roquefort population) and another population containing strains from all blue cheeses produced worldwide (named Non-Roquefort population). A new cheese population has been recently described (Crequer et al. 2023), with strains isolated from an uninoculated blue cheese from the village Termignon in the French Alps (named the Termignon population). The two HGTs identified previously in the non-Roquefort population (*cheesyTer* (Ropars et al. 2015) and *Wallaby* (Cheeseman et al. 2014), retroactively identified as *Starships* (Gluck-Thaler et al. 2022; Urquhart et al. 2023)), have also been identified in the Termignon population. We find that strains from the wood/lumber and silage populations have significantly fewer *Starships* than cheese-related strains from both the Non-Roquefort and Termignon populations (p < =0.027; **Fig. 2B**; **fig. S9**). However, we also see that the Roquefort population has significantly fewer *Starships* than other cheese strains (p <= 0.01). We newly defined a ‘Contaminants’ population, comprising strains isolated from contaminated food products, with a *Starship* count similar to the Non-Roquefort and Termignon clades.

**Figure 2.**
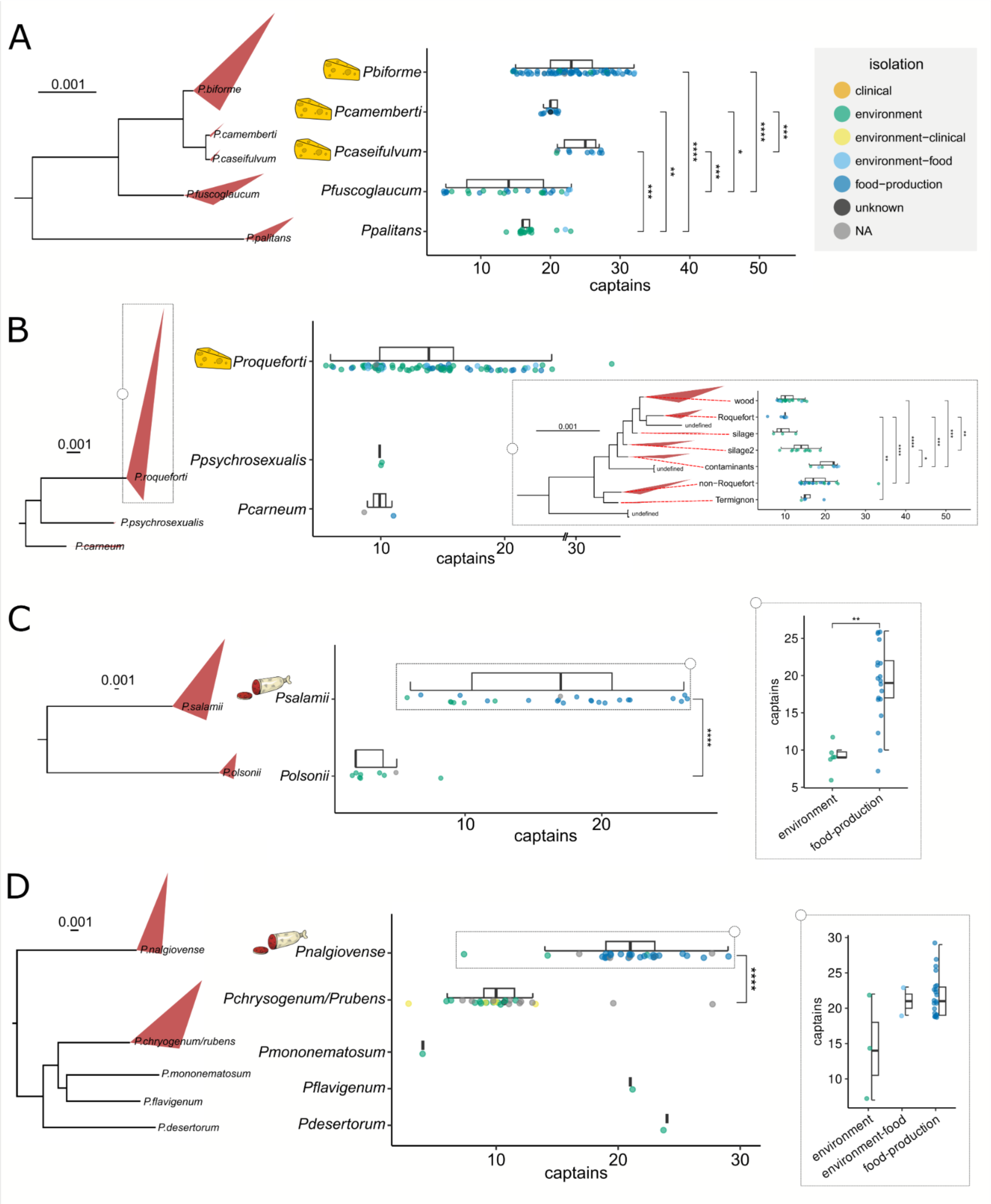
Domesticated *Penicillium* species compared by Captain count to their close relatives and split by isolation origins. Each row contains a specific subset of species shown in the phylogeny on the far left (maximum likelihood tree). The species specific nodes display a triangle with corners extending to both the maximum and minimum of the branch lengths. Scale s of branch lengths are displayed above each phylogenetic tree on the left panel. Boxplots and dotplots show the Captain count distribution with the dots coloured by isolation origins. Dotted squares on the far right of rows B, C and D, display a subset of information similarly highlighted to their left. P-values for the differences between groups were calculated using Wilcoxon rank-sum tests with Bonferroni adjusted p-values for multiple tests (* = p <= 0.05; ** = p <= 0.01; *** = p <= 0.001; **** = p <= 0.0001). Non-significant differences are not labelled.

In dry-cured meats, two *Penicillium* species are isolated in most cases, namely *P. nalgiovense* and *P. salamii* (Iacumin et al. 2009; Perrone et al. 2015; Lo et al. 2023). Other *Penicillium* species have also been isolated from fermented meat products, including *P. biforme* (also found in cheese) and *P. nordicum.* On the contrary to *Penicillium* cheese species, dry-cured meat strains of *P. salamii* and *P. nalgiovense* are not genetically distinct from their wild counterparts when considering SNPs, but very few non-dried sausage strains were analysed (Lo et al. 2023). However, dry-cured meat strains carry HGTs, which are likely *Starships*, that are shared between all three species and absent from non-dry-cured meat strains (Lo et al. 2023). Our analysis found that *P. salamii* contains significantly more *Starships* than its wild sister species *P. olsonii* (p < 0.0001; **Fig. 2C**; **fig. S10**). Furthermore, within the *P. salamii* species, strains isolated from dry-cured meat exhibit significantly more *Starships* compared to environmentally isolated strains (p = 0.00158). For *P. nalgiovense* we show that it contains significantly more *Starships* than both sister species *P. chrysogenum* and *P. rubens* (p < 0.0001; **Fig. 2D**; **fig. S11**). As in *P. salamii, P. nalgiovense* strains isolated from dry-cured meats appear to have more *Starships* compared to their environmentally isolated counterparts.

Two distantly related species of *Aspergillus*, namely *A. sojae* and *A. oryzae,* are used for fermenting soybean-and rice-based products such as miso and sake (Gibbons et al. 2012; Acevedo et al. 2023). We show that *A. sojae* contains a significantly larger number of *Starships* compared to highly genetically similar *A. parasiticus* strains isolated from environmental sources (p = 0.00083; **Fig. 3A**; **fig. S12**). Similarly, *A. oryzae* displays a significant increase in *Starship* count when compared to strains of the closely related species *A. flavus* and *A. minisclerotigenes* (p < 0.0001; **Fig. 3B**; **fig. S13**). Moreover, within *A. oryzae*, strains with food-production origins contain significantly more *Starships* than environmentally isolated strains (p < 0.0001). Of particular note, the few clinical isolates of *A. oryzae* contain, on average, a similar number of *Starships* as the food-production isolates and were phylogenetically distinct (**fig. S13**). The last food-related *Aspergillus* species, *A. luchuensis*, primarily used in alcohol production, contains significantly more *Starships* than its closely related and primarily environmental sister species *A. tubingensis* (p = 0.00018; **Fig. 3C**; **fig. S14**).

**Figure 3.**
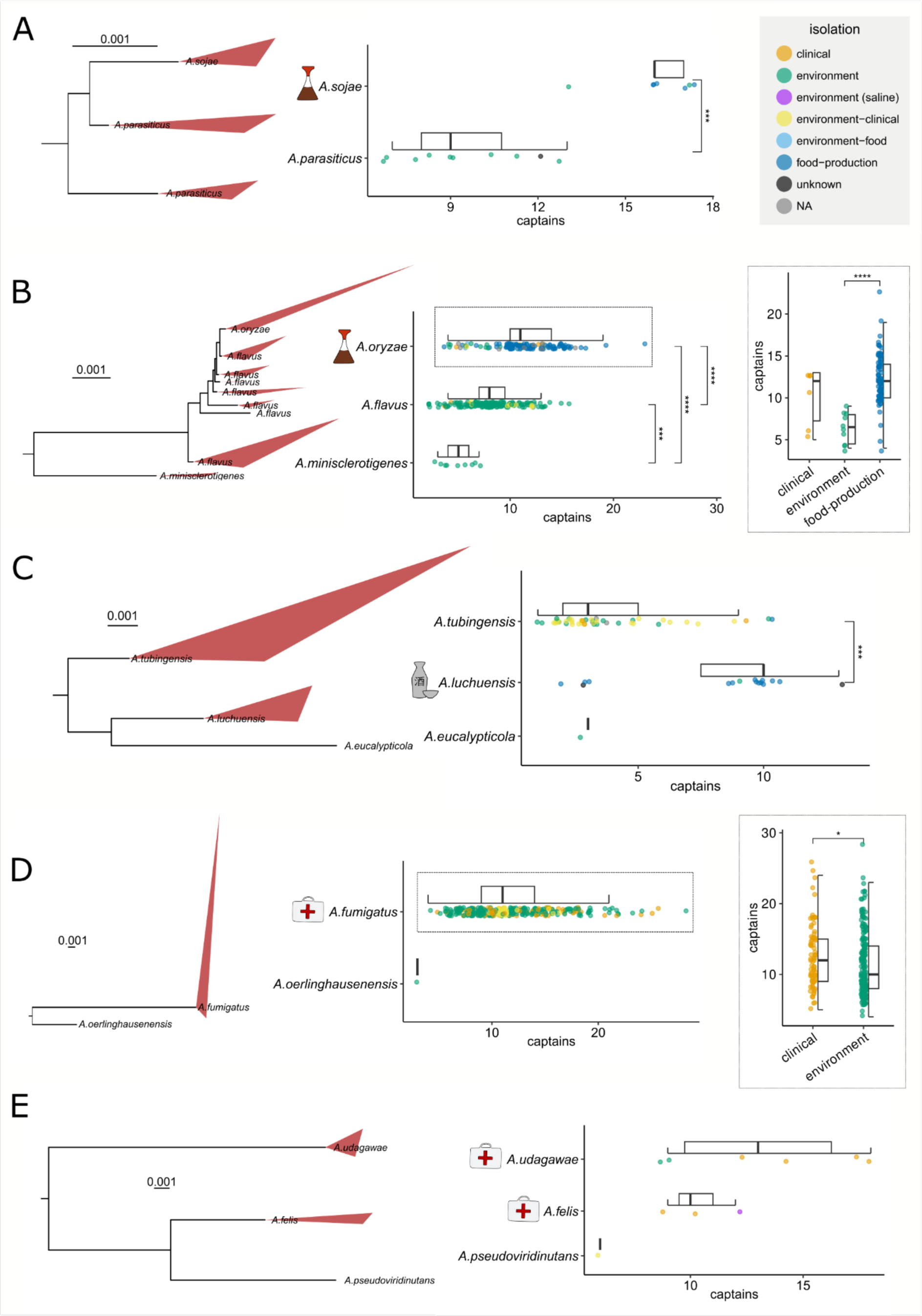
Domesticated and pathogenic *Aspergillus* species compared by Captain count to their close relatives and split by isolation origins. Each row contains a specific subset of species shown in the maximum likelihood phylogenetic tree on the far left. Scales of branch lengths are displayed above each phylogenetic tree on the left panel. The species specific nodes display a triangle with corners extending to both the maximum and minimum of the branch lengths. Boxplots and dotplots show the Captain count distribution with the dots coloured by isolation origins. Dotted squares on the far right of rows B and D, display a subset of information similarly highlighted to their left. P-values for the differences between groups were calculated using Wilcoxon rank-sum tests with Bonferroni adjusted p-values for multiple tests (* = p <= 0.05; ** = p <= 0.01; *** = p <= 0.001; **** = p <= 0.0001). Non-significant differences are not labelled.

Similarly, in *A. niger,* the average number of Captains for the food-production, environment-food and clinical isolates are higher than that of environment and environment-clinical (**fig. S15**; **fig. S16**). Lastly, although *A. niger* is generally used for enzyme, protein and secondary metabolite production in industry, no industrial strains were available publically.

#### Accumulation of *Starships* in clinical isolates

Aspergillosis encompasses a spectrum of diseases caused by species within the genus *Aspergillus* and represents a significant health concern globally. Although almost 20 species have been implicated as causative agents (Pfaller et al. 2006), the majority of infections are caused by four species, namely *A. fumigatus, A. flavus, A. niger* and *A. terreus. Aspergillus fumigatus* is currently the major air-borne fungal pathogen, with a relatively high incidence and mortality, comparable to *Candida* and *Cryptococcus* species (Bongomin et al. 2017; Denning 2024). We found that clinical strains of *A. fumigatus* contain on average more *Starships* than environmental strains (p = 0.02; **Fig. 3D**). Notably, the clinical strains are phylogenetically distinct and widely distributed (**fig. S17**). *Aspergillus udagawae* and *A. felis* are two species considered emerging causes of invasive aspergillosis (Sugui et al. 2010; Barrs et al. 2013). Although *A. udagawae*, *A. felis* and its sister species *A. pseudoviridinutans* have few genomes available, those at hand hint at a potentially positive relationship between clinical appearance and *Starship* count (**Fig. 3E**; **fig. S18**). These findings suggest a role of *Starships* in the virulence and/or emergence of fungal pathogens.

Looking at *Starship* content across the entire *Aspergillus* genus, the species *A. montevidensis*/*amstelodami* stands out with an average of 46 *Captains* (range: 44-48) across nine genomes, the highest per species with more than one genome, compared to the genus median of 11 (**Fig. 1**; **fig. S19**). This species is both considered pathogenic (Fernandez-Pittol et al. 2022) and commonly found in fermented foods such as teas, meju and cocoa beans. Additionally, *A. montevidensis* is closely related to *A. cristatus*, another species domesticated for tea fermentation and with high salt tolerance (Xie et al. 2024), with an average of 30 captains.

### Accumulation of specific genes functionally relevant for food adaptation in *Starships*

Using long-read genomes we aimed to, first, confirm our earlier results and secondly, explore both the cargo content and structure of *Starships*. To do so, we employed a robust genome-graph based pipeline, using pggb and odgi (Guarracino et al. 2022; Garrison et al. 2023), to conduct clade-specific all-vs-all whole genome alignments, exclusively using contiguous long-read assembled genomes. This new approach allowed us to extract large candidate *Starship*-like regions (SLRs) from all genomes simultaneously and without reference bias. The criteria for these initial candidate regions were straightforward: they needed to be larger than 30 kb in size and absent in at least one close relative genome. These regions were then classified as a SLR if at least one *Starship*-related gene was present (See Methods section *‘Starship*-like region detection’). We performed alignments of these SLRs and further confirmed that 1) these SLRs are completely absent in some close relatives and 2) inserted at non-homologous loci in the different genomes, further supporting the horizontal acquisition of these regions (examples provided in fig. S20-28). We did not aim to further characterise these *Starship-like* regions, for example by identifying more precisely the *Starship* borders, nor require the presence of a Captain, considering fragmented assemblies, structural variation and Repeat Induced Point-mutation (RIP) can disrupt the detection of full length elements and/or *Starship* related genes (Gluck-Thaler et al. 2022).

#### Functional cargo convergence within primarily unique *Starships*

Using 76 long read assemblies from 36 species, we constructed 11 clade-specific genome-graphs and detected a total of 857 *Starship-like* regions with a combined total of 202Mb and 963 Captains (**table S2**). Of the 857 regions, 79% contained at least one Captain. With this data we were able to recapitulate what we previously discovered, i.e., that food and clinical strains consistently have a larger number of *Starships* compared to their environmental relatives, and additionally that this equates to a larger sum total of *Starship*-related material (**Fig. 4**; **fig. S29**). The importance of the ability to detect all regions associated with *Starship*-like genes, considering the variation in *Starship* size, is highlighted by certain cases where although the number of Captains were similar, the amount of *Starship*-like material could differ largely. For example, the *A. oryzae* strain KBP3, with a similar number of Captains, contains over twice as much *Starship*-like material compared to other strains (**Fig. 4A**). Additionally, we were able to assess which regions were shared between different genomes within each genome graph constructed. This highlighted that some very closely related strains have a large number of recently acquired, unique *Starships* (**Fig. 4A**; **fig. S29**). A clear example is between the two varieties of *P. camemberti,* var. *caseifulvum* and var. *camemberti*, which share 99.9% DNA identity and yet contain 12/26 and 12/21 unique Starship-like regions respectively, suggesting *Starships* have accumulated independently after the divergence of the two varieties. This result was verified by manually inspecting alignments of all *Starship*-like regions.

**Figure 4.**
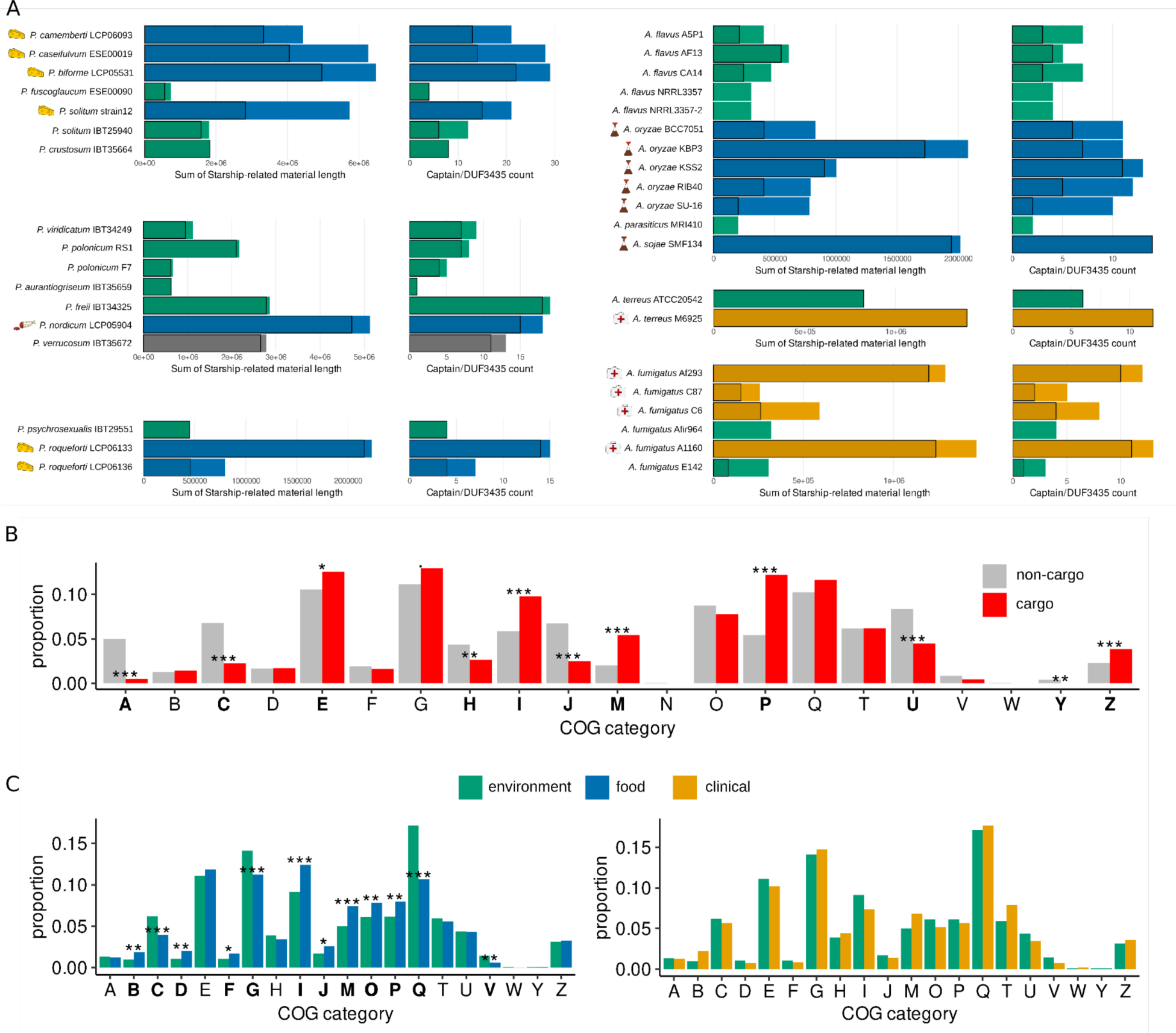
Genome graphs based *Starship* statistics and functional enrichment showed a greater number of *Starship-like* material and gene COG annotations relevant for the food environment. (A) Each set of genomes represents a genome-graph constructed with the assemblies for each strain present. The darker, black outlined column represents unique *Starship*-like material and Captains within them. (B and C) *Starship* cargo genes were analysed for functional enrichment, comparing the total proportion of each COG within the entire dataset. COG categories: A - RNA processing and modification; B - Chromatin structure and dynamics; C - Energy production and conversion; D - Cell cycle control, cell division, chromosome partitioning; E - Amino acid transport and metabolism; F - Nucleotide transport and metabolism; G - Carbohydrate transport and metabolism; H - Coenzyme transport and metabolism; I - Lipid transport and metabolism; J - Translation, ribosomal structure and biogenesis; K - Transcription; L - Replication, recombination and repair; M - Cell wall/membrane/envelope biogenesis; N - Cell motility; O - Posttranslational modification, protein turnover, chaperones; P - Inorganic ion transport and metabolism; Q - Secondary metabolites biosynthesis, transport and catabolism; R - General function prediction only; S - Function unknown; T - Signal transduction mechanisms; U - Intracellular trafficking, secretion, and vesicular transport; V - Defense mechanisms; W - Extracellular structures; Y - Nuclear structure; Z - Cytoskeleton. All p-values were calculated from hypergeometric tests and adjusted for multiple comparisons using Benjamini–Hochberg (ᐧ = p <= 0.1; * = p <= 0.05; ** = p <= 0.01; *** = p <= 0.001. (B) Cargo and non-cargo gene COG annotations for three cheese related strains (*P. caseifulvum* ESE00019, *P. camemberti* LCP6093 and *P. biforme* LCP05531). (C) Cargo gene COG annotations were grouped by strain isolation origins and compared for functional enrichment.

We then investigated whether *Starship-*cargo was functionally enriched for certain COG categories compared to non-cargo genes. We used our four newly assembled and annotated long-read assemblies from three cheese-related strains (*P. camemberti* var*. caseifulvum* ESE00019, *P. camemberti* var*. camemberti* LCP06093 and *P. biforme* LCP05531) and a closely related environmental strain (*P. fuscoglaucum* ESE00090). Using all COGs, and identifying genes without any COGs, we saw a very strong enrichment in genes without a COG annotation in all three genomes (**fig. S30**). In our cheese related strains we also see the depletion of several other COG categories. Only the Z (Cytoskeleton) category was enriched in *P. biforme*. We then filtered out COGs that were absent, unknown (S), or associated with TEs (L and K) and found, only in cheese strains, several COGs significantly enriched in cargo genes compared to the non-cargo gene dataset (**fig. S31**). Combining the data from all three genomes showed the same COG differences (**Fig. 4B**). The enriched COGs are E (Amino acid transport and metabolism), I (Lipid transport and metabolism), M (Cell wall/membrane/envelope biogenesis), P (Inorganic ion transport and metabolism) and Z, whilst there is a depletion of COGs A (RNA processing and modification), C (Energy production and conversion), H (Coenzyme transport and metabolism), J (Translation, ribosomal structure and biogenesis), U (Intracellular trafficking, secretion, and vesicular transport) and Y (Nuclear structure).

We also tested cargo COGs depending on the isolation origins of the strain by using all annotated long-read assemblies. We compared all cargo from food, clinical and environmental isolations (**Fig 4C**). We found no significant differences between clinical and environmental cargo Interestingly, several differences were significant when comparing either clinical and environmental cargo to food-related cargo. Cargo from food-related isolates are enriched for several categories (**Fig 4C**).

#### *Starships* contain cargo-clusters for rapid adaptation to human-related environments

Using the *Starship*-like regions, we found not only genes of general interest such as those related to drug resistance (CDR1, metallo-beta-lactamase, FCR1, PDR16, ergosterol biosynthesis genes, YOR1,…), pathogenicity (RBT1, SEF1, AGS1, CAT1,…) and metal ion homeostasis (VCX1, ZRT2, SMF3,…) (**table S3.6**), but also gene clusters likely involved in lactose metabolism, dityrosine biosynthesis, salt tolerance, arsenic resistance and ethanol utilisation (**Fig. 5**; **fig. S32**; **fig. S33**; **fig. S34**; **table S3**). In all cases, the rapid loss of synteny at the edges of each cluster suggests that they have all been acquired as cargo-clusters several times by different *Starships*.

**Figure 5.**
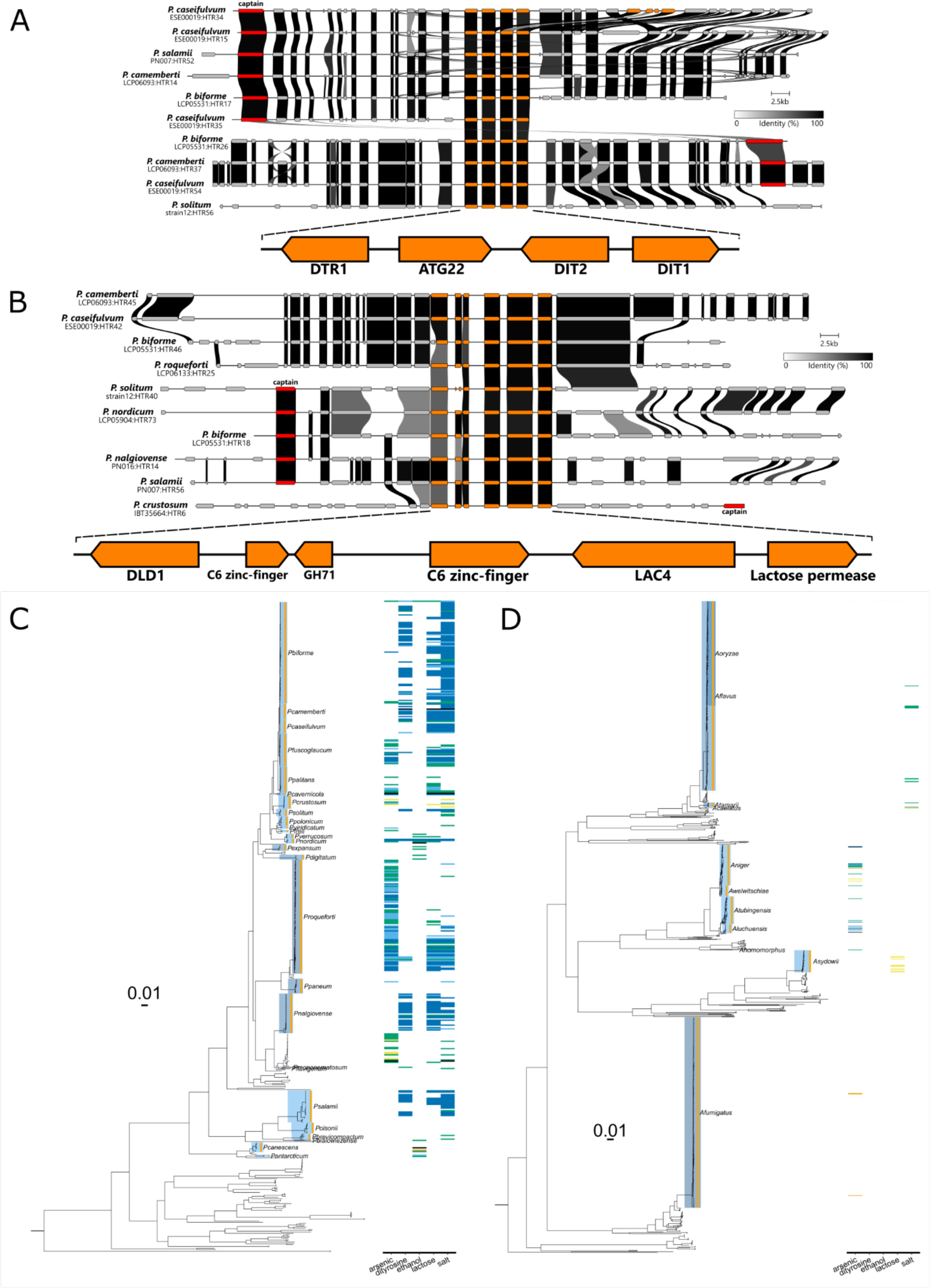
Cargo-clusters in *Starship-like* regions. Genes within a subset of *Starship-like* regions were aligned and centred upon genes within the respective gene cluster, A) a dityrosine synthesis or B) lactose metabolism clusters. Genes conserved within the cluster are highlighted in orange. Captains are coloured in red. Colour of links between genes indicate pairwise identity as indicated in the scale. The presence/absence of each cluster within all genomes in a heatmap aligned against the BUSCO based phylogeny for C) *Penicillium* and D) *Aspergillus*. The colour for each block in the heatmap indicates presence and the isolation origins classification (clinical = orange; environment = green; environment (saline) = purple; environment-clinical = yellow; environment-food = light-blue; food-production=dark-blue; unknown=grey; NA=black).

### Lactose metabolism and dityrosine synthesis clusters in cheese and cured-meat *Penicillium* species acquired independently several times in different *Starships*

We previously discovered *CheesyTer*, a retroactively identified *Starship* (Ropars et al. 2015; Gluck-Thaler et al. 2022; Urquhart et al. 2023), shared between species primarily used for cheese production, and containing two genes, a beta-galactosidase and a lactose permease, likely important for lactose consumption (Ropars et al. 2015). Here, we were able to identify that the beta-galactosidase (LAC4) and the lactose permease are actually contained within a larger lactose related gene cluster. The other genes within this cluster are two C6 zinc-fingers, a glycoside hydrolase of the family 71 (GH71) and a D-lactate dehydrogenase (DLD1) (**Fig. 5**), likely play a direct role in lactose metabolism (**table S3.1**), and all of which were present in *CheesyTer* (Ropars et al. 2015). Although our current sampling of this cluster suggests all genes are core, there is an apparent lower identity and truncated reading frames in both DLD1 and one of the two C6-zinc fingers. In addition to the previously described species *P. roqueforti*, *P. camemberti*, *P. biforme* and *P. nalgiovense*, we also found this gene cluster in genomes from *P. solitum*, *P. salamii*, *P. nordicum* and *P. crustosum,* which are all found in food environments, although *P. crustosum* is often considered purely a food contaminant (Pitt and Hocking 2009). These genomes represent multiple, independent events of adaptation to food, for both cheese and cured-meat. Additionally, *P. biforme* (LCP05531) contained two copies of the cluster in different *Starship-like* regions (**Fig. 5**).

We also identified another biosynthesis cluster, namely the dityrosine cluster, containing two genes, DIT1 and DIT2 that cluster tail-to-tail as already described across a variety of ascomycetes, particularly the Saccharomycetes (Nickles et al. 2023). These genes have been shown in *Saccharomyces cerevisiae* to be involved in the biosynthesis of dityrosine within the spore cell wall and enhance their tolerance to stresses (Briza et al. 1994). In *Candida albicans*, with the same cluster, a dit2 mutant was found to be both more susceptible to the antifungal drug caspofungin and to exhibit hyphal growth in minimal media (Melo et al. 2008).

We describe for the first time a DIT1/DIT2 cluster in two distinct *Starship-like* regions (**Fig. 5**; **table S3.2**). In addition to DIT1 and DIT2, this cluster contains, DTR1, a multi-drug resistance protein and putative dityrosine transporter of the major facilitator superfamily (Felder et al. 2002), and ATG22, a vacuolar effluxer for amino acids after autophagy that primarily targets tyrosine (Yang et al. 2006). Notably, this set of four genes has been previously described as a co-regulated dityrosine cluster in *Pichia stipitis* (Jeffries and Van Vleet 2009). This cluster has been independently acquired in different *Starships*, in the cheese species *P. camemberti var. camemberti* (LCP06093), *P. camemberti var. caseifulvum* (ESE00019) and *P. biforme* (LCP05531). Of note, there are five copies of the cluster within *P. camemberti var. caseifulvum* (ESE00019), involving four different *Starships* of which one, ESE00019:HTR34, contains a duplicate copy without DTR1 and an apparent splitting of DIT2 into two ORFs (**Fig. 5**). Similar in respects to the lactose cluster, the dityrosine cluster is found in cheese and cured-meat related strains.

### Expanding upon the Arsenic resistance cluster of *Hephaestus*

Although arsenic is a naturally occurring toxic compound present ubiquitously in the environment, elevated concentrations are found in certain areas due to human-activities such as mining and agriculture (Patel et al. 2023). Arsenic resistance has evolved many times (William and Magpantay 2024), including the ACR cluster in *S. cerevisiae* (Stefanini et al. 2022) and there is also evidence for the horizontal transmission of arsenic resistance genes by large ICE elements in bacteria (Arai et al. 2019). An arsenic cluster found within the *Starship Hephaestus* contains five genes (arsH, arsC, arsB, Pho80 and arsM) and confers arsenic resistance in the environmental fungus *Paecilomyces variotii* (Urquhart et al. 2022). With a larger sampling of all arsenic clusters in *Starships,* we describe here the cluster’s variable content, made up of only three core-cluster genes, namely, arsH, arsC and arsB, and additional genes that may be frequent but not required (**fig. S32**; **table S3.3**). Among these cluster-accessory genes, several were frequently present, specifically two genes with no functional description available and a basic region leucine zipper (bZIP), potentially playing a similar role to the bZIP transcription factor ARR1/YAP8, which is both required for the transcription of ARR2 and ARR3 in *S. cerevisiae* (Wysocki et al. 2004) and activated by arsenite binding (Navarro et al. 2022). Additionally, two other cluster-accessory genes were found close to the cluster in several different *Starship-like* regions; TRX1, a thioredoxin both inhibited by arsenic and shown to confer resistance (Park 2020) and a heavy metal associated p-type ATPase, similar to pcaA, shown to confer resistance to cadmium and lead in *Hephaestus*, possibly involved in arsenic/arsenite transport (Antonucci et al. 2017). Several genomes had multiple copies of the cargo-cluster genes, whether from several unique *Starships* and/or duplications within *Starships*. Aside from the previously described species (*P. variotii*, *A. fumigatus*, *A. sydowii*, *A. primulinus* and *A. varians* (Urquhart et al. 2022)), we detected the cluster in six *Penicillium* species (*P. chrysogenum/rubens*, *P. brevicompactum, P. freii, P. crustosum, P. nordicum* and *P. biforme*) and a further one in *A. niger*. Notably, this list contains more environmental species, deviating from the other clusters that contain primarily food-related isolates, and a combination of species from both genera.

### A highly variable salt tolerance gene cluster in food-related species

We also found more extensive evidence for a cluster briefly described in Urquhart et al. 2022 within *A. aureolatus* and containing genes involved in salt tolerance (NHA1, ENA1/2 and SAT4) (**fig. S33**; **table S3.4**). This cluster contains a more varied array of accessory genes and their structure. However, focussing on ENA2 as a central component, involved in Na+ efflux to allow salt tolerance (Ariño et al. 2010)), we see other genes commonly cluster within its vicinity. The two other core genes include the previously described SAT4/HAL4, involved in salt tolerance (Mulet et al. 1999; Urquhart et al. 2022), and a TRK2-like gene involved in potassium transport (Mulet et al. 1999). In the majority of cases, both genes are found clustered with ENA2, however, their variable orientation and position relative to ENA2 exemplify this cluster’s diverse arrangement (**fig. S33**). Additionally, this cluster contains multiple instances of accessory genes such as TRK1 (Potassium transporter, activated by SAT4 and required for SAT4 salt tolerance (Mulet et al. 1999)), pacC/RIM101 (TF involved in pH signal transduction; confers salt tolerance through the regulation of ENA2-like genes (Caracuel et al. 2003)), ERG20 (involved in sterol biosynthesis shown to impact salt-tolerance (Kodedová and Sychrová 2015)), and several others (**table S3.4**). This gene cluster repertoire clearly highlights its functional role centred around ion homeostasis and salt tolerance.

Further cementing this clusters’ functional relevance is that it was found in multiple configurations and unique *Starships* in several strains all related to food-production. For example, three clusters were found each in *P. salamii* (PN007), *P. solitum* (strain12) and *P. camemberti* var*. caseifulvum* (ESE00019), that is a cured-meat and two cheese-related strains respectively. These three strains, belonging to distant species, all shared the same *Starship* with one of these cluster configurations.

### An ethanol utilisation cluster in food spoilage species

The model system of ethanol utilisation in *Aspergillus nidulans* requires both alcA and aldA, encoding for alcohol and aldehyde dehydrogenases that convert alcohol into acetate via acetaldehyde, and the transcription factor alcR, which strongly induces the expression of alcA and aldA (Felenbok et al. 2001). Alongside alcR and alcA, three other genes make up an alc gene cluster and are regulated by alcR. One of these three genes, alcS, is likely an acetate transporter (Robellet et al. 2008). We found a *Starship* gene cluster containing alcR, alcS, alcA (ADH1) and aldA (ALD5) (**fig. S34**; **table S3.5**). Additionally, ERT1/AcuK was found in several clusters, another ethanol regulated transcription factor (Gasmi et al. 2014). This cluster was found in strains of *P. nordicum, P. canescens*, *P. antarcticum*, *P. bialowiezense*, *P. digitatum*, *P. verrucosum* and *P. expansum*, of which the latter four are all related to food spoilage, primarily in fruit.

### Cargo-clusters within the whole genera

Having leveraged long-read assemblies in order to describe complete cargo-clusters, we then took a broader look at their distribution within all genomes of *Penicillium* and *Aspergillus* used in this study (**Fig. 5C**-**D**; **fig. S35**; **table S3.7**). We found in most cases the presence of each cluster within the same species, however there were three main exceptions; The lactose cluster was found in 10/39 genomes of *Aspergillus sydowii*, the arsenic cluster was found in most strains of *P. roqueforti* (deviating from the other clusters found only in food-related clades) and the salt cluster was found in 2/8 *A. tamarii* strains. Looking at the isolation origins of all strains with each cluster, more environmental strains are found containing the ethanol and the arsenic cluster, whilst the other three clusters are found primarily in strains from food, most notably the dityrosine cluster (**fig. S35**).

### A new gene identified at the opposite border to *Starship* Captains

During preliminary pairwise comparisons and manual inspection of *Starship-like* regions, we identified genes with a particular MYB/SANT protein domain that were frequently present and *Starship-like* region specific (**fig. S36**; **Table 1**). Additionally we noticed that these genes were not only frequently close to the edge of *Starship-like* regions but both in the opposite sense and at the opposite edge to the Captain, if present. Using our newly generated list of *Starship-like* regions, we generated a non-redundant dataset and then compared the distance of each *Starship*-related gene, including the MYB/SANT, to the closest edge. This showed that DUF3435 and MYB/SANT genes are on average significantly closer to an edge than all other *Starship*-related genes (p <= 0.042; **Fig. 6A**). We also looked at the relative distance of each *Starship*-related gene from each Captain, within the same *Starship-like* region, and showed that MYB/SANT genes are significantly further away than other *Starship*-related genes (p <= 0.001; **Fig. 6B**). These results affirm the idea that MYB/SANT genes, like Captains (DUF3435), appear to have a structurally conserved position at the edge of *Starships* opposite to the Captain.

**Figure 6.**
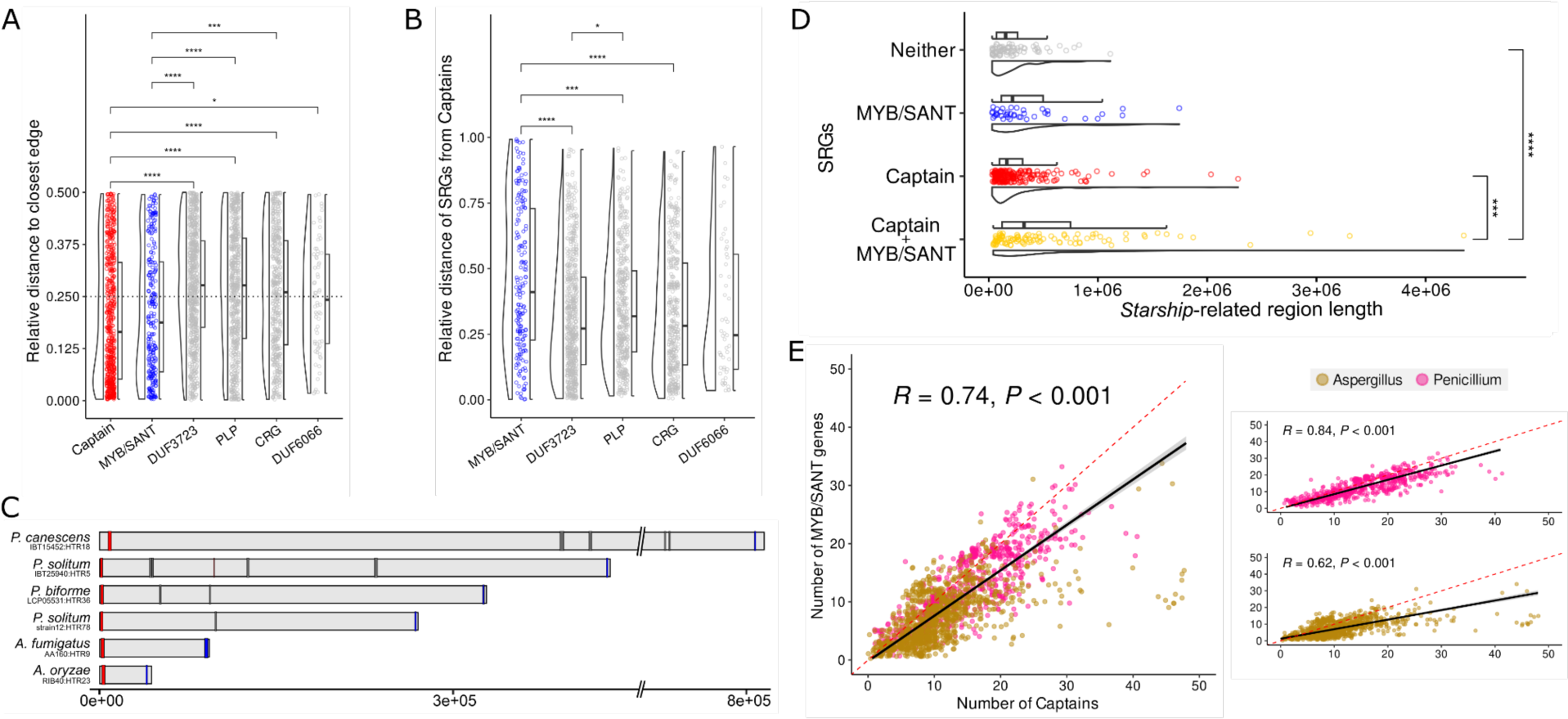
The structural association of MYB/SANT genes with *Starship-like* regions. (A) *Starship*-related genes and their relative distance to the closest edge of their associated Starship-like region. (B) The relative distances of *Starship*-related genes to Captains within the same *Starship-like* region. (C) Six examples of *Starship-like* regions indicated by grey rectangles highlighting the position of Captains (red rectangles), MYB/SANT genes (blue) and other *Starship-like* regions (grey). (D) The size of Starship-like regions that contain both a Captain and MYB/SANT gene, only a Captain, only a MYB/SANT gene or neither. (E) The linear relationship between the number of Captains and MYB/SANT genes per assembly. Pearson coefficients were calculated for *Aspergillus* and *Penicillium* species together and separately. The dotted red line is a straight line indicating a one to one ratio. (A, B, D) P-values for the differences between groups were calculated using Wilcoxon rank-sum tests with Bonferroni adjusted p-values for multiple tests (* = p <= 0.05; ** = p <= 0.01; *** = p <= 0.001; **** = p <= 0.0001). Non-significant differences are not labelled.

**Table 1.** *Starship*-related genes used for identifying *Starship-like* regions and their identifying features. Identifying features come from annotation methods and are a combination of InterPro protein domains, InterPro protein families and EggNog categories.

| Starship-related gene | Annotation features |
| --- | --- |
| DUF3435/Captain | IPR021842 (with no other IPRs)<br>IPR021842 + IPR013087 + IPRXXXXXX<br>IPR021842 + IPR011010 (with no other IPRs)<br>IPR021842 + IPR011010 + IPR013762 (with no other IPRs)<br>IPR021842 + IPR013087 + IPR011010 + IPR013762 (with no other IPRs) |
| MYB/SANT | IPR001005 (with no other IPRs)<br>IPR001005 + IPR009057 (with no other IPRs)<br>IPR001005 + IPR009057 + IPR013087 (with no other IPRs) |

| *Starship*-related gene | Annotation features |
| --- | --- |
| DUF3723 | IPR022198 (with no other IPRs) |
| PLP | ENOG503NY7U |
| CRG | ENOG503P5Q6 |
| DUF6066 | PF20255 |
| DUF3435/MYB hybrid | IPR021842 + IPR001005 (with no other IPRs) |

Additionally, we have many examples of *Starships* found within contigs, therefore more likely to represent complete *Starship* elements, and/or have been transferred between distant genomes that are clearly bordered by a Captain at one edge and the MYB/SANT gene at the other ( **Fig. 6C**; **fig. S23**; **fig. S25**; **fig. S26**; **fig. S33**; **fig. S36**; **fig. S37**). The six given examples all contained Terminal Inverted Repeats flanking Captain and MYB elements. This configuration was also detected in four of the previously identified complete *Starships* from Gluck-Thaler et al. 2022 (**fig. S37**). Notably, we also found genes that contained both the DUF3435 and MYB/SANT protein domains, including in the previously annotated *Starship* (Starship-Aory1) (Gluck-Thaler et al. 2022) (**fig. S37**). Using the set of non-redundant *Starship-like* regions, we also compared their size separated by whether they contained both Captain and MYB/SANT genes together, either one of the genes without the other (e.g. contains a Captain but no MYB/SANT gene), or neither. We see that *Starship-like* regions containing both genes were significantly larger than those containing only the Captain or neither (p <= 0.00012; **Fig. 6D**). No significant difference in size was observed with *Starship-like* regions containing only the MYB/SANT gene although the size is on average smaller. This smaller average size could be accounted for by both assembly breaks that therefore place the Captain and MYB/SANT on separate contigs, or structural variation which would separate them physically within the genomes. Lastly, the *Starship-like* regions containing only the MYB/SANT gene are on average larger than those with only the Captain. Considering both Captain-and MYB/SANT-only regions would be subject to the same impact of assembly breaks and structural variations, this suggests that the MYB/SANT-containing regions initially come from on average larger *Starships*. Taken together, these results indicate that there is an association between larger *Starships* and the presence of MYB/SANT genes.

Lastly, we constructed a MYB/SANT database using all genes identified as MYB/SANT *Starship*-related genes and used this to replace the Captain database in *Starfish*. *Starfish* was then run on the same long and short-read genome datasets for *Penicillium* and *Aspergillus*. The MYB/SANT count recapitulated much of the same differences when comparing clades and we see a strong correlation between MYB/SANT count and Captain count, where the number of Captains is on average slightly more than MYB/SANT genes (**Fig. 6E**).

## Discussion

There is now growing evidence that *Starships* are widespread in a subphylum of filamentous fungi, namely the Pezizomycotina (Bucknell and McDonald 2023; Gluck-Thaler and Vogan 2024; A. Urquhart et al. 2024; A.S. Urquhart et al. 2024), horizontally transferring between individuals. The Captain (DUF3435), a gene encoding a tyrosine site-specific recombinase, has been shown to be the only gene required for the excision and reintegration of the entire *Starships* (Urquhart et al. 2023); it has also been shown that *Starships* are not duplicated in genomes (cut and paste model) and that genes present in *Starships* are very rarely duplicated in genomes (Gluck-Thaler et al. 2022). In our dataset, we also confirmed these previous findings, i.e. no multiple copies of same *Starships* but we found duplicated cargo genes in some cheese strains within different *Starships*. When analysing some specific *Starships* with long-read assemblies, we found that i) the same *Starships* found in different species were located at non-homologous loci (**fig. S20**; **fig. S22**; **fig. S23**; **fig. S24**), ii) the presence of *Starships* was not following the phylogeny and iii) all strains of a given species did not carry the same *Starships,* confirming *Starships* are horizontally transferred. Much of what we have found and discussed here further elevates the idea of the recently proposed *Starship* compartment (A. Urquhart et al. 2024) as a pervasive region within genomes with distinct evolutionary trajectories that in turn influence the evolution of their hosts.

### Identifying the other edge of large *Starships*

The only structurally conserved gene within *Starships* described so far is the putative tyrosine recombinase, also referred as Captain/DUF3435, positioned close to one edge of *Starships* (Gluck-Thaler et al. 2022; Urquhart et al. 2023). Here we show that a MYB/SANT transcription factor is present in a subset of *Starships* and structurally positioned at the opposite edge to the Captain. This combination of genes, a transposase and MYB/SANT gene, can also be found in Harbinger transposons, both of which are required for transposition (Sinzelle et al. 2008); Harbinger DNA transposons have been identified in diverse phyla of life including protists, plants, insects and vertebrates. Considering this similarity, we could hypothesise that the MYB/SANT gene in *Starships* plays a similar role, aiding in both the nuclear import and TIR recruitment of the transposase. However, MYB/SANT do not appear required for *Starships*, as they are often absent from seemingly complete *Starships* and are found at a lower frequency than Captains. Therefore, MYB/SANT presence may be only beneficial for transposition, particularly in larger *Starships*, as our data suggests. Alternatively, the MYB/SANT TF may impact the movement of other *Starships* also present within the same genome. Much like previous HGT regions retroactively described as *Starships*, we found MYB/SANT genes in previously described *Starships* confirming their presence regardless of the pipeline used to discover the region.

### *Starships*, a convergent mechanism for food and clinical filamentous fungi to rapidly adapt to their environments

Using both inter-and intra-species comparisons, of varying phylogenetic distances with seven independent events of domestication for food, our findings reveal that all species/populations linked to food production in both the *Aspergillus* and *Penicillium* genera exhibit a distinct increase in the number of *Starships* compared to their environmentally isolated counterparts (**Fig. 1**; **Fig. 2**; **Fig. 3**; **Fig. 4**; **table S1**) within seven independent instances of species domesticated for food production, including cheeses, dry-cured meats, and various soy and rice-based foods. Furthermore, many genes within *Starships* have putative functions relevant for adaptation to food environments, such as genes involved in lactose metabolism and salt tolerance. We found, building upon previous observations (Cheeseman et al. 2014; Ropars et al. 2015; Lo et al. 2023), that *Starships* and their food relevant cargo are commonly shared among food-related species and in turn highlighting *Starships* as a generalisable mechanism for these filamentous fungi to rapidly adapt to these new niches.

Fermented food environments are rich in terms of both nutrients and microbes (number and species diversity), including bacteria, yeasts and filamentous fungi, making them ideal media for genetic exchange (Wolfe and Dutton 2015). For example, several *Penicillium* and *Aspergillus* species have been described to co-occur in cheeses alongside many other fungi (Martin and Cotter 2023). These exchanges between fungi co-occuring in the cheese mimics what has been described in cheese-related bacteria, as HGT events appear to be common, contain genes with functional relevance for cheese adaptation and in some cases facilitated by ICEs (Bonham et al. 2017). Similar to ICEs, due to a *Starship*’s relatively large size, these transfer events are certainly costly and should be maintained under strong selection. This is particularly relevant in some cases such as in the cheese-related species *P. camemberti var. caseifulvum,* where we identified that greater than 15% of the ESE00019 genome is made up of *Starship*-related material. Furthermore, considering that *Starships* have been transferred between species from the same food-related environments (Cheeseman et al. 2014; Ropars et al. 2015; Lo et al. 2023), that they can be very numerous in a single host genome and that most *Starships* identified are unique, even if at times containing similar cargo, this establishes their crucial role in adaptation and their ease of transfer; extrapolated, additional cargo may yet be acquired in domesticated strains. However, we still don’t know how often *Starships* move or whether they can be lost as easily as they are acquired.

Although food environments could encourage the likelihood of cross species *Starship* transfers (assuming they only require physical contact), this may only be part of the story, particularly due to elevated *Starship* content in clinical strains. A recent and thorough investigation into *Starships* in *A. fumigatus* showed that they not only promote strain heterogeneity but also that certain *Starships* are enriched in clinical strains (Gluck-Thaler et al. 2024). The current prevailing theory is that clinical strains of *Aspergillus*, with pathogenic adaptations, have acquired these prior to colonisation (Rhodes et al. 2022). The primary example in *A. fumigatus,* azole resistance, is said to develop in agricultural settings due to the use of azole based antifungal treatments. This would therefore likely be the same pressure driving the increase in *Starship* content, and considering that most environmental strains of *A. fumigatus* came from agricultural settings, may explain why there is a smaller difference between clinical and environmental strains, compared to other food-related analyses.

### *Starships,* a more general mechanism to rapidly adapt and survive to extreme environments?

Beyond food and clinical isolates, we were also able to identify a relative increase in *Starships* in several phylogenetically distant fungi occurring in saline environments, unrelated to food. However, food environments linked to *Starships* are almost all salted, i.e. cheese, cured meat, soy sauce, miso, suggesting salt could play a part in *Starship* movement. A variety of stresses, including salt, have already been identified as a means of inducing the transfer of plasmids and TE transposition (Beuls et al. 2012). In this scheme, certain conditions such as high salinity might not only select for *Starships* that confer resistance but may also increase general *Starship* mobility and subsequent accumulation. However, we also identified potentially elevated *Starships* in strains isolated from heavy metal contaminated soil (**fig. S38**) aligning with the previously described *Starship* acquired heavy metal resistance (Urquhart et al. 2022; Urquhart et al. 2023). Furthermore, the ethanol utilisation cluster was found primarily in species known to cause post-harvest disease in fruits. Ethanol resistance and utilisation may be in response to early colonising yeast that can rapidly produce volatile ethanol in order to deter competition (Freimoser et al. 2019).

### Cargo-clusters drive the functional convergence of *Starships*

Our data on *Starship*-related gene clusters highlights that ‘cargo-clusters’ frequently move among different *Starships* as single elements. This is supported by the recently published results of Urquhart et al. 2023, where they describe the same cargo-cluster, responsible for resistance to formaldehyde, in several different *Starships*, referred to as cargo-swapping (A.S. Urquhart et al. 2024). Within the formaldehyde cluster, it was experimentally shown that ssfB granted much greater formaldehyde resistance compared to the six other genes and that it was the only gene consistently maintained within the cluster, whilst the other genes’ presence varied (A.S. Urquhart et al. 2024). The arsenic cluster in the *Starship Hephaestus* has similarly been shown to have been acquired as a single element including a truncated flanking gene (Urquhart et al. 2022). Here we showed several cargo-clusters, unlike those containing biosynthetic pathways, that contain variable gene content with similarly functional roles. For example, the arsenic cluster, previously described in the *Starship Hephaestus*, contains a set of three core genes (ArsC, ARR3 and ArsH) alongside seven accessory cargo-cluster genes (**fig. S32**). This variable arrangement is similar to that of arsenic gene islands in prokaryotes (Ben Fekih et al. 2018). It is therefore possible that cargo-clusters may only orient themselves around certain more functionally important genes, providing a more malleable adaptive structure whilst conserving the clusters’ function. This is particularly likely considering that cargo is in itself accessory material.

In conclusion, *Starships* thus appear to be a way of rapidly adapting and surviving to novel conditions, depending on the cargo genes acquired, having been employed heavily within human-related environments. By focusing on recent evolutionary events, domestication and pathogenesis, we were able to clearly detect more *Starships* in food and clinical isolates compared to their environmental counterparts. Furthermore, we have described numerous cargo genes involved in resistance to salt, sulfites and even drugs. Many of these are antimicrobial compounds employed by humans for food preservation or against pathogens of both humans and plants. This may become important as fungal pathogens are becoming increasingly resistant to antifungal treatments (Yin et al. 2023) and their ability to rapidly acquire and transfer resistance would certainly worsen treatment outcomes. Our findings have thus clear implications for the future of agriculture, human health and the food industry as we provided a comprehensive overview on a widely recurrent mechanism of gene transfer between filamentous fungi by focusing on two genera. This also indicates a need for caution regarding the use of bioengineered fungal strains in our agriculture or food systems, as already proposed in the fermented food species *Aspergillus oryzae* which was successfully genetically modified using the CRISPR-Cas9 system (Maini Rekdal et al. 2024). We cannot predict how these genes will be spread and whether they can be future threats.

## Methods

### Sampling

We newly isolated 68 strains from cheese rinds. To do so, we performed successive dilutions of pieces of rinds to obtain isolated spores on solid malt extract agar. We checked the species identity using the beta-tubulin locus using bt2a and bt2a primers (seq primers). **Table S1** recapitulates strain details, including species identification, origin and country of origin.

### DNA extraction, genome sequencing

We used the Nucleospin Soil Kit (Macherey-Nagel, Düren, Germany) for DNA extraction of the 68 newly isolated strains grown for five days on malt extract agar. DNA was sequenced using Illumina technologies (paired-end, 100bp and 150bp) (**table S1**).

### Illumina genome assembly

We assembled the genomes of the 68 newly isolated strains and of the 49 already published strains (Ropars et al 2020; **table S1**). Illumina reads were checked with FastQC (v0.11.9) (Andrews 2020). Leading or trailing low quality or N bases below a quality score of three were removed. For each read, only parts that had an average quality score higher than 20 on a four base window were kept, and only reads with at least a length of 36 bp were kept. Cleaned reads were assembled with SPAdes (v3.15.3) (Prjibelski et al. 2020) not using unpaired reads using the--careful parameter.

### Long-read genome assembly

We newly assembled 3 genomes (ESE00019, ESE00090 and LCP05531) with Oxford Nanopore Technology (ONT), newly assembled 1 genomes (LCP05904) with Pacbio and re-assembled 3 genomes (LCP06093, PN007, PN016) with publicly available Pacbio data. We assembled each dataset with a variety of assemblers and with different preprocessed read datasets and settled on two assemblies depending on the technology. Additionally, illumina data was available for all ONT sequenced strains, and one re-assembled strain, PN016. For ONT sequenced strains, data was basecalled using Guppy (v5; model r941_min_fast_g507) and assembled using Canu (v2.2) (Koren et al. 2017) with all raw reads and Flye (v2.9-b1768) (Kolmogorov et al. 2019) after using Filtlong (v0.2.1) (github.com/rrwick/Filtlong) to remove all reads less than 5kb. These assemblies were merged using Ragtag’s (v2.1.0) (Alonge et al. 2022) *patch* function with the Canu assembly as the ‘target’ and the flye assembly as the ‘query’. A second round of merging was done replacing the flye assembly with the merged assembly from the first round. This merged assembly was then polished with Medaka (v1.6.0) (github.com/nanoporetech/medaka) using all the ONT reads. The assembly was then polished using the illumina reads with Hapo-G (v1.3.2) (Aury and Istace 2021). For Pacbio based genomes, both the Canu and Flye assembly was assembled using the Filtlong based reads, removing all reads shorter than 5kb and merging was performed using Quickmerge (v0.3) (Chakraborty et al. 2016) with the flye assembly as the ‘reference’ and the Canu assembly as the ‘query’. One round of Hapo-G polishing was performed for PN017. All assemblies were then treated to remove redundant contigs. Contig alignment was performed by BLAST (Camacho et al. 2009) and contigs were removed if both 99% or more was covered by another larger contig with 99% or more identity. For the contig naming; contigs were organised by largest to smallest then numbered and mitochondrial contigs were identified by BLASTing all contigs against a manually identified complete mitochondrial genome from ESE00019.

We assessed genome completeness using the BUSCO (v5.3.1) eurotiales_odb10 database (Manni et al. 2021) and Merqury (-k 18) (Rhie et al. 2020) for those with illumina data (**table S4**). We also compared genomes using assembly statistics (**table S4**). Additionally we compared our long read assemblies to short-read assemblies if available. This shows that long-read assembly and reassembling of these genomes resulted in significant improvements in contiguity, BUSCO and Merqury completeness compared to previous assemblies both long and short.

### Long-read genome annotation

All genomes excluding PN016 and PN007 were annotated. Genomes were first analysed by RepeatModeler (v2.0.2) (Flynn et al. 2020) using the *BuildDatabase* and *RepeatModeler* functions (-engine ncbi). Repeat families were further clustered using cd-hit-est (v4.8.1) (-d 0 -aS 0.8 -c 0.8 -G 0 -g 1 -b 500) (Li and Godzik 2006). This repeat library was then fed to RepeatMasker (v4.1.2-p1) (http://www.repeatmasker.org) to identify and mask all repeats (-xsmall -s). The masked assembly was then annotated by Braker2 (v2.1.6) (--fungus --gff3) (Brůna et al. 2021) using a set of all BUSCOs in fungal taxonomic groups (Manni et al. 2021). Braker was then rerun using the previous run’s augustus protein hints for 4 rounds. The resulting protein dataset was then analysed using the initial BUSCO database, showing 99.9% of all proteins were annotated for ONT based assemblies and 94.7-94.6% for Pacbio based assemblies. The resulting proteins were annotated using Funannotate (v1.8.11) (Palmer and Stajich 2020). Firstly a funannotate database was generated using the *setup* command (-b fungi). We then used the funannotate’s *iprscan* wrapper (Jones et al. 2014) and emapper (v2.1.8) (Cantalapiedra et al. 2021) to annotate protein domains and Eggnog functions respectively. The masked genome, the braker gff3, the funannotate fungi database, the iprscan output and the eggnog annotations were then combined using butanoate’s *annotate* function.

### Phylogenomics

We placed all genomes used in our study within a phylogenomic framework, separated into the *Penicillium* and *Aspergillus* genera, using several genomes of the other genus for rooting (**table S1**). This served several purposes, such as verifying species identification for public assemblies (which were frequently incorrect and have been tagged in **table S1**), confirming species relationships including their distances and highlighting complex species naming systems such as in *Aspergillus* species complexes. This genome dataset excluded any NCBI genomes labelled as abnormal or from metagenomic data. All genomes were analysed by BUSCO using the eurotiales_odb10 database (Manni et al. 2021). Any genome with less than 80% complete single copy BUSCOs was removed from further analysis. We then established a protein dataset consisting of only BUSCOs contained within 99% of all genomes (3154/4190 and 3258/4190 for *Penicillium* and *Aspergillus* respectively) which were then used to build each tree. Using BUSCOs allowed us to heavily limit any impact of horizontally transferred genes on the topography. The BUSCO datasets were aligned by mafft (v7.310) (Katoh and Standley 2013) and trimmed by trimal (v1.4.rev15) (Capella-Gutiérrez et al. 2009). All protein alignments were concatenated using *catfasta2phyml/catfasta2phyml.pl* (https://github.com/nylander/catfasta2phyml) (-c -f) to produce a single fasta file. We then used this to generate a species tree with VeryFastTree (-double-precision) (v4.0.3) with the default SH-like node support calculations (**Supp. files 1** and **2**) (Piñeiro et al. 2020). The resulting treefile was analysed in R (R Core Team 2024). Each tree was rooted using the outgroup genomes with treeio::MRCA (v1.22) (Wang et al. 2020), ape::root.phylo (v5.7.1) (Paradis and Schliep 2019), TreeTools::Preorder (v1.10.0) (Smith 2019) and treeio::droptip and visualised using the ggtree suite (Yu et al. 2017). Renaming of species was performed, based on the phylogeny as noted in **table S1**, with most being supported by other publications, also noted in **table S1**. It should be kept in mind that renaming is specific to the genome, regardless of the strain name assigned in NCBI. Renaming efforts were mainly targeted towards species with numerous genomes and we do not want to enter the debate surrounding the naming of *A. terreus*/*A. pseudoterreus*. For phylogenies with collapsed clades (e.g **Fig. 2** and **Fig. 3**) we used *scaleClade* (scale=0.1) then *collapse* (mode=‘mixed’) from the ggtree suite. Therefore these species or clade specific nodes display a triangle a tenth the size of the actual branches and with corners extending to both the maximum and minimum of these scaled branch lengths. The phylogenetics pipeline can be found at https://github.com/SAMtoBAM/publicgenomes-to-buscophylogeny.

### Strain isolation origins and classification

For each genome assembly we attempted to manually gather information on the strains’ isolation origins. Classifying these origins into simple categories was then also done manually. For environmental origins we used several environmental categories, i.e. environmental, environment-clinical, environment-food and environment (saline). Environment (saline) was used for Aspergillus strains where the environment would be generally considered as extreme in its saline content such as sea-water or saline soils. Environment-food was used primarily for contamination of processed food-products such as unwanted growth on dairy products. Environment-clinical strains mainly come from a single study which isolated fungi from healthy human faecal samples. Other strains in this latter category were isolated from hospital and/or human samples but unassociated with disease. **Table S1** contains the detailed and simplified isolation origins.

### Starfish identification on all (short-and long-read) genomes

In order to estimate the starship count within and between species, we used Starfish to identify DUF3435/Captain genes in both long and short-read assemblies. The *Captain* count was then used as a proxy for the number of *Starships*. We ran *Starfish annotate* with default settings and the default Captain database (YRsuperfamRefs.faa). The number of captains and the isolation origins per genome was then plotted against the phylogeny in R. To reorder plots of Captain counts against phylogenetic trees, bar graphs were made by ggplot (Wickham 2016) then combined with ggtree (v3.6.2) (Yu et al. 2017) phylogenies using the R library aplot (Yu 2023).

To identify changes in *Starship* content, species known to have been domesticated for food production or cause disease were compared against either wild sister species, or within species split by isolation origins. In all cases, to test for differences in Captain counts between species and/or isolation environments, we performed Wilcoxon rank-sum tests with Bonferroni adjusted p-values for multiple comparisons.

We tested the accuracy of this proxy by looking for positive correlations between Captain count and genome size within species/species-complexes in which a large number of genomes were available. In all cases we detected strong and significant positive relationships (**fig. S1**; **fig. S2**) establishing that we are likely correct in equating Captain count with *Starship* insertions.

To test the accuracy of Captain count as a proxy, alongside the captains specificity within *Starships*, genome sizes were compared against the number of captains using the Pearson correlation coefficient in R. In **Fig. 1** two strains of *P. janthinellum* were removed due to an extremely high Captain count which was most likely erroneous due to no corresponding increase in genome size.

In *A.fumigatus*, in order to remove any bias in how genomes were assembled across projects, we restricted the isolation-*Starship* comparison to a large subset of 252/336 genomes, containing both environmental (202/252) and clinical (50/252) isolates, that all originated from two bioprojects/studies (PRJNA697844 and PRJNA595552) from the same group and therefore were all assembled and treated with the same pipeline. In doing so we still see, as shown with all genomes, that clinical strains contain more starships (p=0.034; **fig. S29**).

### Pairwise captain identity and Average Amino Acid Identity

To also test the assumption that captain occurrence is associated with HGT we tested the similarity between captains across all species. We concatenated all captain proteins identified by Starfish and constructed a Diamond database (Buchfink et al. 2021). We then searched this database using the same entire set of captain proteins, all vs all, using *diamond blastp*, keeping only matches where both the reference and query protein were covered >=95% and an amino acid identity >=95% (--very-sensitive --id 95 --query-cover 95 --subject-cover 95 --no-self-hits). This gave us a set of highly similar captain proteins based on length and composition.

For all pairs of genomes identified by captain similarity we calculated the average amino acid identity (AAI) using 50 random BUSCOs, all of which were used to build both the Penicillium and Aspergillus phylogenies. We analysed the BUSCO dataset in the same fashion as the Captains, using Diamond, but with only a filter for 80% reference and query coverage (--very-sensitive --query-cover 80 --subject-cover 80 --no-self-hits).

Due to some species having many assemblies, and in order to focus on identifying likely examples of HGT, we reduced the dataset to a non-redundant set. That is, we filtered for unique pairs of captains in terms of both the captain length and identity, the AAI and the two species involved. Following this we examined examples of high discrepancy between pairwise Captain identity and their respective genomes AAI, i.e. where Captains from genomes were much more similar than the BUSCOs. To examine these cases we aligned the entire genomes using nucmer, extracting all regions that aligned >7.5kb, taking 50kb up and downstream of these alignments and visualising the loci alongside the captain position using gggenomes (fig. S20-28).

### Genome graph

Using NCBI (Sayers et al. 2022) we identified publicly available, contiguous (relatively high contig N50), long-read technology assemblies within species of interest and combined them with our newly assembled genomes (**table S2**). We separated them into groups based on their relatedness and generated genome graphs for each using pggb (v0.5.0-hdfd78af) (-p 75 -s 30000 -m -S -k 19-G 7919,8069 -Y _) (Garrison et al. 2023). We also set the option -n as the number of genomes used per grouping. The maximum distance (-p) was only reduced to 50% (-p 50) for the Brevicompacta graph to help with the larger distances between *P. brevicompactum* and *P. salamii*. The script for generating a genome-graph and detecting *Starship-*like regions can be found here https://github.com/SAMtoBAM/pggb_starship_pipeline.

### *Starship*-like region detection

To extract presence and absence data we used odgi (v0.8.1-1-ge91b1cd) (Guarracino et al. 2022) *paths*, extracted both the paths and associated genomes, then fed this into odgi *pav* alongside 1kb windows for each genome (generated using bedtools (v2.26.0) (Quinlan and Hall 2010) *makewindows*). We then extracted all regions absent from at least one genome, merged these windows within 1bp using bedtools *merge*, removed all remaining regions smaller than 10kb then remerged regions with a max gap of 20 kb. The final resulting regions were then filtered for a minimum size of 30kb. This dataset was then annotated with braker, emapper and funannotate, as with the whole genomes, except using only a single round of braker annotation.

Following this the resulting annotation files were searched for protein domains and Eggnogs indicative of *Starship-like* genes. Table 1 below shows the list of genes and their respective tags used to find them. These *Starship*-related genes are made up of previously identified components of starships from Gluck-Thaler et al. 2022 (DUF3435/Captain, DUF3723, PLPs) alongside newly added conidiophore related genes (CRGs), DUF6066 and the MYB/SANT. These new genes alongside the previously identified genes were shown to be highly starship specific in all genomes analysed.

The *Starship*-related gene annotation and identification pipeline was initially tested on the *Starships* released in Gluck-Thaler et al. 2022 showing we were able to effectively identify the *Captains* in 33/39 (85%) of these *Starships* and additionally MYB/SANT genes at the distal end of 4/39 (**fig. S27**).

Additionally *Starfish* (Gluck-Thaler and Vogan 2024) was used on each dataset similarly in order to detect DUF3435/Captain genes using the default database (YRsuperfamRefs.faa) and default settings. Additional *Captains*, that were not already identified, were added to the set of *Starship*-related genes within the defined candidate regions. Notably this added an additional 338/986 Captains (34%). Any region containing a *Starship*-related gene was then considered a *Starship-like* region (**Supp. file 3**).

*Starship-*like regions were reduced to a non-redundant set using a typical repeat clustering method of 85% identity and 85% coverage using BLAST, taking the largest SLR in each cluster. The SLRs in each cluster were then aligned against one another and a genome identified as missing the SLR using nucmer from MUMMER4. The aligned regions were then visualised using R and gggenomes.

### Functional enrichment

For functional enrichment we used COG gene annotations all from the same annotation pipeline. Genes were classified by their COG category or given ‘Absent’ in the absence of annotation. Our first dataset contained our four newly assembled and annotated long-read assemblies. This contains three strains from cheese-related species (ESE00019, LCP06093, LCP05531) and one strain from an environmental species (ESE00090). For each genome we separated genes into those present or absent from the *Starship*-like regions detected by the genome-graph. For each genome, all genes were annotated simultaneously, therefore we limit bias in annotation pipelines or runs that could explain differences in cargo and non-cargo genes. Hypergeometric tests for enrichment and depletion were performed for each COG category and the p-value was adjusted for multiple comparisons (Benjamini–Hochberg) using phyper and p.adjust respectively (R Core Team 2024). We also performed an analysis on the combined cargo and non-cargo genes from all three cheese-related strains, which showed the same differences. Second, we performed the same tests on the COG categories of cargo in *Starship*-like regions, split into groups based on the strains isolation category, either food, clinical or environment. The above tests were all performed with and without the COG categories ‘S’, ‘L’, ‘K’ and ‘Absent’. S and Absent are unknown categories. L and K are commonly associated with Transposable elements and our Starship-like regions were not masked prior to annotation.

### Clinker

A gff3 file for each *Starship-like* region, containing a gene cluster, was converted to an EMBL file using EMBLmyGFF3 (v2.3) (Norling et al. 2018). The EMBL file was then converted to a genbank file using biopython (*SeqIO.convert(sys.stdin, ’embl’, sys.stdout, ’genbank’)*) (Cock et al. 2009). This two step process avoided any formatting issues given by other tools that converted gff3 files directly to gbk. We then ran Clinker (v0.0.28) (-i 0.5), with a minimum of 50% sequence alignment identity, on all the genbank files simultaneously for each gene cluster. The html output was manually modified in order to highlight the clusters and shorten the length of regions. The Saccharomyces Genome Database (Engel et al. 2022), Candida genome database (Skrzypek et al. 2017), fungidb (Basenko et al. 2018) and NCBI (Sayers et al. 2022) were indispensable in manually investigating the function and relevance of these genes.

### Cargo-clusters in all assemblies

All assemblies used in this study (Table S1) were combined within a single BLAST database and queried using the core genes for each cluster, with a minimum query coverage and identity of 95%. A cluster was considered present if all core genes contained a match. The presence/absence plot (Fig 6) was generated using ggplot and geom_tile (Wickham 2016) then combined with ggtree (v3.6.2) (Yu et al. 2017) phylogenies using the R library aplot (Yu 2023).

**S. O’Donnell**: Conceptualization, Data Curation, Formal Analysis, Investigation, Methodology, Software, Validation, Visualisation, Writing-original draft, Writing-review and editing. **G. Rezende**: Investigation. **J.-P. Vernadet**: Data Curation, Investigation, Software. **A. Snirc**: Investigation. **J. Ropars**: Conceptualization, Data Curation, Funding acquisition, Investigation, Methodology, Project administration, Resources, Supervision, Validation, Writing-original draft, Writing-review and editing.

## Acknowledgements

We would like to thank Emile Gluck-Thaler for his insights and suggestions on the manuscript.

## References

Acevedo KL, Eaton E, Zhao S, Chacon-Vargas K, McCarthy CM, Choi D, Yu J-H, Gibbons JG. 2023. Population genomics of Aspergillus sojae is shaped by the food environment.:2023.09.07.556736. Available from: https://www.biorxiv.org/content/10.1101/2023.09.07.556736v1

Alonge M, Lebeigle L, Kirsche M, Jenike K, Ou S, Aganezov S, Wang X, Lippman ZB, Schatz MC, Soyk S. 2022. Automated assembly scaffolding using RagTag elevates a new tomato system for high-throughput genome editing. Genome Biol. 23:258.

Andrews S. 2020. FastQC: a quality control tool for high throughput sequence data. Available online at: http://www.bioinformatics.babraham.ac.uk/projects/fastqc. Available from: https://www.bioinformatics.babraham.ac.uk/projects/fastqc/

Antonucci I, Gallo G, Limauro D, Contursi P, Ribeiro AL, Blesa A, Berenguer J, Bartolucci S, Fiorentino G. 2017. An ArsR/SmtB family member regulates arsenic resistance genes unusually arranged in Thermus thermophilus HB27. Microb. Biotechnol. 10:1690–1701.

Arai N, Sekizuka T, Tamamura Y, Kusumoto M, Hinenoya A, Yamasaki S, Iwata T, Watanabe-Yanai A, Kuroda M, Akiba M. 2019. Salmonella Genomic Island 3 Is an Integrative and Conjugative Element and Contributes to Copper and Arsenic Tolerance of Salmonella enterica. Antimicrob. Agents Chemother. 63:10.1128/aac.00429-19.

Ariño J, Ramos J, Sychrová H. 2010. Alkali Metal Cation Transport and Homeostasis in Yeasts. Microbiol. Mol. Biol. Rev. 74:95–120.

Aury J-M, Istace B. 2021. Hapo-G, haplotype-aware polishing of genome assemblies with accurate reads. NAR Genomics Bioinforma. 3:lqab034.

Barrs VR, van Doorn TM, Houbraken J, Kidd SE, Martin P, Pinheiro MD, Richardson M, Varga J, Samson RA. 2013. Aspergillus felis sp. nov., an Emerging Agent of Invasive Aspergillosis in Humans, Cats, and Dogs. PLoS ONE 8:e64871.

Basenko EY, Pulman JA, Shanmugasundram A, Harb OS, Crouch K, Starns D, Warrenfeltz S, Aurrecoechea C, Stoeckert CJ, Kissinger JC, et al. 2018. FungiDB: An Integrated Bioinformatic Resource for Fungi and Oomycetes. J. Fungi 4:39.

Ben Fekih I, Zhang C, Li YP, Zhao Y, Alwathnani HA, Saquib Q, Rensing C, Cervantes C. 2018. Distribution of Arsenic Resistance Genes in Prokaryotes. Front. Microbiol. 9:2473.

Beuls E, Modrie P, Deserranno C, Mahillon J. 2012. High-Salt Stress Conditions Increase the pAW63 Transfer Frequency in Bacillus thuringiensis. Appl. Environ. Microbiol. 78:7128– 7131.

Bongomin F, Gago S, Oladele RO, Denning DW. 2017. Global and Multi-National Prevalence of Fungal Diseases—Estimate Precision. J. Fungi 3:57.

Bonham KS, Wolfe BE, Dutton RJ. 2017. Extensive horizontal gene transfer in cheese-associated bacteria. Handley K, editor. eLife 6:e22144.

Briza P, Eckerstorfer M, Breitenbach M. 1994. The sporulation-specific enzymes encoded by the DIT1 and DIT2 genes catalyze a two-step reaction leading to a soluble LL-dityrosine-containing precursor of the yeast spore wall. Proc. Natl. Acad. Sci. U. S. A. 91:4524–4528.

Brůna T, Hoff KJ, Lomsadze A, Stanke M, Borodovsky M. 2021. BRAKER2: automatic eukaryotic genome annotation with GeneMark-EP+ and AUGUSTUS supported by a protein database. NAR Genomics Bioinforma. 3:lqaa108.

Buchfink B, Reuter K, Drost H-G. 2021. Sensitive protein alignments at tree-of-life scale using DIAMOND. Nat. Methods 18:366–368.

Bucknell A, Wilson HM, Santos KCG do, Simpfendorfer S, Milgate A, Germain H, Solomon PS, Bentham A, McDonald MC. 2024. Sanctuary: A Starship transposon facilitating the movement of the virulence factor ToxA in fungal wheat pathogens.: 2024.03.04.583430. Available from: https://www.biorxiv.org/content/10.1101/2024.03.04.583430v1

Bucknell AH, McDonald MC. 2023. That’s no moon, it’s a Starship: Giant transposons driving fungal horizontal gene transfer. Mol. Microbiol.

Camacho C, Coulouris G, Avagyan V, Ma N, Papadopoulos J, Bealer K, Madden TL. 2009. BLAST+: architecture and applications. BMC Bioinformatics 10:421.

Cantalapiedra CP, Hernández-Plaza A, Letunic I, Bork P, Huerta-Cepas J. 2021. eggNOG-mapper v2: Functional Annotation, Orthology Assignments, and Domain Prediction at the Metagenomic Scale. Mol. Biol. Evol. 38:5825–5829.

Capella-Gutiérrez S, Silla-Martínez JM, Gabaldón T. 2009. trimAl: a tool for automated alignment trimming in large-scale phylogenetic analyses. Bioinformatics 25:1972–1973.

Caracuel Z, Casanova C, Roncero MIG, Di Pietro A, Ramos J. 2003. pH Response Transcription Factor PacC Controls Salt Stress Tolerance and Expression of the P-Type Na+-ATPase Ena1 in Fusarium oxysporum. Eukaryot. Cell 2:1246–1252.

Chakraborty M, Baldwin-Brown JG, Long AD, Emerson JJ. 2016. Contiguous and accurate de novo assembly of metazoan genomes with modest long read coverage. Nucleic Acids Res. 44:e147.

Cheeseman K, Ropars J, Renault P, Dupont J, Gouzy J, Branca A, Abraham A-L, Ceppi M, Conseiller E, Debuchy R, et al. 2014. Multiple recent horizontal transfers of a large genomic region in cheese making fungi. Nat. Commun. 5:2876.

Cock PJA, Antao T, Chang JT, Chapman BA, Cox CJ, Dalke A, Friedberg I, Hamelryck T, Kauff F, Wilczynski B, et al. 2009. Biopython: freely available Python tools for computational molecular biology and bioinformatics. Bioinformatics 25:1422–1423.

Crequer E, Ropars J, Jany J-L, Caron T, Coton M, Snirc A, Vernadet J-P, Branca A, Giraud T, Coton E. 2023. A new cheese population in Penicillium roqueforti and adaptation of the five populations to their ecological niche. Evol. Appl. 16:1438–1457.

Denning DW. 2024. Global incidence and mortality of severe fungal disease. Lancet Infect. Dis. [Internet]. Available from: https://www.sciencedirect.com/science/article/pii/S1473309923006928

Dumas E, Feurtey A, Rodríguez de la Vega RC, Le Prieur S, Snirc A, Coton M, Thierry A, Coton E, Le Piver M, Roueyre D, et al. 2020. Independent domestication events in the blue-cheese fungus Penicillium roqueforti. Mol. Ecol. 29:2639–2660.

Engel SR, Wong ED, Nash RS, Aleksander S, Alexander M, Douglass E, Karra K, Miyasato SR, Simison M, Skrzypek MS, et al. 2022. New data and collaborations at the Saccharomyces Genome Database: updated reference genome, alleles, and the Alliance of Genome Resources. Genetics 220:iyab224.

Felder T, Bogengruber E, Tenreiro S, Ellinger A, Sá-Correia I, Briza P. 2002. Dtr1p, a Multidrug Resistance Transporter of the Major Facilitator Superfamily, Plays an Essential Role in Spore Wall Maturation in Saccharomyces cerevisiae. Eukaryot. Cell 1:799–810.

Felenbok B, Flipphi M, Nikolaev I. 2001. Ethanol catabolism in Aspergillus nidulans: A model system for studying gene regulation. In: Progress in Nucleic Acid Research and Molecular Biology. Vol. 69. Academic Press. p. 149–204. Available from: https://www.sciencedirect.com/science/article/pii/S0079660301690470

Fernandez-Pittol M, Alejo-Cancho I, Rubio-García E, Cardozo C, Puerta-Alcalde P, Moreno-García E, Garcia-Pouton N, Garrido M, Villanueva M, Alastruey-Izquierdo A, et al. 2022. Aspergillosis by cryptic *Aspergillus* species: A case series and review of the literature. Rev. Iberoam. Micol. 39:44–49.

Flynn JM, Hubley R, Goubert C, Rosen J, Clark AG, Feschotte C, Smit AF. 2020. RepeatModeler2 for automated genomic discovery of transposable element families. Proc. Natl. Acad. Sci. 117:9451–9457.

Freimoser FM, Rueda-Mejia MP, Tilocca B, Migheli Q. 2019. Biocontrol yeasts: mechanisms and applications. World J. Microbiol. Biotechnol. 35:154.

Friesen TL, Stukenbrock EH, Liu Z, Meinhardt S, Ling H, Faris JD, Rasmussen JB, Solomon PS, McDonald BA, Oliver RP. 2006. Emergence of a new disease as a result of interspecific virulence gene transfer. Nat. Genet. 38:953–956.

Garrison E, Guarracino A, Heumos S, Villani F, Bao Z, Tattini L, Hagmann J, Vorbrugg S, Marco-Sola S, Kubica C, et al. 2023. Building pangenome graphs.: 2023.04.05.535718. Available from: https://www.biorxiv.org/content/10.1101/2023.04.05.535718v1

Gasmi N, Jacques P-E, Klimova N, Guo X, Ricciardi A, Robert F, Turcotte B. 2014. The Switch from Fermentation to Respiration in Saccharomyces cerevisiae Is Regulated by the Ert1 Transcriptional Activator/Repressor. Genetics 198:547–560.

Gibbons JG, Salichos L, Slot JC, Rinker DC, McGary KL, King JG, Klich MA, Tabb DL, McDonald WH, Rokas A. 2012. The Evolutionary Imprint of Domestication on Genome Variation and Function of the Filamentous Fungus Aspergillus oryzae. Curr. Biol. 22:1403–1409.

Gluck-Thaler E, Forsythe A, Puerner C, Stajich JE, Croll D, Cramer RA, Vogan AA. 2024. Giant transposons promote strain heterogeneity in a major fungal pathogen. bioRxiv:2024.06.28.601215.

Gluck-Thaler E, Ralston T, Konkel Z, Ocampos CG, Ganeshan VD, Dorrance AE, Niblack TL, Wood CW, Slot JC, Lopez-Nicora HD, et al. 2022. Giant Starship Elements Mobilize Accessory Genes in Fungal Genomes. Mol. Biol. Evol. 39:msac109.

Gluck-Thaler E, Vogan AA. 2024. Systematic identification of cargo-mobilizing genetic elements reveals new dimensions of eukaryotic diversity. Nucleic Acids Res.:gkae327.

Guarracino A, Heumos S, Nahnsen S, Prins P, Garrison E. 2022. ODGI: understanding pangenome graphs. Bioinformatics 38:3319–3326.

Iacumin L, Chiesa L, Boscolo D, Manzano M, Cantoni C, Orlic S, Comi G. 2009. Moulds and ochratoxin A on surfaces of artisanal and industrial dry sausages. Food Microbiol. 26:65– 70.

Jeffries TW, Van Vleet JRH. 2009. Pichia stipitis genomics, transcriptomics, and gene clusters. FEMS Yeast Res. 9:793–807.

Jones P, Binns D, Chang H-Y, Fraser M, Li W, McAnulla C, McWilliam H, Maslen J, Mitchell A, Nuka G, et al. 2014. InterProScan 5: genome-scale protein function classification. Bioinformatics 30:1236–1240.

Katoh K, Standley DM. 2013. MAFFT Multiple Sequence Alignment Software Version 7: Improvements in Performance and Usability. Mol. Biol. Evol. 30:772–780.

Keeling PJ. 2024. Horizontal gene transfer in eukaryotes: aligning theory with data. Nat. Rev. Genet.:1–15.

Kodedová M, Sychrová H. 2015. Changes in the Sterol Composition of the Plasma Membrane Affect Membrane Potential, Salt Tolerance and the Activity of Multidrug Resistance Pumps in Saccharomyces cerevisiae. PLoS ONE 10:e0139306.

Kolmogorov M, Yuan J, Lin Y, Pevzner PA. 2019. Assembly of long, error-prone reads using repeat graphs. Nat. Biotechnol. 37:540–546.

Koren S, Walenz BP, Berlin K, Miller JR, Bergman NH, Phillippy AM. 2017. Canu: scalable and accurate long-read assembly via adaptive k-mer weighting and repeat separation. Genome Res. 27:722–736.

Legras J-L, Galeote V, Bigey F, Camarasa C, Marsit S, Nidelet T, Sanchez I, Couloux A, Guy J, Franco-Duarte R, et al. 2018. Adaptation of S. cerevisiae to Fermented Food Environments Reveals Remarkable Genome Plasticity and the Footprints of Domestication. Mol. Biol. Evol. 35:1712–1727.

Li W, Godzik A. 2006. Cd-hit: a fast program for clustering and comparing large sets of protein or nucleotide sequences. Bioinformatics 22:1658–1659.

Lo Y-C, Bruxaux J, Rodríguez de la Vega RC, O’Donnell S, Snirc A, Coton M, Le Piver M, Le Prieur S, Roueyre D, Dupont J, et al. 2023. Domestication in dry-cured meat Penicillium fungi: Convergent specific phenotypes and horizontal gene transfers without strong genetic subdivision. Evol. Appl. 16:1637–1660.

Maini Rekdal V, van der Luijt CRB, Chen Y, Kakumanu R, Baidoo EEK, Petzold CJ, Cruz-Morales P, Keasling JD. 2024. Edible mycelium bioengineered for enhanced nutritional value and sensory appeal using a modular synthetic biology toolkit. Nat. Commun. 15:2099.

Manni M, Berkeley MR, Seppey M, Simão FA, Zdobnov EM. 2021. BUSCO Update: Novel and Streamlined Workflows along with Broader and Deeper Phylogenetic Coverage for Scoring of Eukaryotic, Prokaryotic, and Viral Genomes. Mol. Biol. Evol. 38:4647–4654.

Martin JGP, Cotter PD. 2023. Filamentous fungi in artisanal cheeses: A problem to be avoided or a market opportunity? Heliyon 9:e15110.

Melo NR, Moran GP, Warrilow AGS, Dudley E, Smith SN, Sullivan DJ, Lamb DC, Kelly DE, Coleman DC, Kelly SL. 2008. CYP56 (Dit2p) in Candida albicans: Characterization and Investigation of Its Role in Growth and Antifungal Drug Susceptibility. Antimicrob. Agents Chemother. 52:3718–3724.

Moran NA, Jarvik T. 2010. Lateral transfer of genes from fungi underlies carotenoid production in aphids. Science 328:624–627.

Mulet JM, Leube MP, Kron SJ, Rios G, Fink GR, Serrano R. 1999. A Novel Mechanism of Ion Homeostasis and Salt Tolerance in Yeast: the Hal4 and Hal5 Protein Kinases Modulate the Trk1-Trk2 Potassium Transporter. Mol. Cell. Biol. 19:3328–3337.

Navarro C, Navarro MA, Leyva A. 2022. Arsenic perception and signaling: The yet unexplored world. Front. Plant Sci. 13:993484.

Nickles GR, Oestereicher B, Keller NP, Drott MT. 2023. Mining for a new class of fungal natural products: the evolution, diversity, and distribution of isocyanide synthase biosynthetic gene clusters. Nucleic Acids Res. 51:7220–7235.

Norling M, Jareborg N, Dainat J. 2018. EMBLmyGFF3: a converter facilitating genome annotation submission to European Nucleotide Archive. BMC Res. Notes 11:584.

Novo M, Bigey F, Beyne E, Galeote V, Gavory F, Mallet S, Cambon B, Legras J-L, Wincker P, Casaregola S, et al. 2009. Eukaryote-to-eukaryote gene transfer events revealed by the genome sequence of the wine yeast Saccharomyces cerevisiae EC1118. Proc. Natl. Acad. Sci. 106:16333–16338.

Palmer JM, Stajich J. 2020. Funannotate v1.8.1: Eukaryotic genome annotation.

Paradis E, Schliep K. 2019. ape 5.0: an environment for modern phylogenetics and evolutionary analyses in R. Bioinformatics 35:526–528.

Park WH. 2020. Upregulation of thioredoxin and its reductase attenuates arsenic trioxide-induced growth suppression in human pulmonary artery smooth muscle cells by reducing oxidative stress. Oncol. Rep. 43:358–367.

Patel KS, Pandey PK, Martín-Ramos P, Corns WT, Varol S, Bhattacharya P, Zhu Y. 2023. A review on arsenic in the environment: contamination, mobility, sources, and exposure. RSC Adv. 13:8803–8821.

Peck LD, Llewellyn T, Bennetot B, O’Donnell S, Nowell RW, Ryan MJ, Flood J, Vega RCR de la, Ropars J, Giraud T, et al. 2023. Horizontal transfers between fungal Fusarium species contributed to successive outbreaks of coffee wilt disease.: 2023.12.22.572981. Available from: https://www.biorxiv.org/content/10.1101/2023.12.22.572981v1

Perrone G, Samson RA, Frisvad JC, Susca A, Gunde-Cimerman N, Epifani F, Houbraken J. 2015. *Penicillium salamii*, a new species occurring during seasoning of dry-cured meat. Int. J. Food Microbiol. 193:91–98.

Pfaller MA, Pappas PG, Wingard JR. 2006. Invasive Fungal Pathogens: Current Epidemiological Trends. Clin. Infect. Dis. 43:S3–S14.

Piñeiro C, Abuín JM, Pichel JC. 2020. Very Fast Tree: speeding up the estimation of phylogenies for large alignments through parallelization and vectorization strategies. Bioinformatics 36:4658–4659.

Pitt JI, Hocking AD. 2009. Fungi and Food Spoilage. Springer Science & Business Media

Prjibelski A, Antipov D, Meleshko D, Lapidus A, Korobeynikov A. 2020. Using SPAdes De Novo Assembler. Curr. Protoc. Bioinforma. 70:e102.

Quinlan AR, Hall IM. 2010. BEDTools: a flexible suite of utilities for comparing genomic features. Bioinformatics 26:841–842.

R Core Team. 2024. R: A language and environment for statistical computing. R Found. Stat. Comput. Vienna Austria [Internet]. Available from: https://www.R-project.org/

Rhie A, Walenz BP, Koren S, Phillippy AM. 2020. Merqury: reference-free quality, completeness, and phasing assessment for genome assemblies. Genome Biol. 21:245.

Rhodes J, Abdolrasouli A, Dunne K, Sewell TR, Zhang Y, Ballard E, Brackin AP, van Rhijn N, Chown H, Tsitsopoulou A, et al. 2022. Population genomics confirms acquisition of drug-resistant Aspergillus fumigatus infection by humans from the environment. Nat. Microbiol. 7:663–674.

Robellet X, Flipphi M, Pégot S, MacCabe AP, Vélot C. 2008. AcpA, a member of the GPR1/FUN34/YaaH membrane protein family, is essential for acetate permease activity in the hyphal fungus Aspergillus nidulans. Biochem. J. 412:485–493.

Ropars J, Didiot E, Rodríguez de la Vega RC, Bennetot B, Coton M, Poirier E, Coton E, Snirc A, Le Prieur S, Giraud T. 2020. Domestication of the Emblematic White Cheese-Making Fungus Penicillium camemberti and Its Diversification into Two Varieties. Curr. Biol. CB 30:4441–4453.e4.

Ropars J, Rodríguez de la Vega RC, López-Villavicencio M, Gouzy J, Sallet E, Dumas É, Lacoste S, Debuchy R, Dupont J, Branca A, et al. 2015. Adaptive Horizontal Gene Transfers between Multiple Cheese-Associated Fungi. Curr. Biol. 25:2562–2569.

Sayers EW, Bolton EE, Brister JR, Canese K, Chan J, Comeau DC, Connor R, Funk K, Kelly C, Kim S, et al. 2022. Database resources of the national center for biotechnology information. Nucleic Acids Res. 50:D20–D26.

Schrader L, Schmitz J. 2019. The impact of transposable elements in adaptive evolution. Mol. Ecol. 28:1537–1549.

Sinzelle L, Kapitonov VV, Grzela DP, Jursch T, Jurka J, Izsvák Z, Ivics Z. 2008. Transposition of a reconstructed Harbinger element in human cells and functional homology with two transposon-derived cellular genes. Proc. Natl. Acad. Sci. 105:4715–4720.

Skrzypek MS, Binkley J, Binkley G, Miyasato SR, Simison M, Sherlock G. 2017. The Candida Genome Database (CGD): incorporation of Assembly 22, systematic identifiers and visualization of high throughput sequencing data. Nucleic Acids Res. 45:D592–D596.

Smith MR. 2019. TreeTools: create, modify and analyse phylogenetic trees. Comprehensive R Archive Network

Stefanini I, Di Paola M, Liti G, Marranci A, Sebastiani F, Casalone E, Cavalieri D. 2022. Resistance to Arsenite and Arsenate in Saccharomyces cerevisiae Arises through the Subtelomeric Expansion of a Cluster of Yeast Genes. Int. J. Environ. Res. Public. Health 19:8119.

Sugui JA, Vinh DC, Nardone G, Shea YR, Chang YC, Zelazny AM, Marr KA, Holland SM, Kwon-Chung KJ. 2010. Neosartorya udagawae (Aspergillus udagawae), an Emerging Agent of Aspergillosis: How Different Is It from Aspergillus fumigatus? J. Clin. Microbiol. 48:220– 228.

Urquhart A, Vogan AA, Gluck-Thaler E. 2024. *Starships*: a new frontier for fungal biology. Trends Genet. 40:1060–1073.

Urquhart AS, Chong NF, Yang Y, Idnurm A. 2022. A large transposable element mediates metal resistance in the fungus Paecilomyces variotii. Curr. Biol. CB 32:937–950.e5.

Urquhart AS, Gluck-Thaler E, Vogan AA. 2024. Gene acquisition by giant transposons primes eukaryotes for rapid evolution via horizontal gene transfer. Sci. Adv. 10:eadp8738.

Urquhart AS, Vogan AA, Gardiner DM, Idnurm A. 2023. Starships are active eukaryotic transposable elements mobilized by a new family of tyrosine recombinases. Proc. Natl. Acad. Sci. 120:e2214521120.

Vogan AA, Ament-Velásquez SL, Bastiaans E, Wallerman O, Saupe SJ, Suh A, Johannesson H. 2021. The Enterprise, a massive transposon carrying Spok meiotic drive genes. Genome Res. 31:789–798.

Wang L-G, Lam TT-Y, Xu S, Dai Z, Zhou L, Feng T, Guo P, Dunn CW, Jones BR, Bradley T, et al. 2020. Treeio: An R Package for Phylogenetic Tree Input and Output with Richly Annotated and Associated Data. Mol. Biol. Evol. 37:599–603.

Wickham H. 2016. ggplot2: Elegant Graphics for Data Analysis. Springer-Verlag New York Available from: https://ggplot2.tidyverse.org

William VU, Magpantay HD. 2024. Arsenic and Microorganisms: Genes, Molecular Mechanisms, and Recent Advances in Microbial Arsenic Bioremediation. Microorganisms 12:74.

Wolfe BE, Dutton RJ. 2015. Fermented Foods as Experimentally Tractable Microbial Ecosystems. Cell 161:49–55.

Wysocki R, Fortier P-K, Maciaszczyk E, Thorsen M, Leduc A, Odhagen Å, Owsianik G, Ulaszewski S, Ramotar D, Tamás MJ. 2004. Transcriptional Activation of Metalloid Tolerance Genes in Saccharomyces cerevisiae Requires the AP-1–like Proteins Yap1p and Yap8p. Mol. Biol. Cell 15:2049–2060.

Xie L, Zhou L, Zhang R, Zhou H, Yang Y. 2024. Material Composition Characteristics of Aspergillus cristatus under High Salt Stress through LC–MS Metabolomics. Molecules 29:2513.

Yang Z, Huang J, Geng J, Nair U, Klionsky DJ. 2006. Atg22 Recycles Amino Acids to Link the Degradative and Recycling Functions of Autophagy. Mol. Biol. Cell 17:5094–5104.

Yin Y, Miao J, Shao W, Liu X, Zhao Y, Ma Z. 2023. Fungicide Resistance: Progress in Understanding Mechanism, Monitoring, and Management. Phytopathology® 113:707– 718.

Yu G. 2023. aplot: Decorate a “ggplot” with Associated Information. Available from: https://github.com/YuLab-SMU/aplot

Yu G, Smith DK, Zhu H, Guan Y, Lam TT-Y. 2017. ggtree: an r package for visualization and annotation of phylogenetic trees with their covariates and other associated data. Methods Ecol. Evol. 8:28–36.

Zhang H-H, Peccoud J, Xu M-R-X, Zhang X-G, Gilbert C. 2020. Horizontal transfer and evolution of transposable elements in vertebrates. Nat. Commun. 11:1362.

