## Supplemental Figures for "Harbouring *Starships*: The accumulation of large Horizontal Gene Transfers in Domesticated and Pathogenic Fungi"

**S. O'Donnell**: Conceptualization, Data Curation, Formal Analysis, Investigation, Methodology, Software, Validation, Visualisation, Writing-original draft, Writing-review and editing. **G. Rezende**: Investigation. **J.-P. Vernadet**: Data Curation, Investigation, Software. **A. Snirc**: Investigation. **J. Ropars**: Conceptualization, Data Curation, Funding acquisition, Investigation, Methodology, Project administration, Resources, Supervision, Validation, Writing-original draft, Writing-review and editing.

ORCIDs

Samuel O'Donnell: https://orcid.org/0000-0003-2447-1993

Jeanne Ropars: https://orcid.org/0000-0002-3740-9673

Key words

adaptation; transposable elements; moulds; filamentous fungi; fermented food; clinical; *Penicillium*; *Aspergillus*; human-made environments; accurate phylogeny; HGT

### Abstract

Human-related environments, including food and clinical settings, present microorganisms with atypical and challenging conditions that necessitate adaptation. Several cases of novel horizontally acquired genetic material associated with adaptive traits have been recently described, contained within giant transposons named *Starships.* While a handful of *Starships* have been identified in domesticated species, their abundance has not yet been systematically explored in human-associated fungi. Here, we investigated whether *Starships* have shaped the genomes of two major genera of fungi occurring in food and clinical environments, *Aspergillus* and *Penicillium.* Using seven independent domestication events, we found in all cases that the domesticated strains or species exhibited significantly greater *Starship* content compared with close relatives from non-human-related environments. We found a similar pattern in clinical contexts. Our findings have clear implications for agriculture, human health and the food industry as we implicate *Starships* as a widely recurrent mechanism of gene transfer aiding the rapid adaptation of fungi to novel environments.

### Supplementary figures and legends

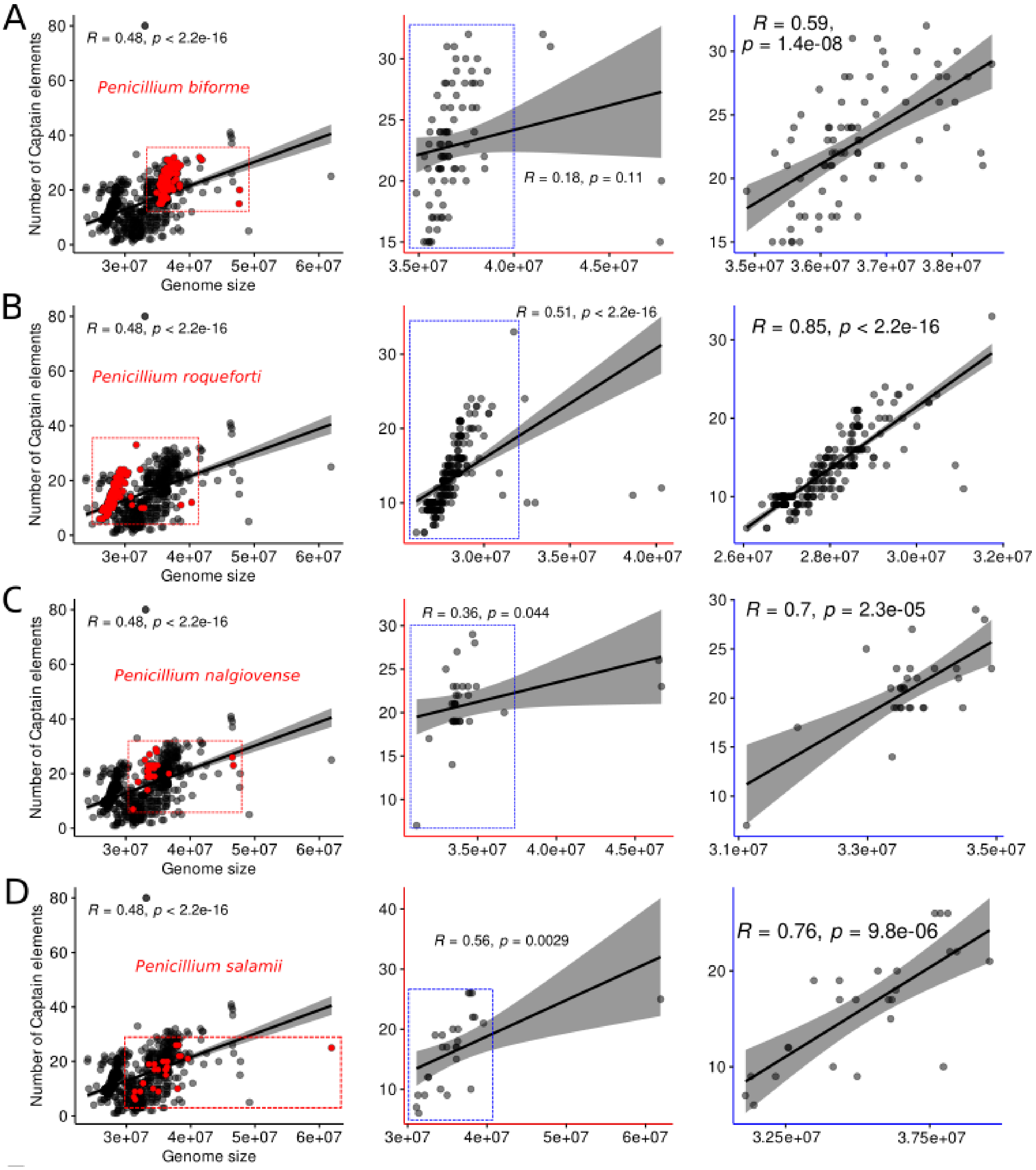


**fig. S1**. **Measuring the linear correlation between the number of Captains and genome size per assembly for *Penicillium* strains.** The Pearson correlation coefficient (R) was calculated for each set of genomes. Subsets of species with a large number of assemblies were selected, written in red in the far left column, and are specific to the row (e.g. row A contains *P. biforme*). All those in the far left column contain the entire genus-assembly dataset, with the assemblies for each row-specific species coloured red and their spread outlined in the dotted red square. The middle column contains the subset of assemblies for each row-species with the dotted blue square removing outliers with large genome sizes. The far right column contains the subset of non-outlier genomes per species (outlined in blue in the middle column).

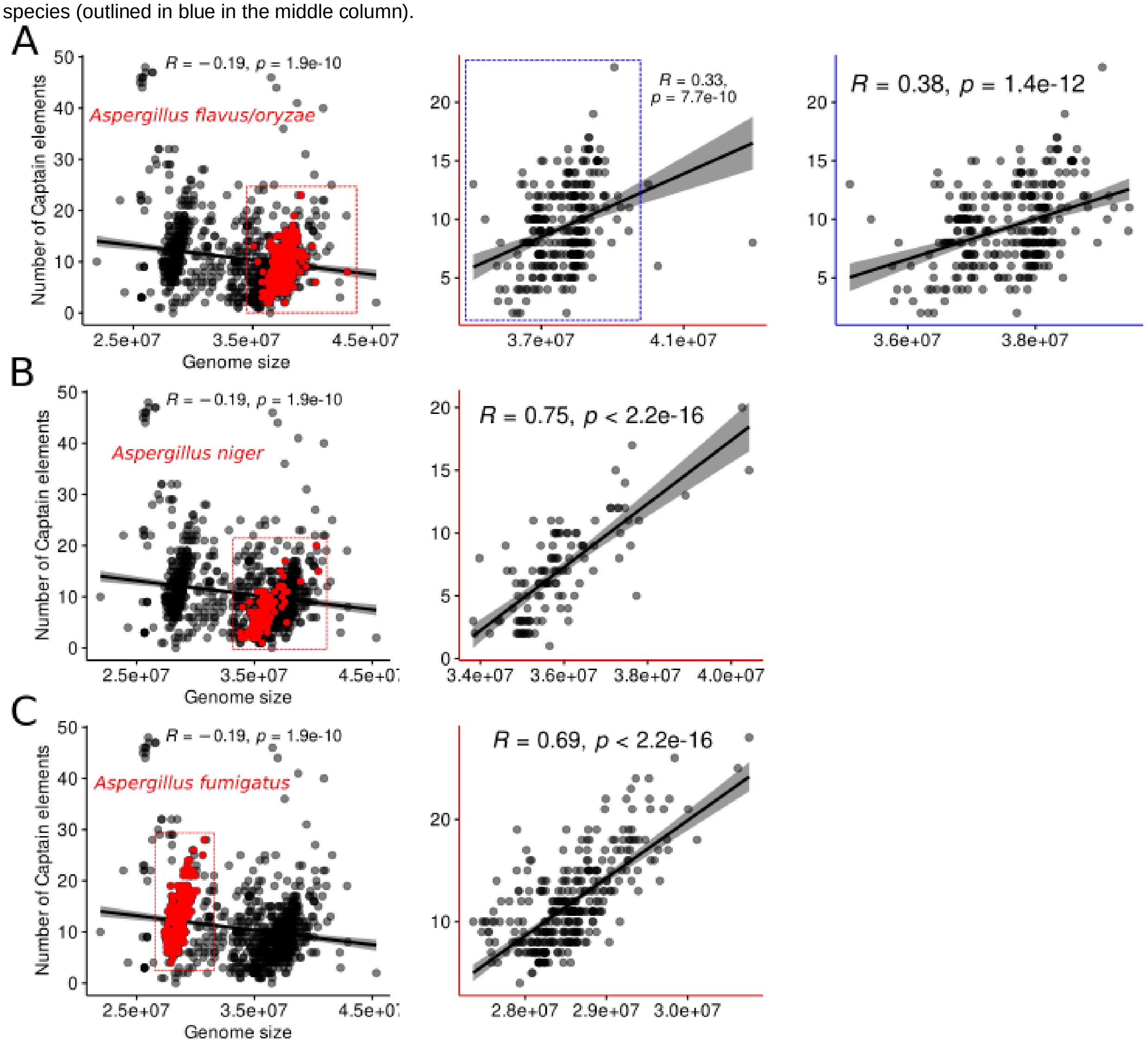


**fig. S2**. **Measuring the linear correlation between the number of Captains and genome size per assembly for *Aspergillus* strains**. The Pearson correlation coefficient (R) was calculated for each set of genomes. Subsets of species with a large number of assemblies were selected, written in red in the far left column, and are specific to the row (e.g. row C contains *A. fumigatus*). All those in the far left column contain the entire genus-assembly dataset, with the assemblies for each row-specific species coloured red and their spread outlined in the dotted red square. The middle column contains the subset of assemblies for each row-species with the dotted blue square removing outliers with large genome sizes. The far right column contains the subset of non-outlier genomes per species. Assemblies for *A. niger* and *A. fumigatus* contained no outliers.

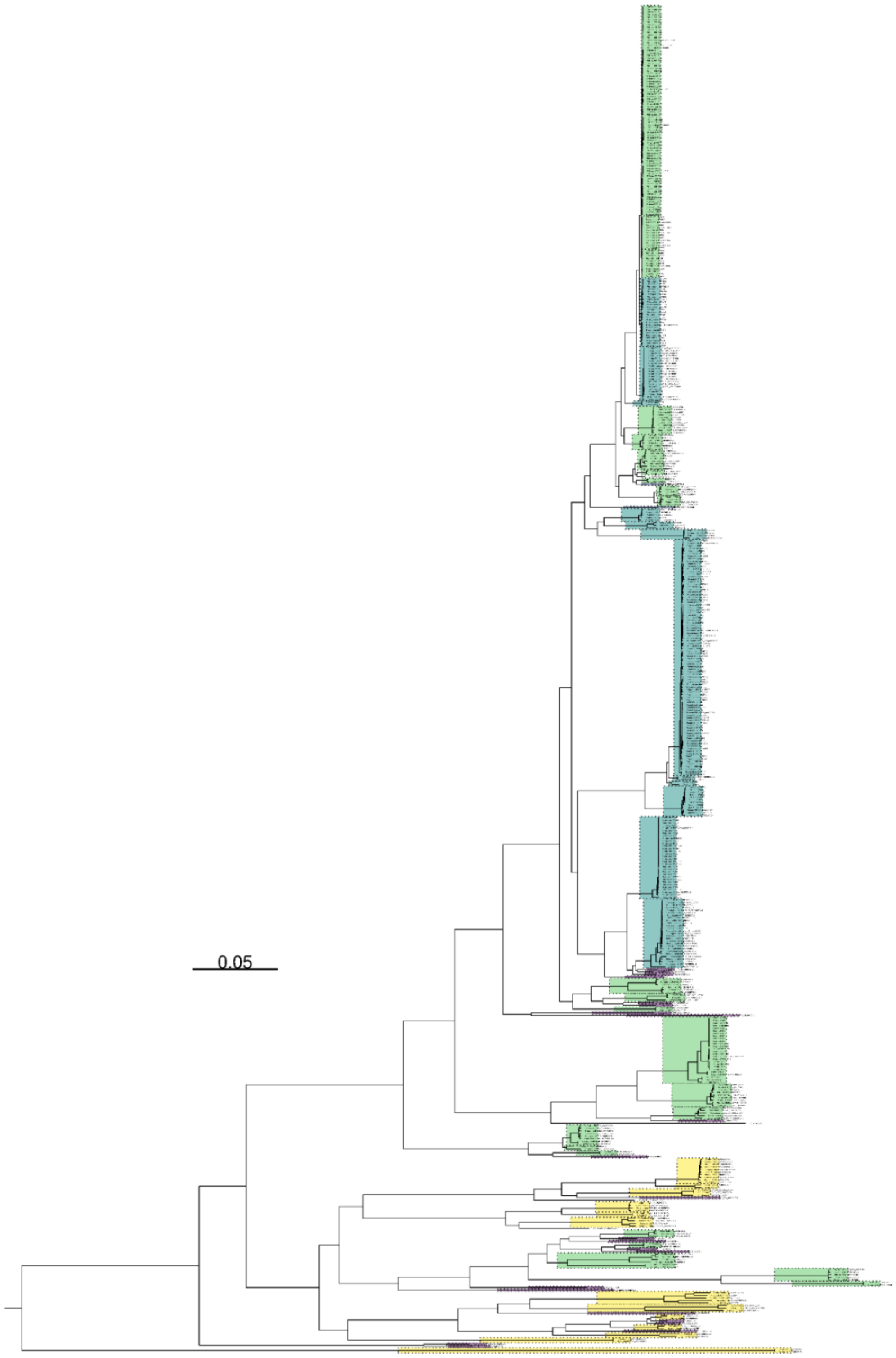


**fig. S3**. **Maximum-Likelihood tree showing phylogenetic relationships of 525 *Penicillium* based on 3154 single copy BUSCO genes**. Each branch terminal node contains the identifying name for each assembly, species-specific nodes have SH-like support values and each species specific clade is highlighted in coloured boxes with dotted outlines. Tree was rooted using *Aspergillus* genomes which have been removed. The scale bar represents 0.05 substitutions per site for branch lengths.

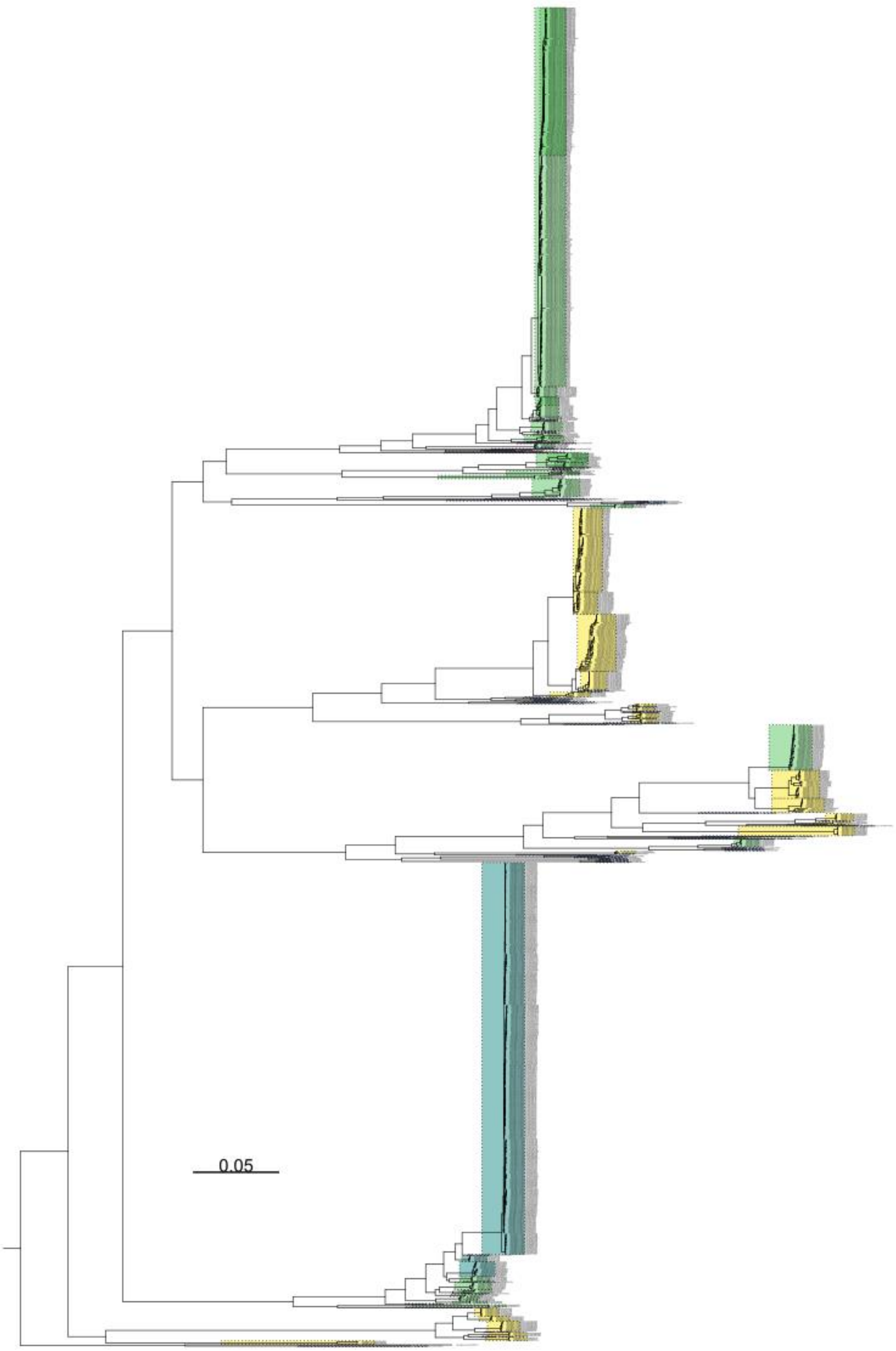


**fig. S4. Maximum-Likelihood tree showing phylogenetic relationships of 1148 *Aspergillus* based on 3258 single copy BUSCO genes**. Each branch terminal node contains the identifying name for each assembly and each species specific clade is highlighted in coloured boxes with dotted outlines. Tree was rooted using Penicillium genomes which have been removed. The scale bar represents 0.05 substitutions per site for branch lengths.

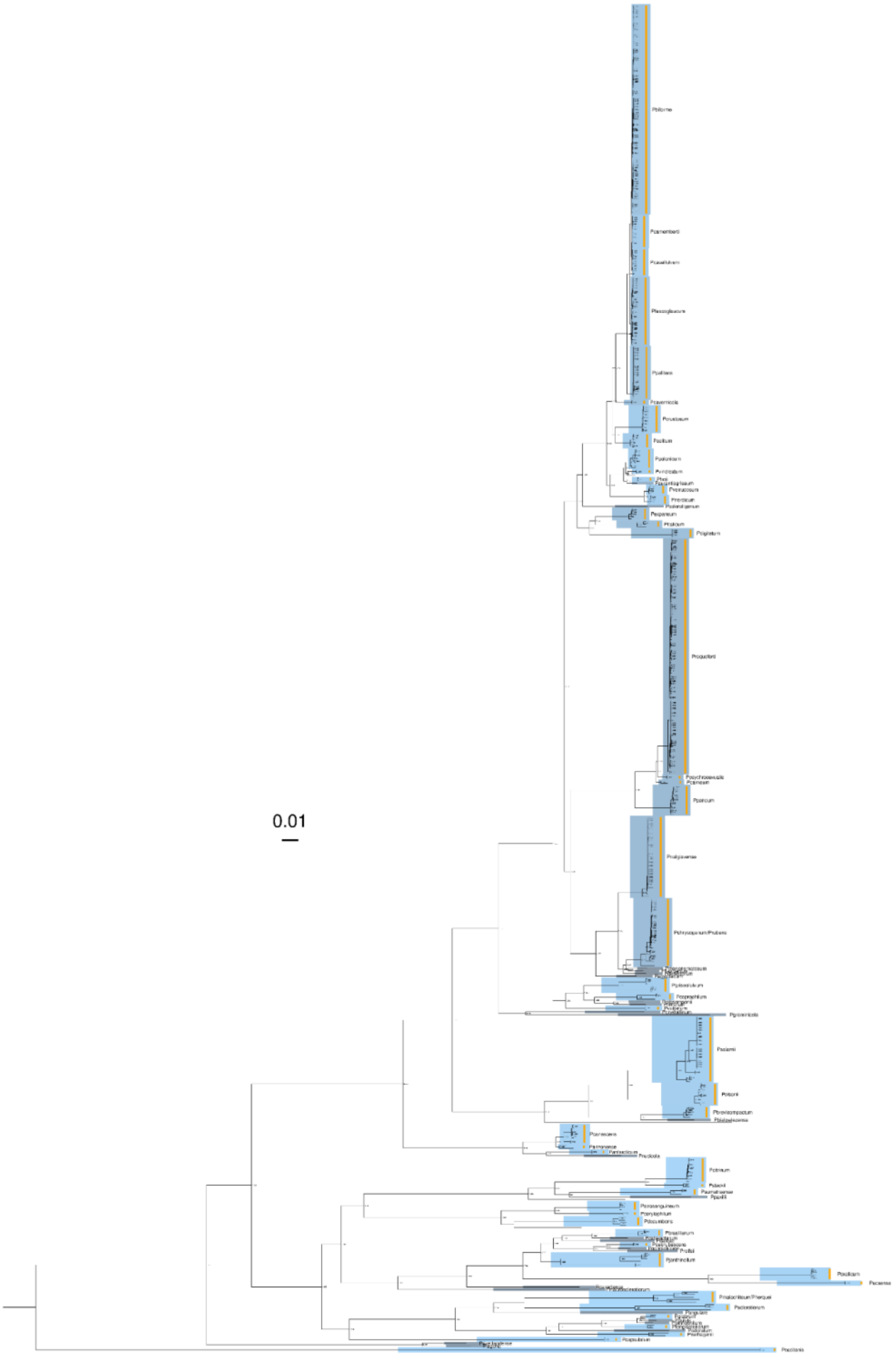


**fig. S5**. **Maximum-Likelihood tree showing phylogenetic relationships of 525 *Penicillium* based on 3154 single copy BUSCO genes**. Each branch terminal node contains the identifying name for each assembly, species-specific nodes have SH-like support values and each species specific clade is highlighted in coloured boxes with dotted outlines. The scale bar represents 0.01 substitutions per site for branch lengths. Tree was rooted using *Aspergillus* genomes which have been removed. The scale bar represents 0.05 substitutions per site for branch lengths.

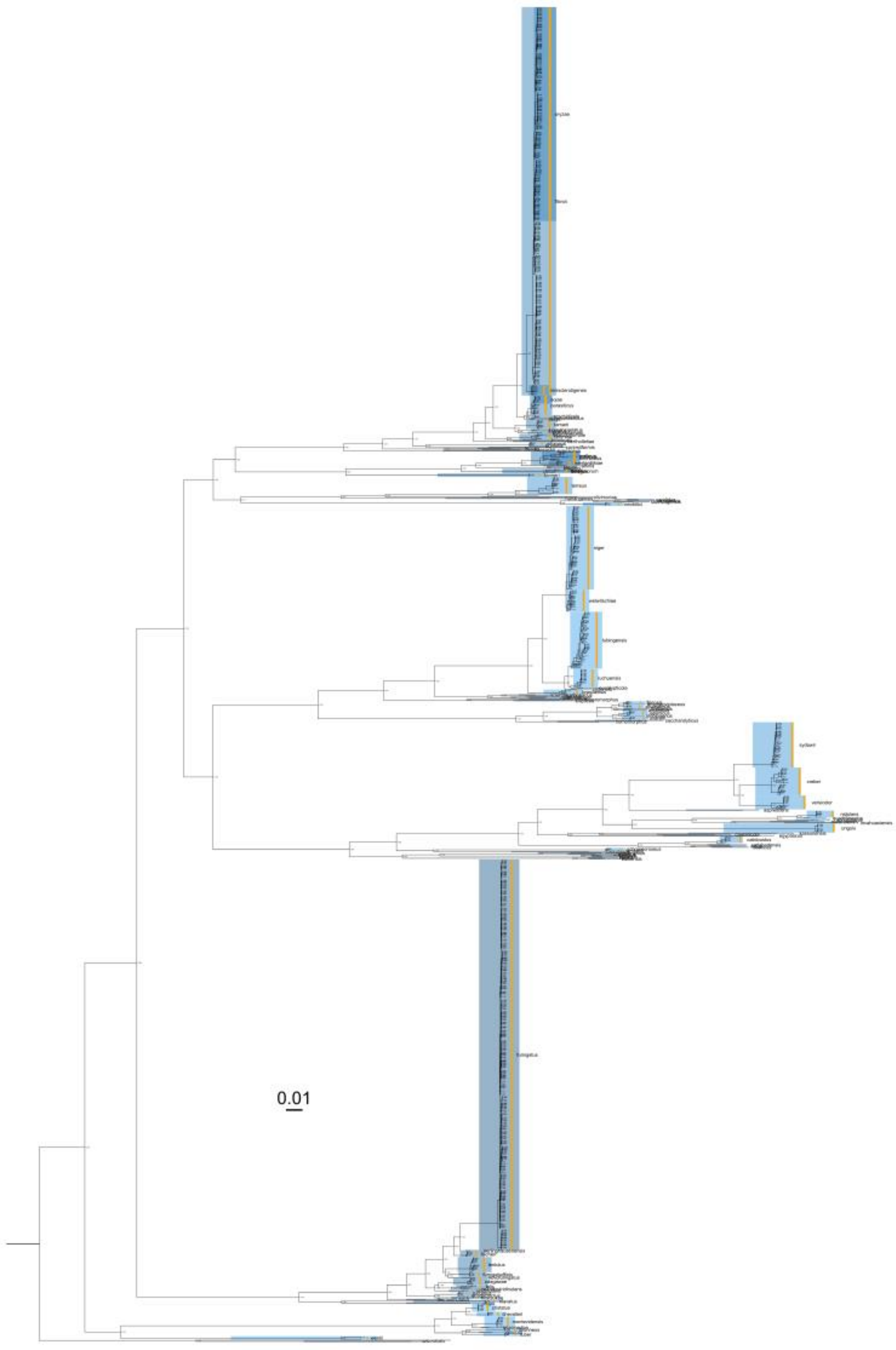

**fig. S6**. **Maximum-Likelihood tree showing phylogenetic relationships of 1148 *Aspergillus* based on 3258 single copy BUSCO genes.** Species-specific nodes have SH-like support values and each species specific clade is highlighted in coloured boxes with dotted outlines. Tree was rooted using Penicillium genomes which have been removed. The scale bar represents 0.01 substitutions per site for branch lengths.

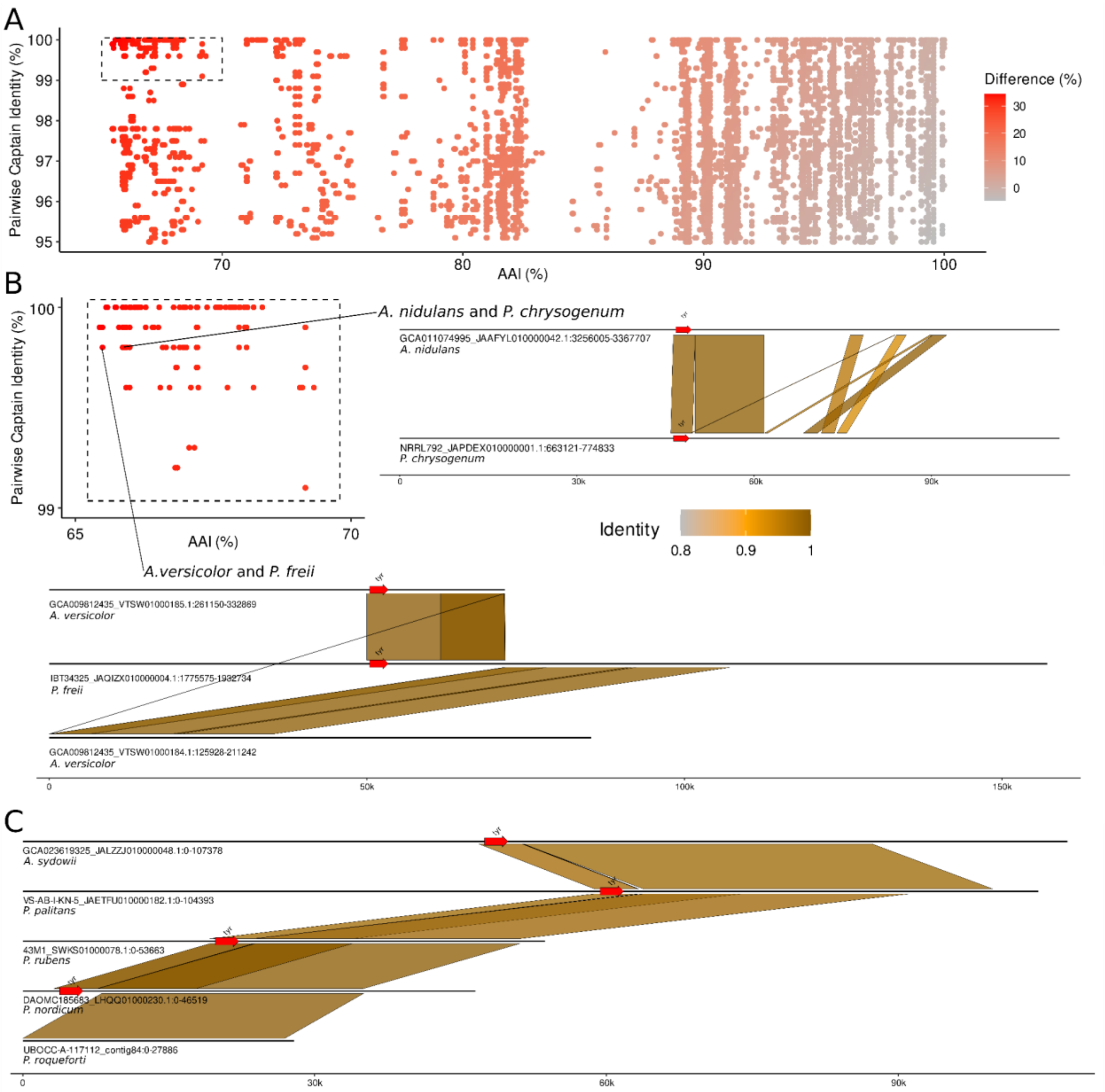


**fig. S7**. Average Amino acid Identity (AAI) compared to Captain protein similarity with examples of genome alignment comprising captains from distant genomes**.** (A) All pairwise Captain proteins with >95% similarity compared to the AAI from their respective genomes calculated using 50 BUSCO proteins. Colour denotes the difference between the two similarity measures. (B) Region of A expanded to show the pairwise comparisons with the largest difference in Captain and AAI similarity. Two examples from this region have then been illustrated further using genome wide alignment. The red arrow denotes the tyrosine recombinase/Captain. (C) The whole genome alignment between five different genomes each of which has a Captain protein with >95% similarity to one another. In each of the examples the alignments indicate the similar captains were associated with regions of high similarity of >20kb and in unique syntenic regions.

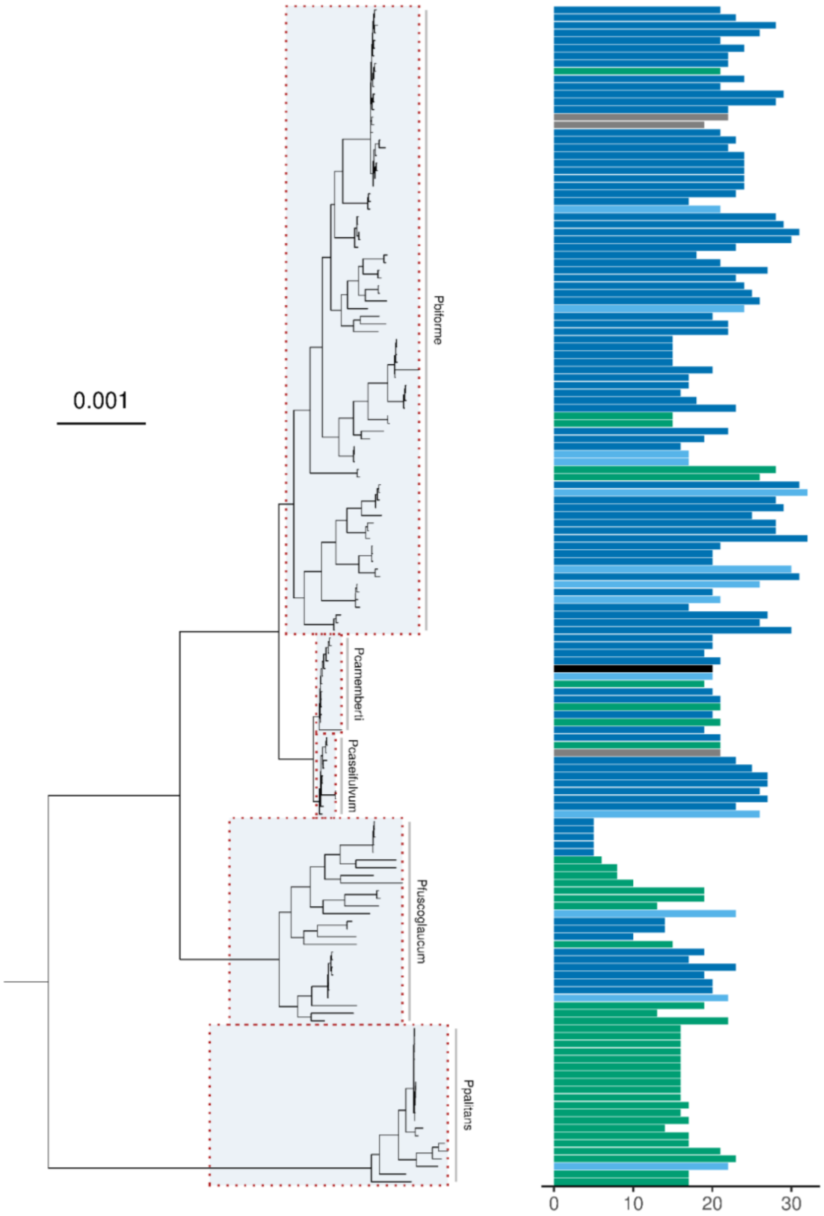


**fig. S8**. **A subset of assemblies (*P. biforme*, *P. camemberti*, *P. caseifulvum*, *P. fuscoglaucum* and *P. palitans*) phylogenetically placed alongside their captain count**. Bars are coloured according to isolation origins (environment = green; environment-clinical = yellow; environment-food = light-blue; food-production=dark-blue; unknown=grey; NA=black).

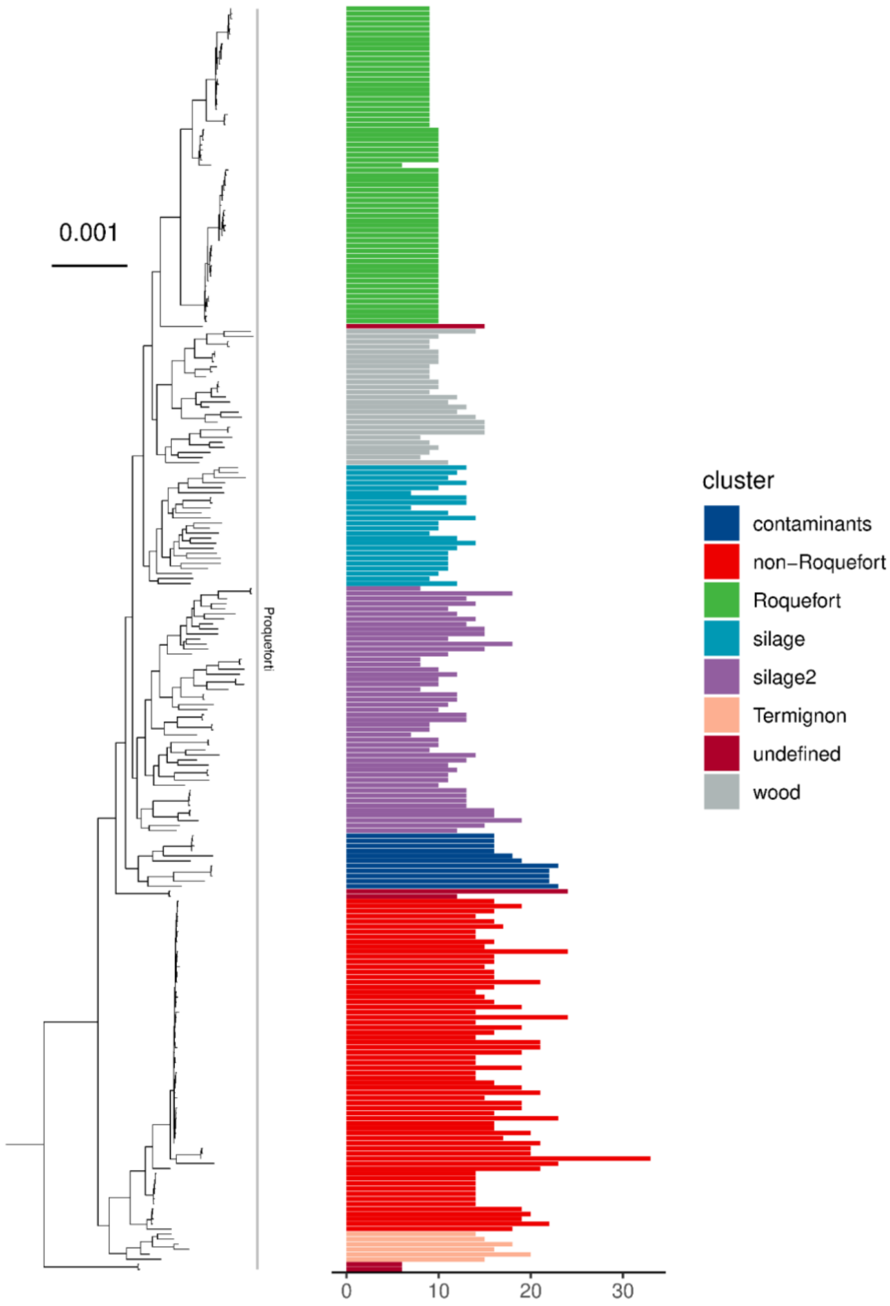


**fig. S9**. **A subset of *Penicillium roqueforti* assemblies phylogenetically placed alongside their Captain count and coloured by the strains cluster named based on their origins.** Bars are coloured according to defined isolation clusters shown in legend.

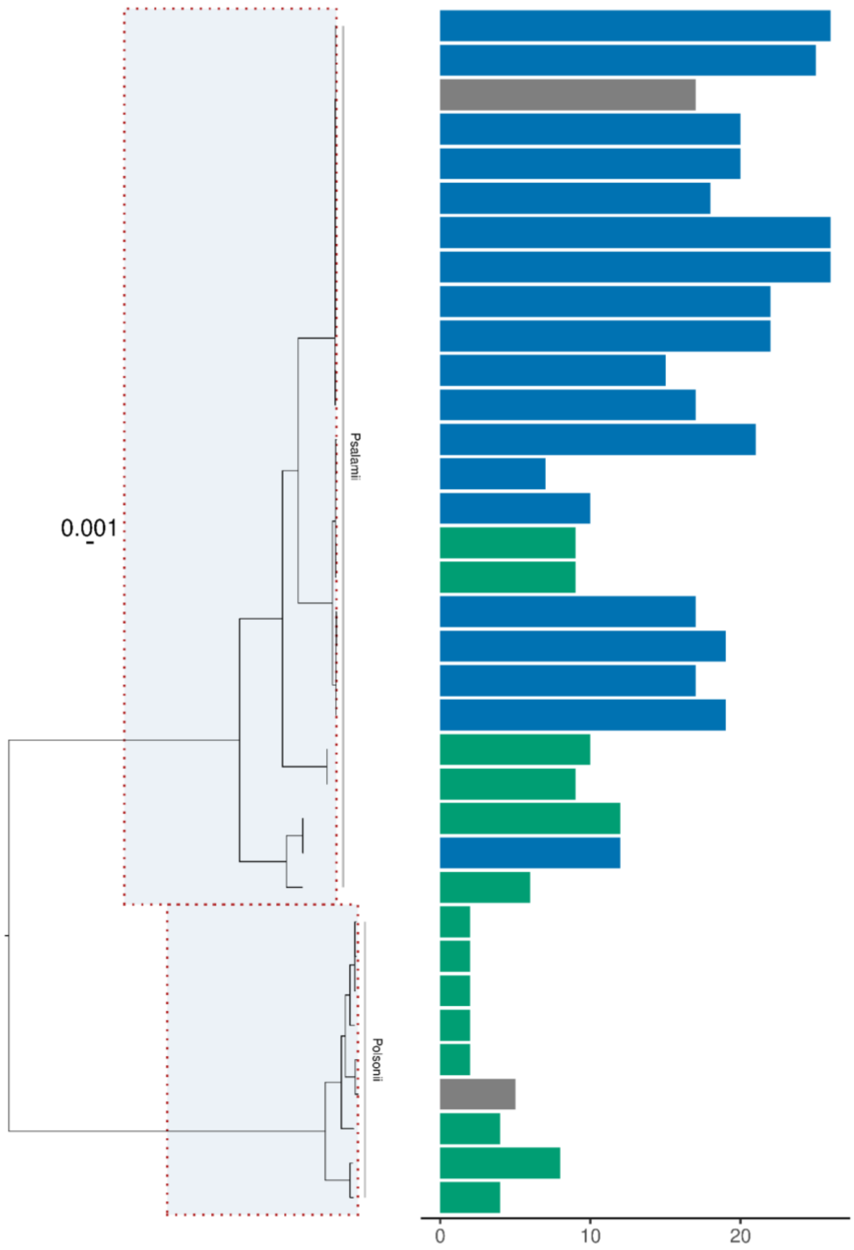


**fig. S10**. **A subset of assemblies (*P. salamii* and *P. olsonii*) phylogenetically placed alongside their captain count.** Bars are coloured according to isolation origins (environment = green; environment-clinical = yellow; environment-food = light-blue; food-production=dark-blue; unknown=grey; NA=black).

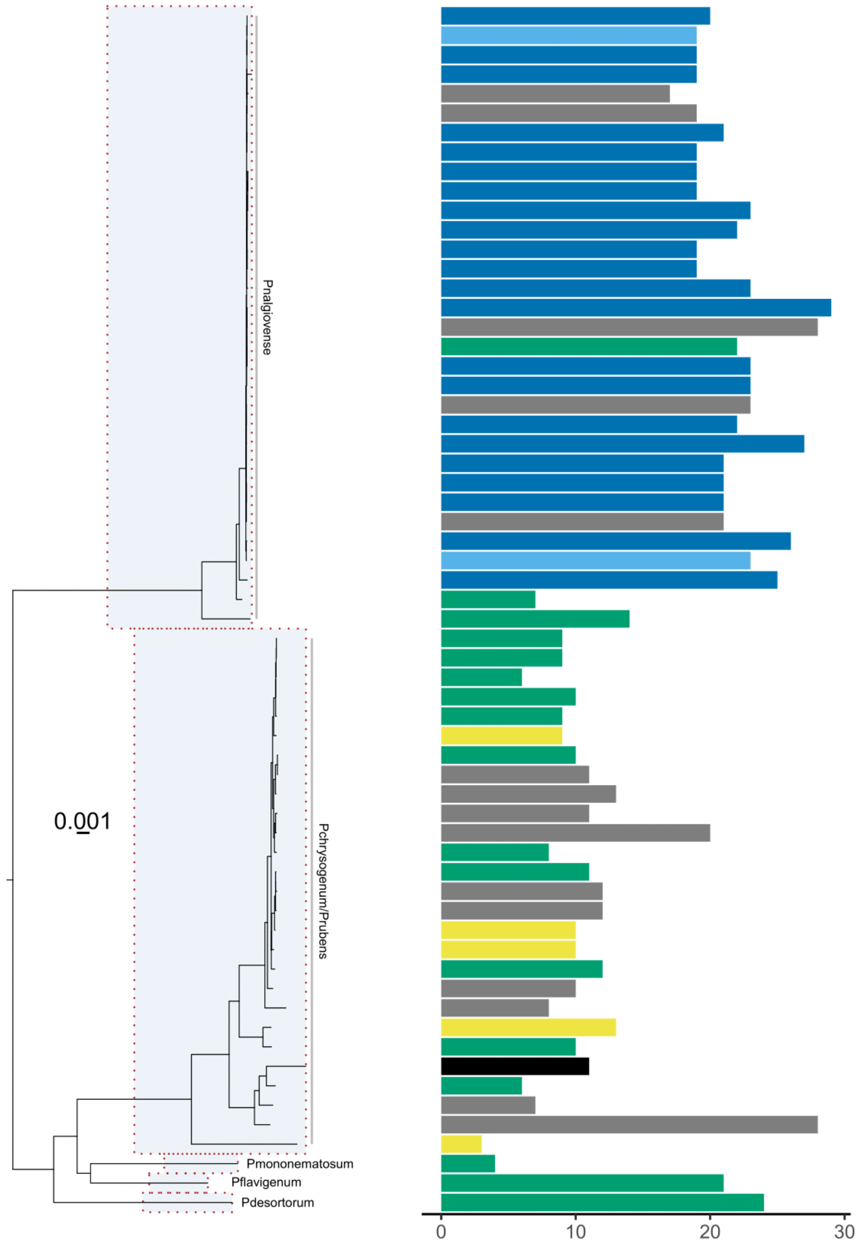


**fig. S11**. **A subset of assemblies (*P. nalgiovense*, *P. chrysogenum/rubens*, *P. mononematosum*, *P. flavigenum* and *P. desertorum*) phylogenetically placed alongside their captain count.** Bars are coloured according to isolation origins (environment = green; environment-clinical = yellow; environment-food = light-blue; food-production=dark-blue; unknown=grey; NA=black).

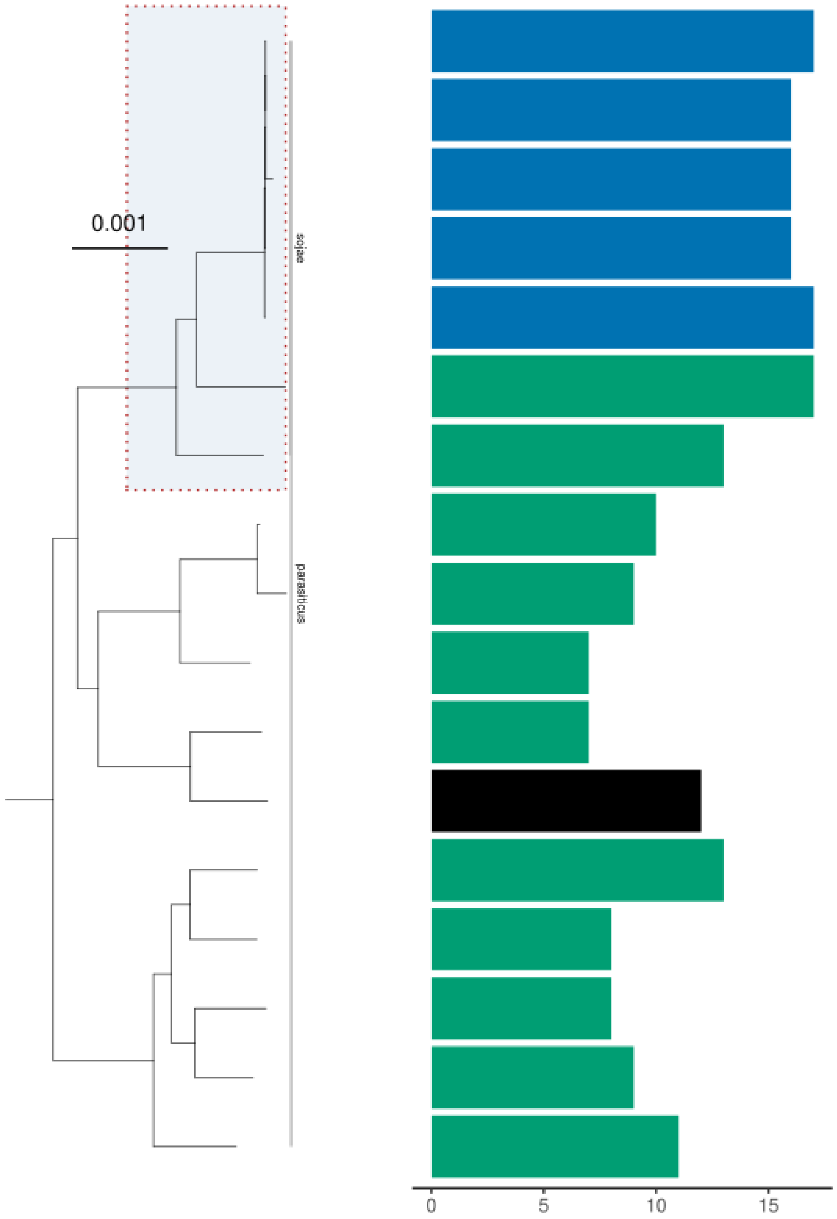


**fig. S12. A subset of assemblies (*A. sojae* and *A. parasiticus*) phylogenetically placed alongside their captain count.** Bars are coloured according to isolation origins (clinical = orange; environment = green; environment (saline) = purple; environment-clinical = yellow; environment-food = light-blue; food-production=dark-blue; unknown=grey; NA=black).

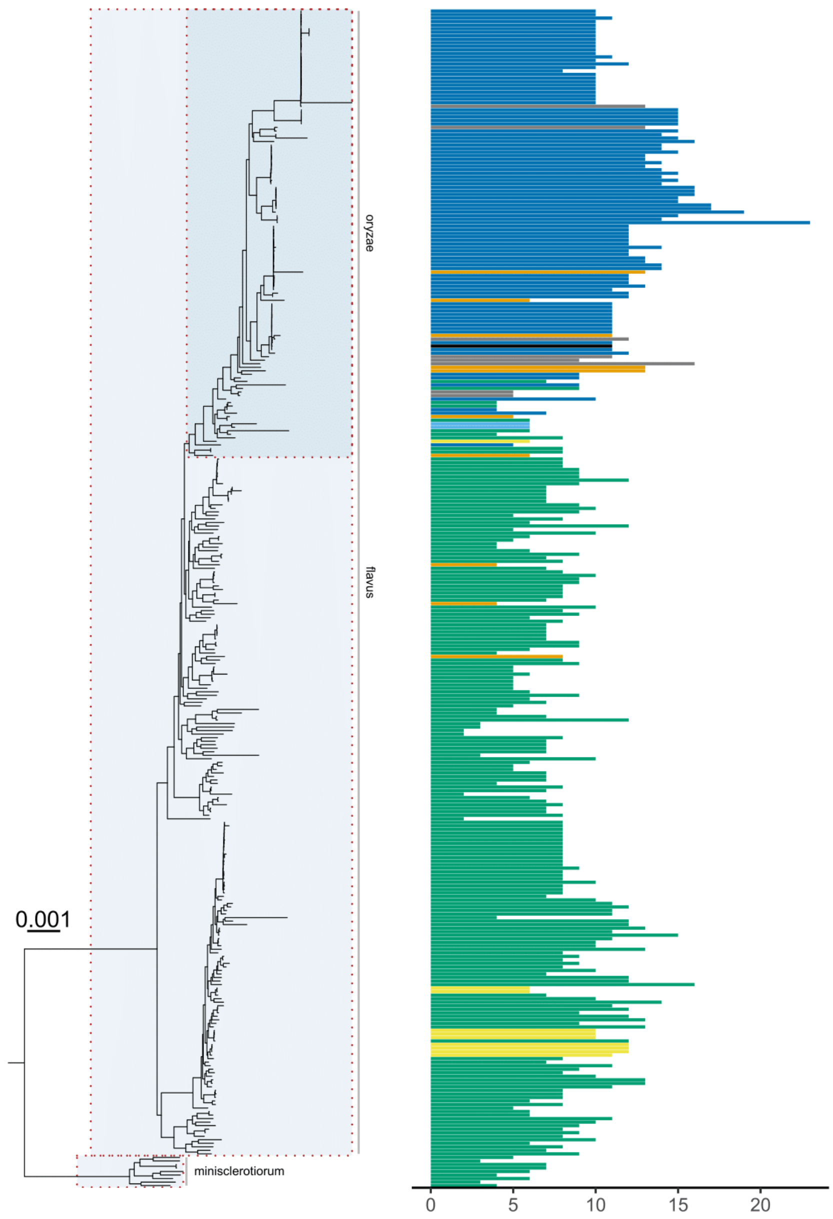


**fig. S13. A subset of assemblies (*A. oryzae*, *A. flavus*, and *A. minisclerotiorum*) phylogenetically placed alongside their captain count.** Bars are coloured according to isolation origins (clinical = orange; environment = green; environment (saline) = purple; environment-clinical = yellow; environment-food = light-blue; food-production=dark-blue; unknown=grey; NA=black).

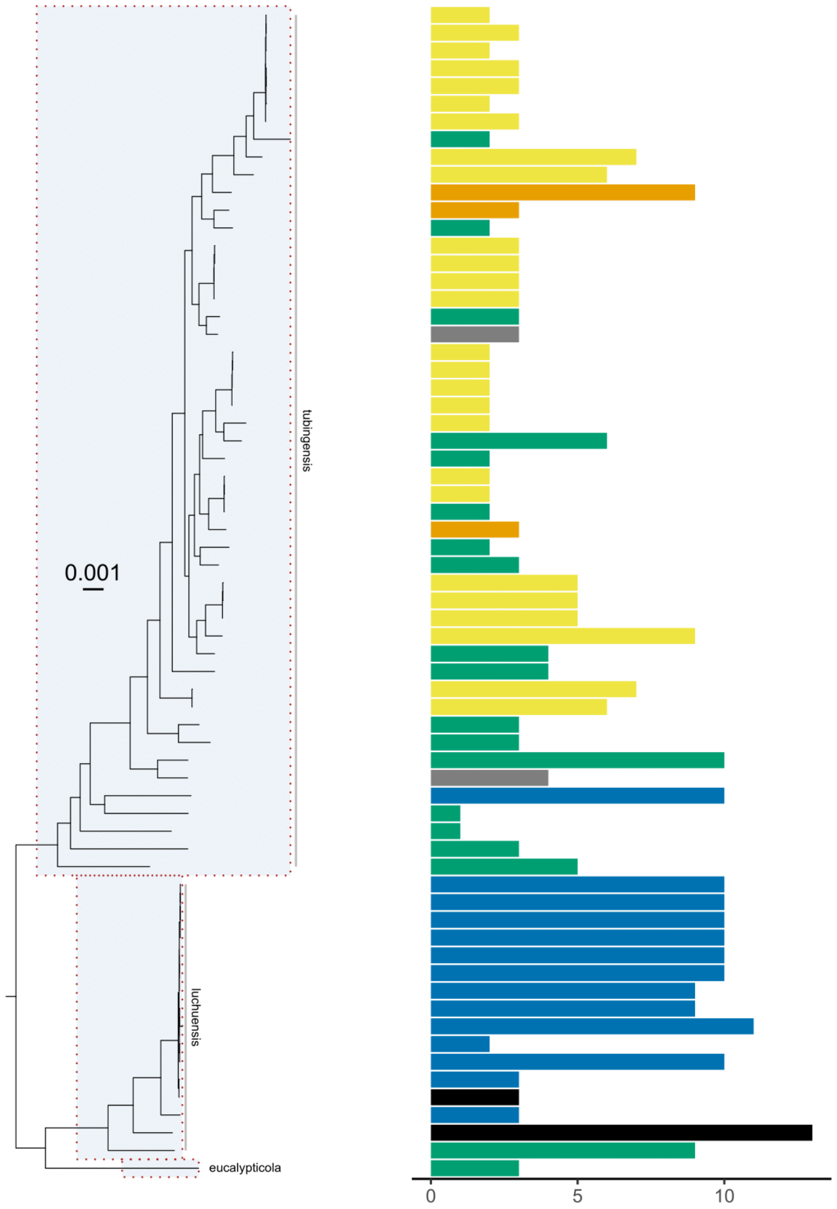


**fig. S14**. **A subset of assemblies (*A. luchuensis*, *A. tubingensis*, and *A. eucalypticola*) phylogenetically placed alongside their captain count**. Bars are coloured according to isolation origins (clinical = orange; environment = green; environment (saline) = purple; environment-clinical = yellow; environment-food = light-blue; food-production=dark-blue; unknown=grey; NA=black).

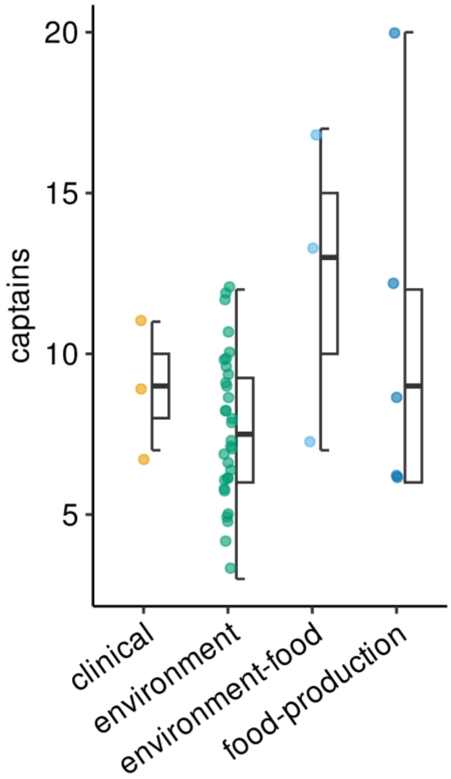


**fig. S15**. **Boxplot of the number of captains found in strains of *Aspergillus niger* split by isolation origins.**

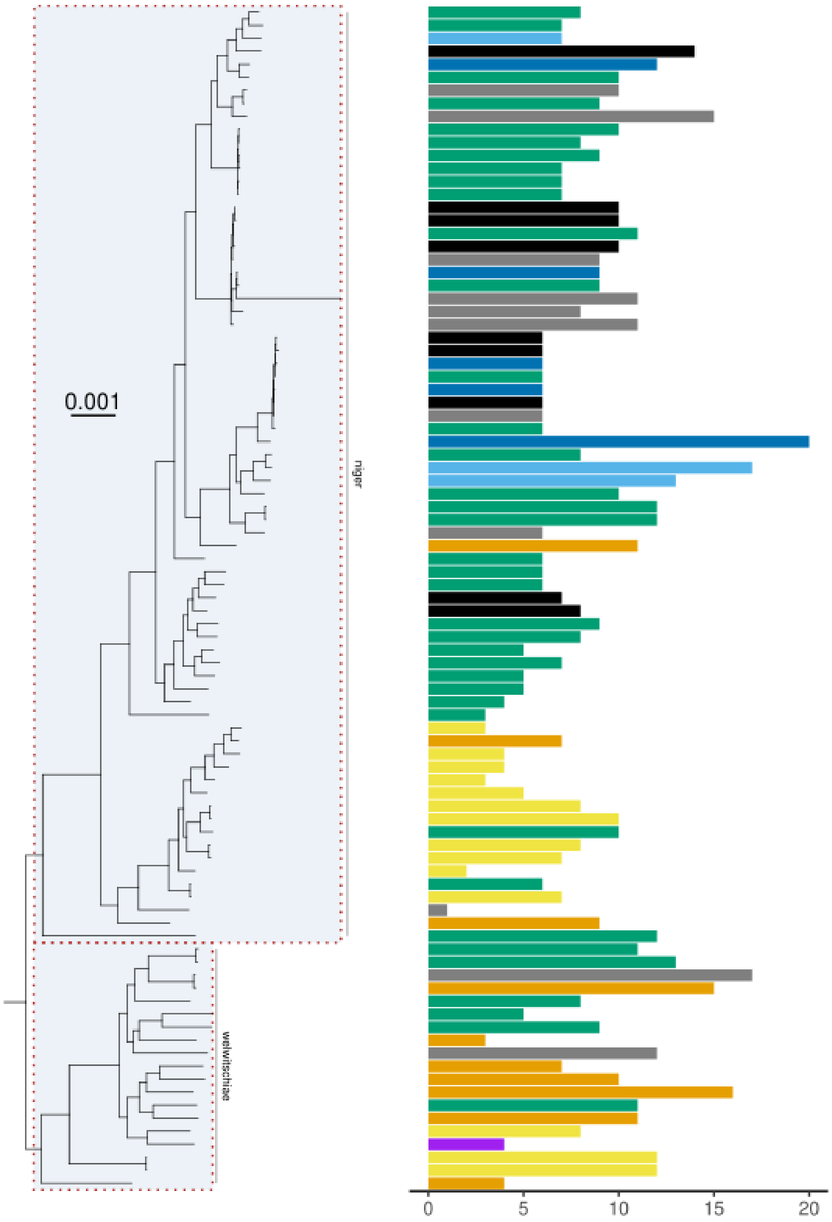


**fig. S16**. **A subset of assemblies (*A. niger* and *A. welwitschiae*) phylogenetically placed alongside their captain count.** Bars are coloured according to isolation origins (clinical = orange; environment = green; environment (saline) = purple; environment-clinical = yellow; environment-food = light-blue; food-production=dark-blue; unknown=grey; NA=black).

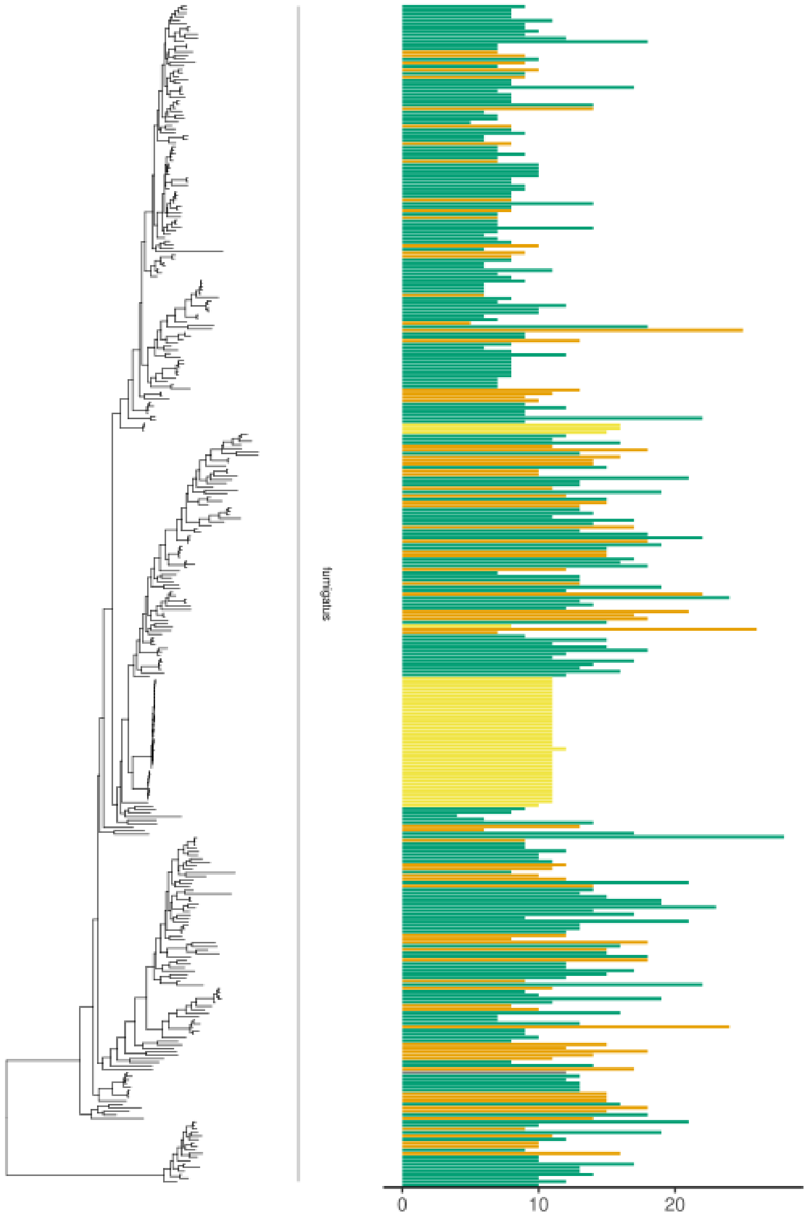


**fig. S17**. **A subset of *Aspergillus fumigatus* assemblies phylogenetically placed alongside their captain count.** Bars are coloured according to isolation origins (clinical = orange; environment = green; environment (saline) = purple; environment-clinical = yellow; environment-food = light-blue; food-production=dark-blue; unknown=grey; NA=black).

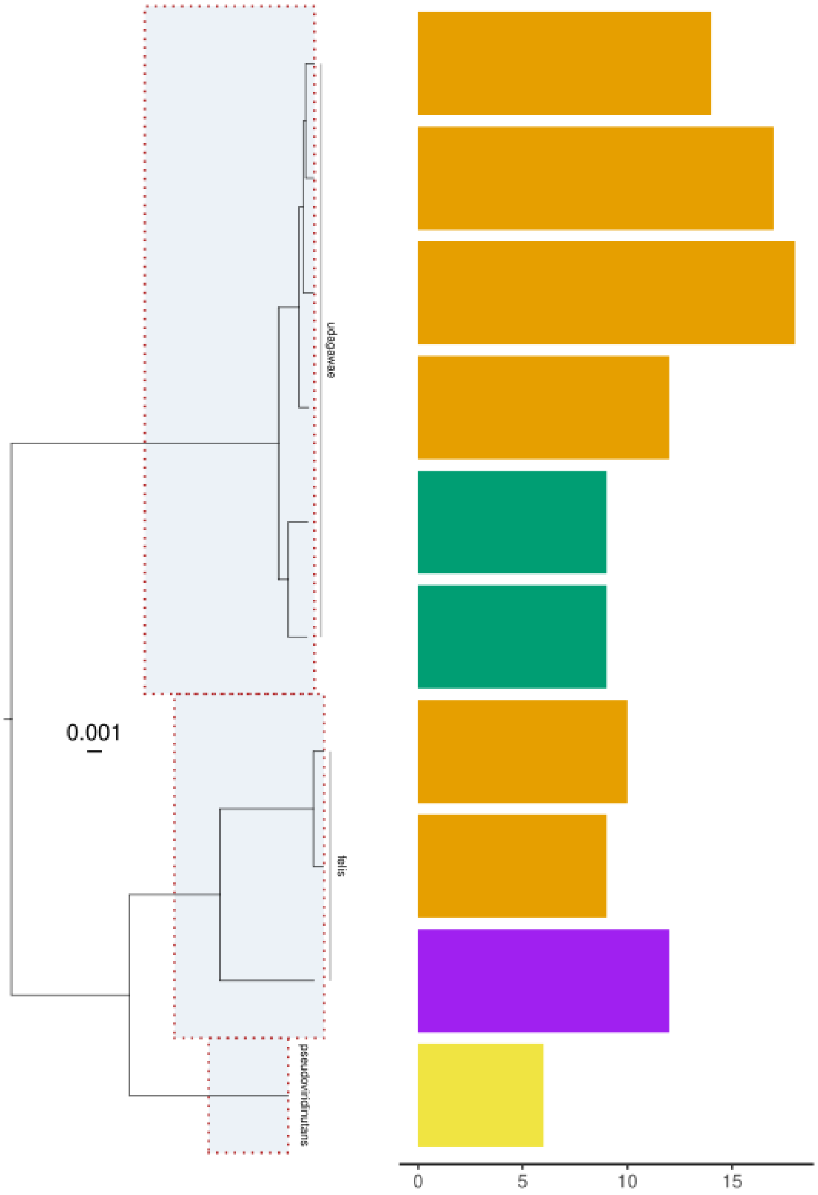


**fig. S18**. **A subset of assemblies (*A. udagawae*, *A. felis*, and *A. pseudoviridinutans*) phylogenetically placed alongside their captain count.** Bars are coloured according to isolation origins (clinical = orange; environment = green; environment (saline) = purple; environment-clinical = yellow; environment-food = light-blue; food-production=dark-blue; unknown=grey; NA=black).

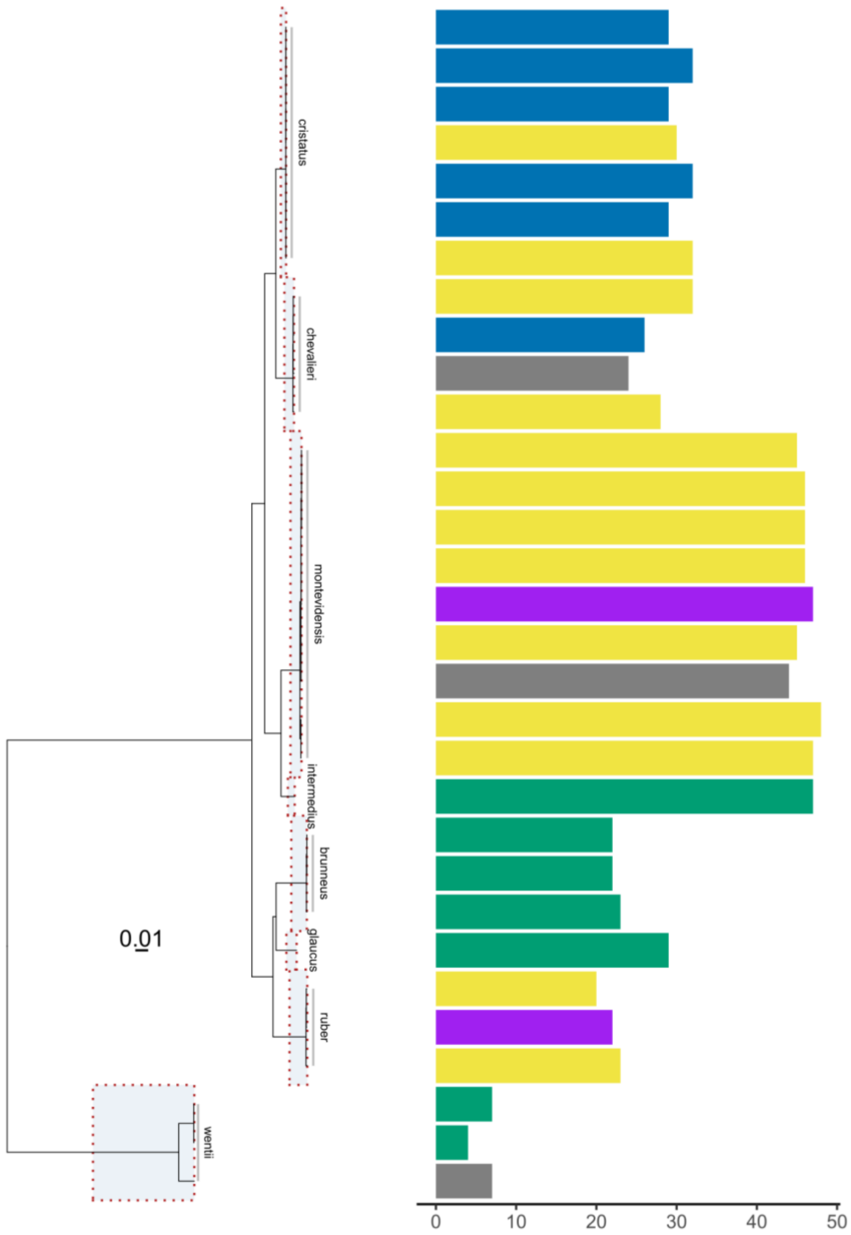


**fig. S19**. **A subset of assemblies (*A. cristatus*, *A. chevalieri*, *A. montevidensis, A. intermedius*, *A. brunneus, A. glaucus, A. ruber* and *A. wentii*) phylogenetically placed alongside their captain count**. Bars are coloured according to isolation origins (environment = green; environment (saline) = purple; environment-clinical = yellow; food-production=dark-blue; unknown=grey).

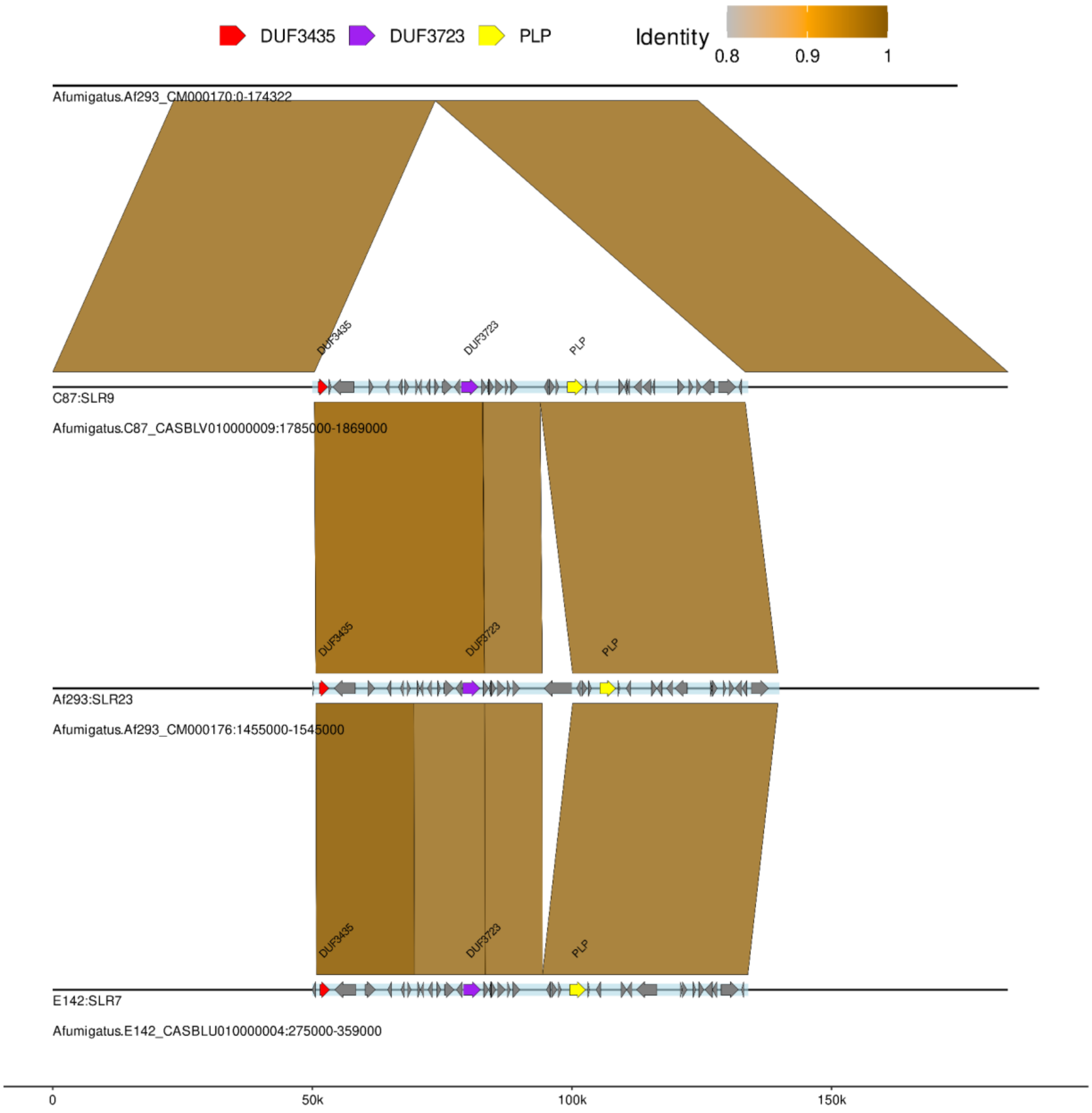


**fig. S20**. **Alignment of C87:SLR9 against a close relative without the SLR and two similar SLRs in other genomes**. Horizontal lines indicate the region of the genome visualised (as indicated in the name below the line) with only the SLR (light blue rectangle) and its annotated genes (arrows inside SLR) added as additional features. Starship-related genes are highlighted in different colours as indicated in the key above. In addition to the SLRs; 50kb up and downstream of the SLRs was used for alignment. Blocks between the horizontal lines indicate alignment with the colour displaying the 'identity' as a proportion of the alignment length to the number of matched bases. In this example the SLR is flanked by a DUF3435 gene and the alignment between each SLR lacks alignment at the flanks indicating this element is found in multiple unique positions The genome for Af293 here is used to show the insertion site of C87:SLR9 and the position of its similar SLR Af293:SLR23.

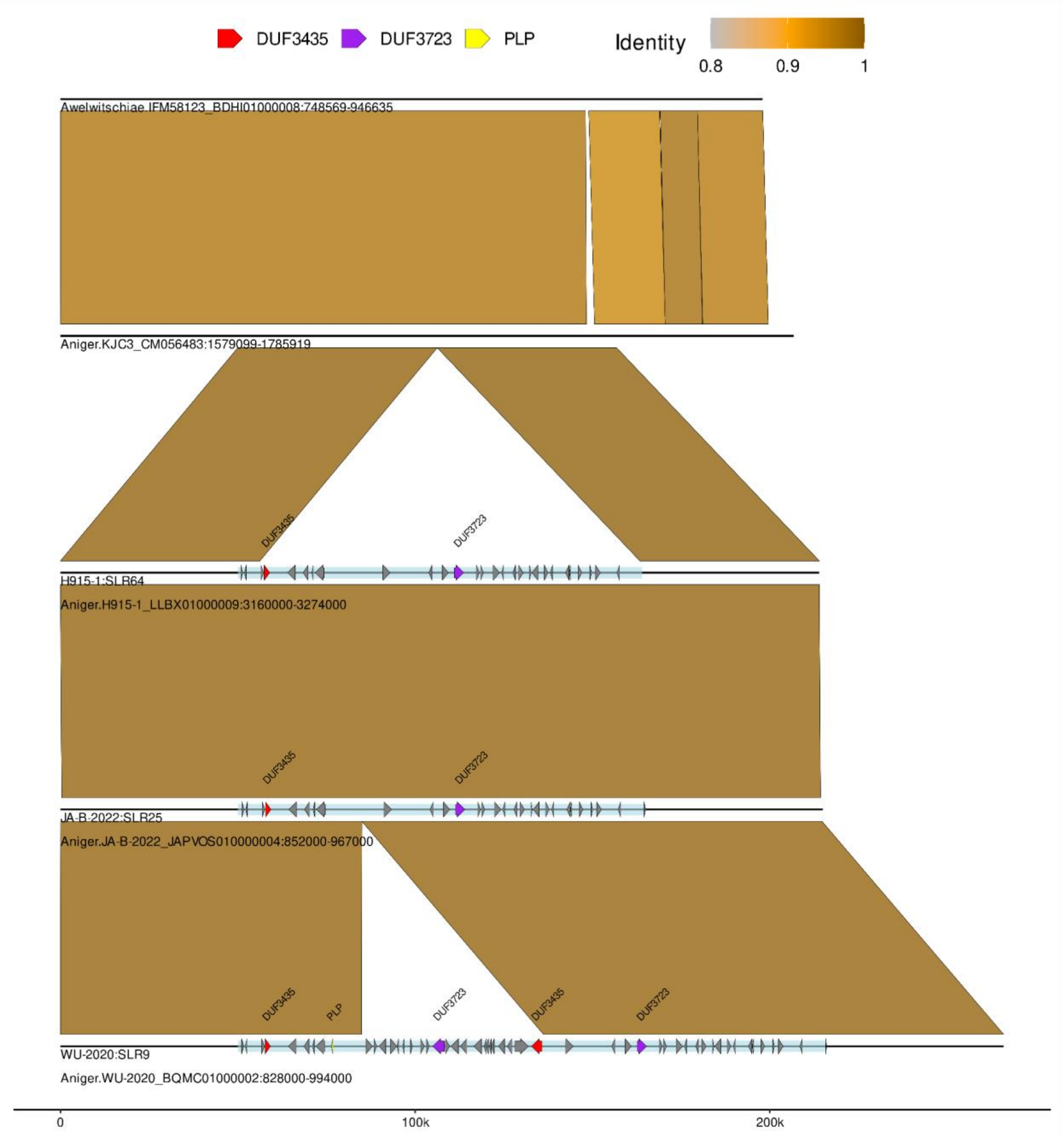


**fig. S21**. **Alignment of H915:SLR64 against close relatives without the SLR and two similar SLRs in other genomes**. Horizontal lines indicate the region of the genome visualised (as indicated in the name below the line) with only the SLR (light blue rectangle) and its annotated genes (arrows inside SLR) added as additional features. Starship-related genes are highlighted in different colours as indicated in the key above. In addition to the SLRs; 50kb up and downstream of the SLRs was used for alignment. Blocks between the horizontal lines indicate alignment with the colour displaying the 'identity' as a proportion of the alignment length to the number of matched bases. In this example all SLRs are flanked by a DUF3435 gene and are at the same insertion site; however WU-2020:SLR9 contains an additional nested *Starship* insertion.

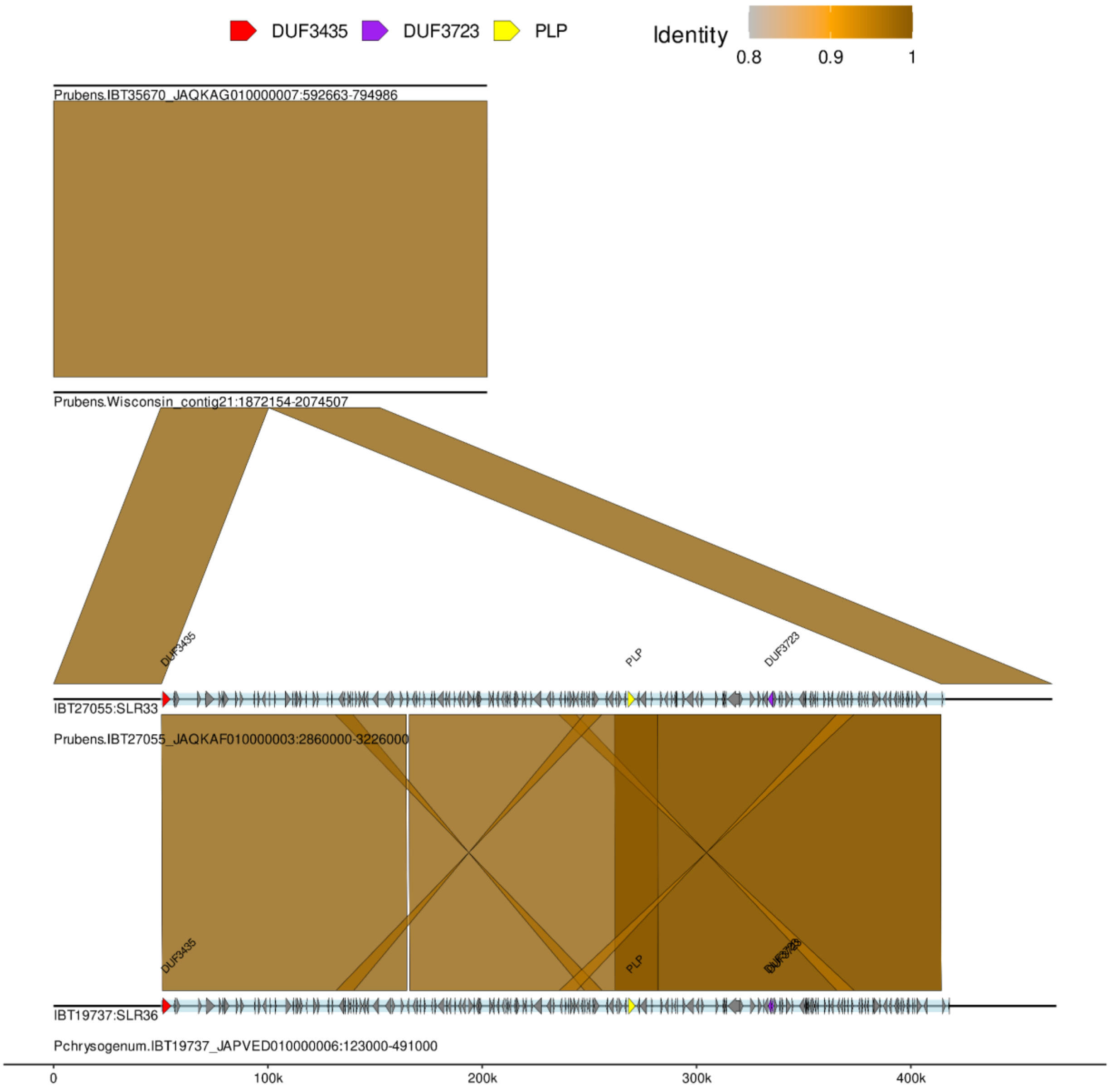


**fig. S22**. **Alignment of IBT27055:SLR33 against close relatives without the SLR and one similar SLR in another genome**. Horizontal lines indicate the region of the genome visualised (as indicated in the name below the line) with only the SLR (light blue rectangle) and its annotated genes (arrows inside SLR) added as additional features. Starship-related genes are highlighted in different colours as indicated in the key above. In addition to the SLRs; 50kb up and downstream of the SLRs was used for alignment. Blocks between the horizontal lines indicate alignment with the colour displaying the 'identity' as a proportion of the alignment length to the number of matched bases.In this example the SLR is flanked by a DUF3435 gene and the alignment between each SLR lacks alignment at the flanks indicating this element of nearly 400kb is found in multiple unique positions

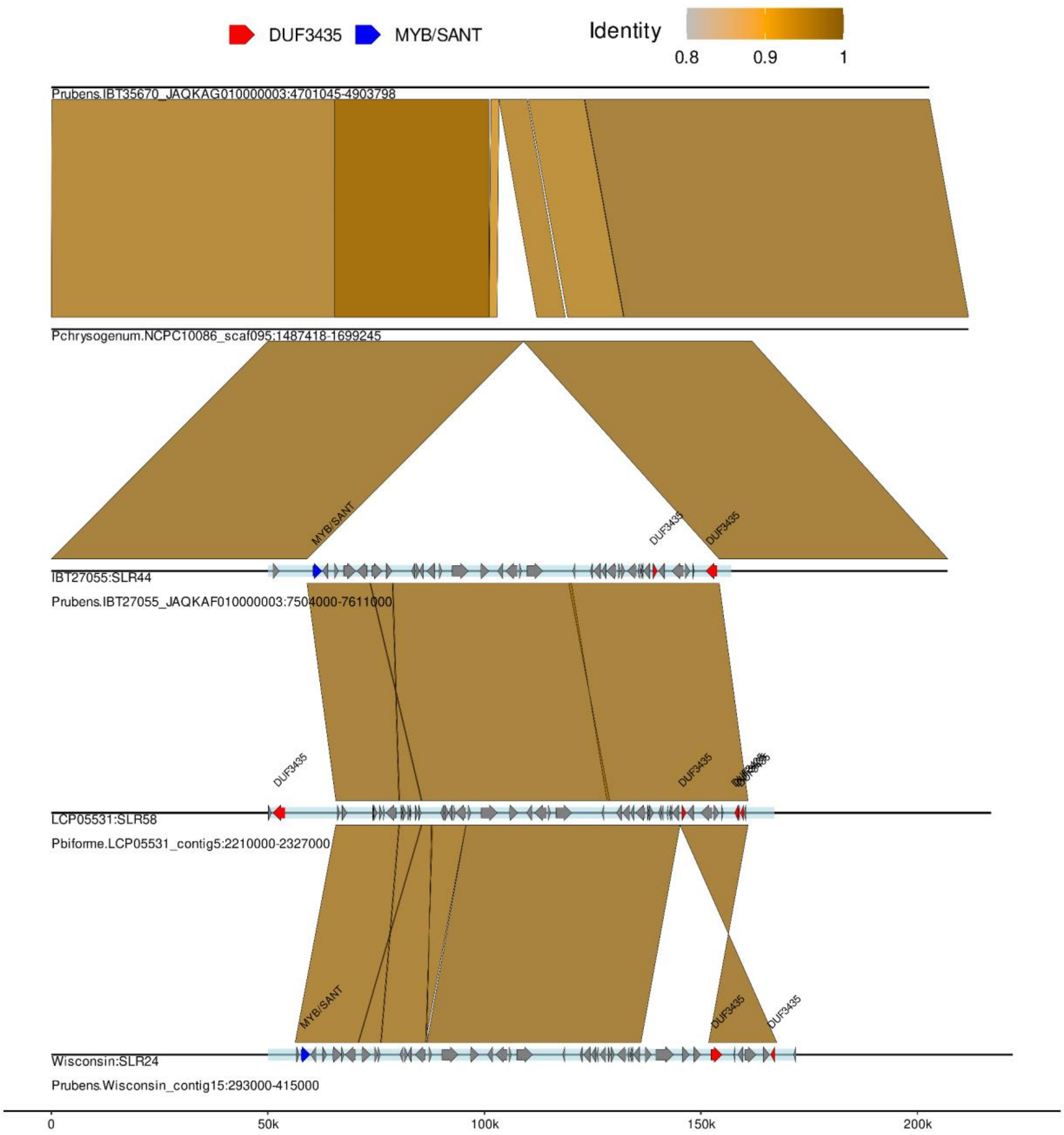


**fig. S23**. **Alignment of IBT27055:SLR44 against close relatives without the SLR and two similar SLRs in other genomes**. Horizontal lines indicate the region of the genome visualised (as indicated in the name below the line) with only the SLR (light blue rectangle) and its annotated genes (arrows inside SLR) added as additional features. Starship-related genes are highlighted in different colours as indicated in the key above. In addition to the SLRs; 50kb up and downstream of the SLRs was used for alignment. Blocks between the horizontal lines indicate alignment with the colour displaying the 'identity' as a proportion of the alignment length to the number of matched bases.In this example the SLR is flanked by a DUF3435 and MYB/SANT gene and the alignment between each SLR lacks alignment at the flanks indicating this element is found in multiple unique positions. The inversion in Wisconsin:SLR24 indicates a change in the captain found closest to the edge (a 'mutiny'). This *Starship* has been transferred between distant species; *Penicillium rubens* and *Penicillium biforme*. The additional DUF3435 gene in LCP05531:SLR58 highlights the sometimes complicated structures in SLRs.

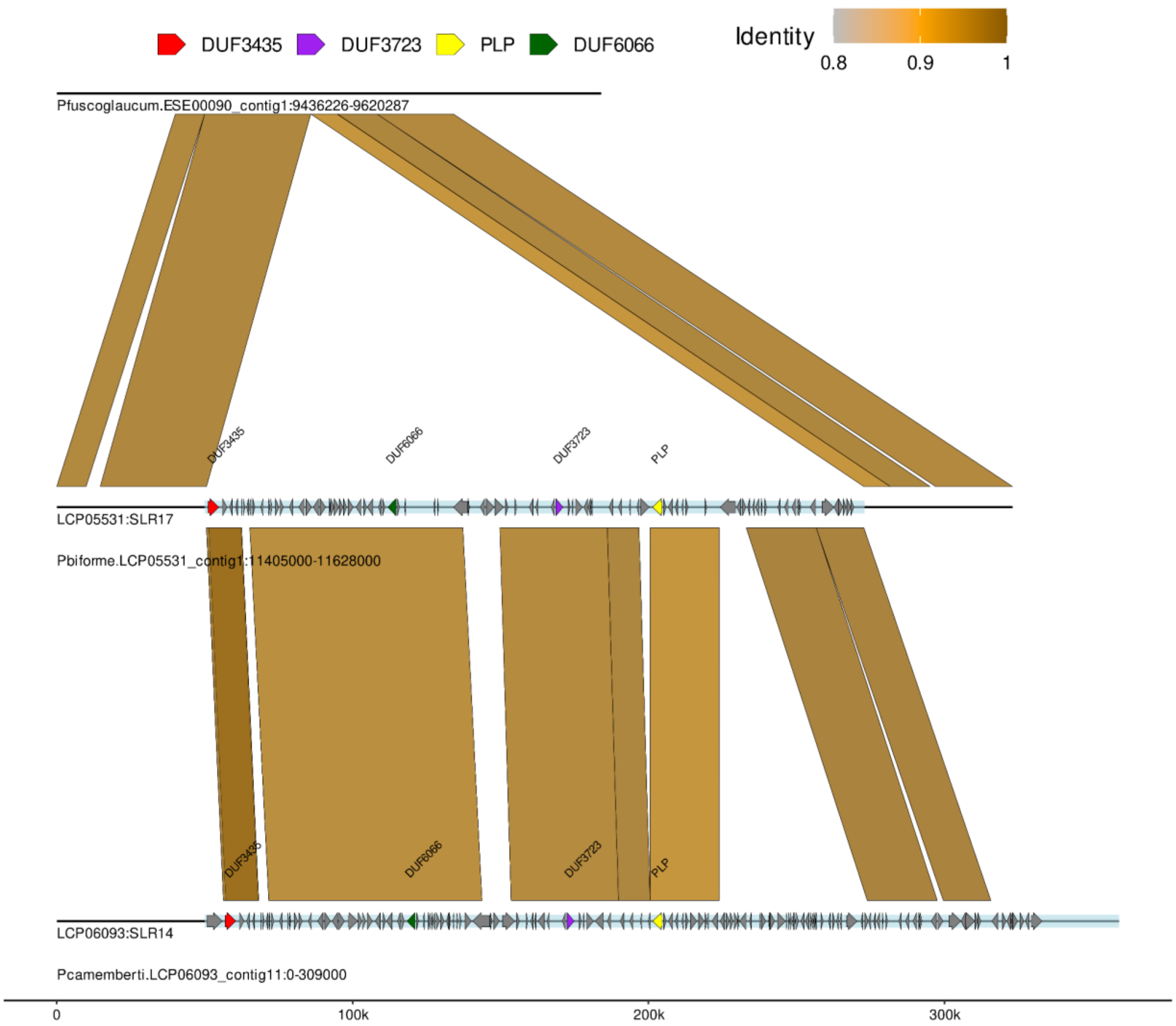


**fig. S24**. **Alignment of LCP05531:SLR17 against a close relative without the SLR and one similar SLR in another genome**. Horizontal lines indicate the region of the genome visualised (as indicated in the name below the line) with only the SLR (light blue rectangle) and its annotated genes (arrows inside SLR) added as additional features. Starship-related genes are highlighted in different colours as indicated in the key above. In addition to the SLRs; 50kb up and downstream of the SLRs was used for alignment. Blocks between the horizontal lines indicate alignment with the colour displaying the 'identity' as a proportion of the alignment length to the number of matched bases. In this example the SLR is flanked by a DUF3435 gene and the alignment between each SLR lacks alignment at the flanks indicating this element is found in multiple unique positions. The nonidentical structure highlights the structural differences found in many similar SLRs.

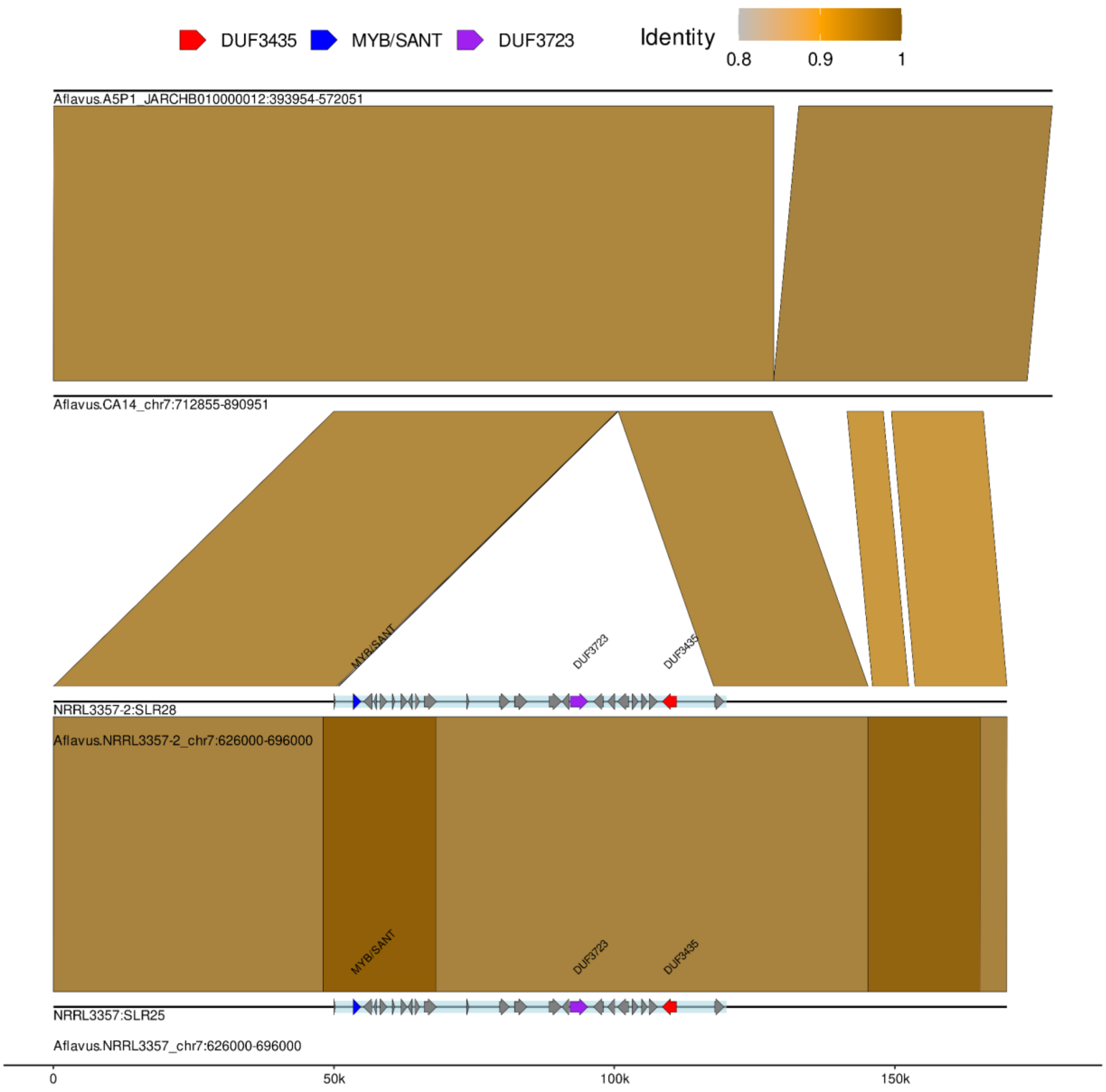


**fig. S25**. **Alignment of NRRL3357-2:SLR28 against close relatives without the SLR and one similar SLR in another genome**. Horizontal lines indicate the region of the genome visualised (as indicated in the name below the line) with only the SLR (light blue rectangle) and its annotated genes (arrows inside SLR) added as additional features. Starship-related genes are highlighted in different colours as indicated in the key above. In addition to the SLRs; 50kb up and downstream of the SLRs was used for alignment. Blocks between the horizontal lines indicate alignment with the colour displaying the 'identity' as a proportion of the alignment length to the number of matched bases. In this example the SLR is flanked by a DUF3435 and MYB/SANT gene and both SLRs are in the same position in genomes from the same *A. flavus* strain NRRL3357.

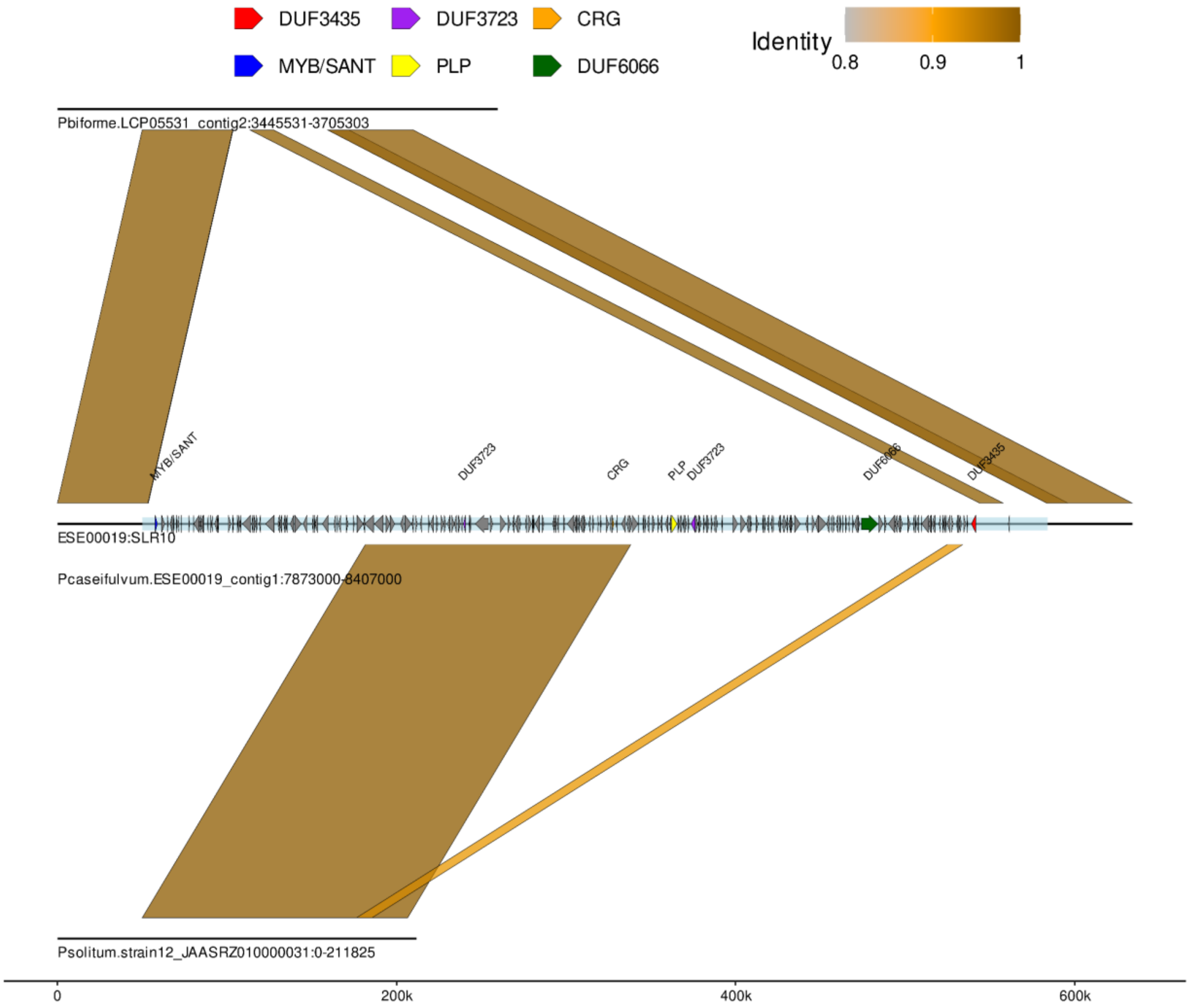


**fig. S26**. **Alignment of ESE00019:SLR10 against a close relative without the SLR and one similar SLR in another genome**. Horizontal lines indicate the region of the genome visualised (as indicated in the name below the line) with only the SLR (light blue rectangle) and its annotated genes (arrows inside SLR) added as additional features. Starship-related genes are highlighted in different colours as indicated in the key above. In addition to the SLRs; 50kb up and downstream of the SLRs was used for alignment. Blocks between the horizontal lines indicate alignment with the colour displaying the 'identity' as a proportion of the alignment length to the number of matched bases. In this example the >550kb SLR is flanked by both a DUF3435 and MYB/SANT gene. The SLR in *Penicillium camemberti var. caseifulvum* contains a similar SLR found in a distant species *Penicillium solitum* however this SLR only matches a large region of 150kb. By manual inspection we found the genome of *P. solitum* contains the remaining regions of the SLR in smaller contigs.

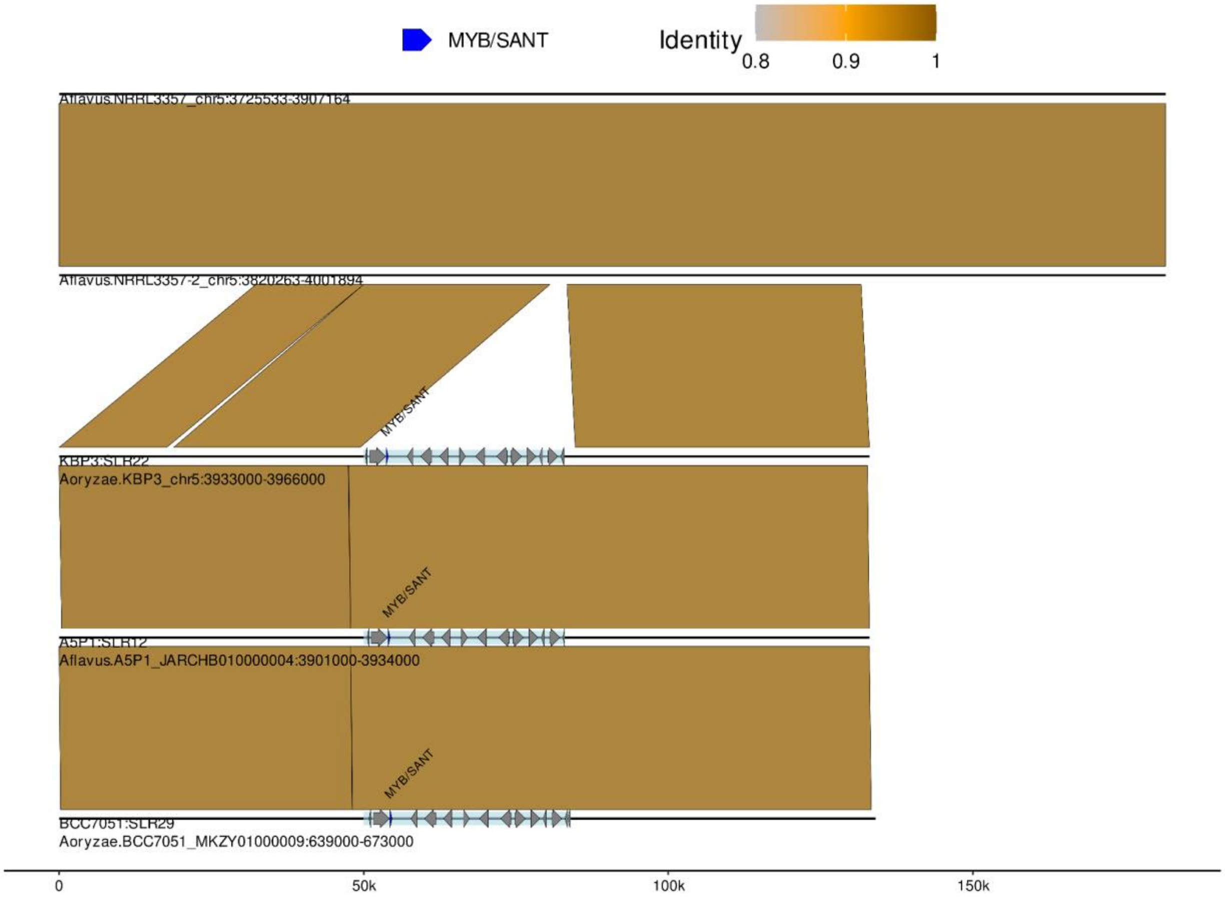


**fig. S27**. **Alignment of KBP3:SLR22 against close relatives without the SLR and two similar SLRs in other genomes**. Horizontal lines indicate the region of the genome visualised (as indicated in the name below the line) with only the SLR (light blue rectangle) and its annotated genes (arrows inside SLR) added as additional features. Starship-related genes are highlighted in different colours as indicated in the key above. In addition to the SLRs; 50kb up and downstream of the SLRs was used for alignment. Blocks between the horizontal lines indicate alignment with the colour displaying the 'identity' as a proportion of the alignment length to the number of matched bases. In this example the SLR is flanked by only a MYB/SANT gene and all SLRs are in the same position. The clear 30kb insertion only being flanked by a MYB/SANT gene indicates the utility of accessory cargo in identifying SLRs.

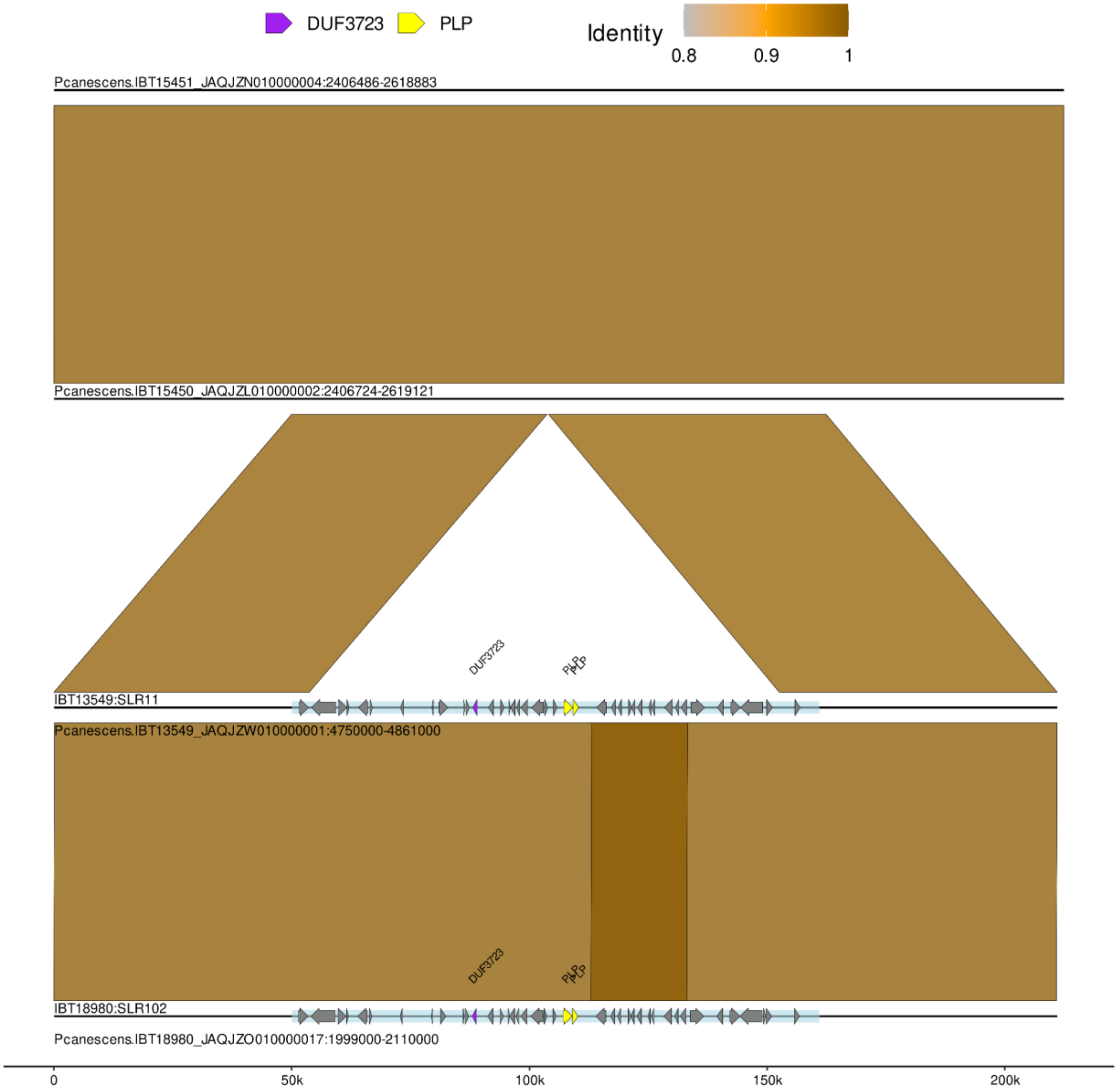


**fig. S28**. **Alignment of IBT13549:SLR11 against close relatives without the SLR and one similar SLR in another genome**. Horizontal lines indicate the region of the genome visualised (as indicated in the name below the line) with only the SLR (light blue rectangle) and its annotated genes (arrows inside SLR) added as additional features. Starship-related genes are highlighted in different colours as indicated in the key above. In addition to the SLRs; 50kb up and downstream of the SLRs was used for alignment. Blocks between the horizontal lines indicate alignment with the colour displaying the 'identity' as a proportion of the alignment length to the number of matched bases. In this example the SLR only contains accessory cargo genes and all SLRs are in the same position. The clear 150kb insertion only containing accessory cargo genes indicates their utility in identifying SLRs.

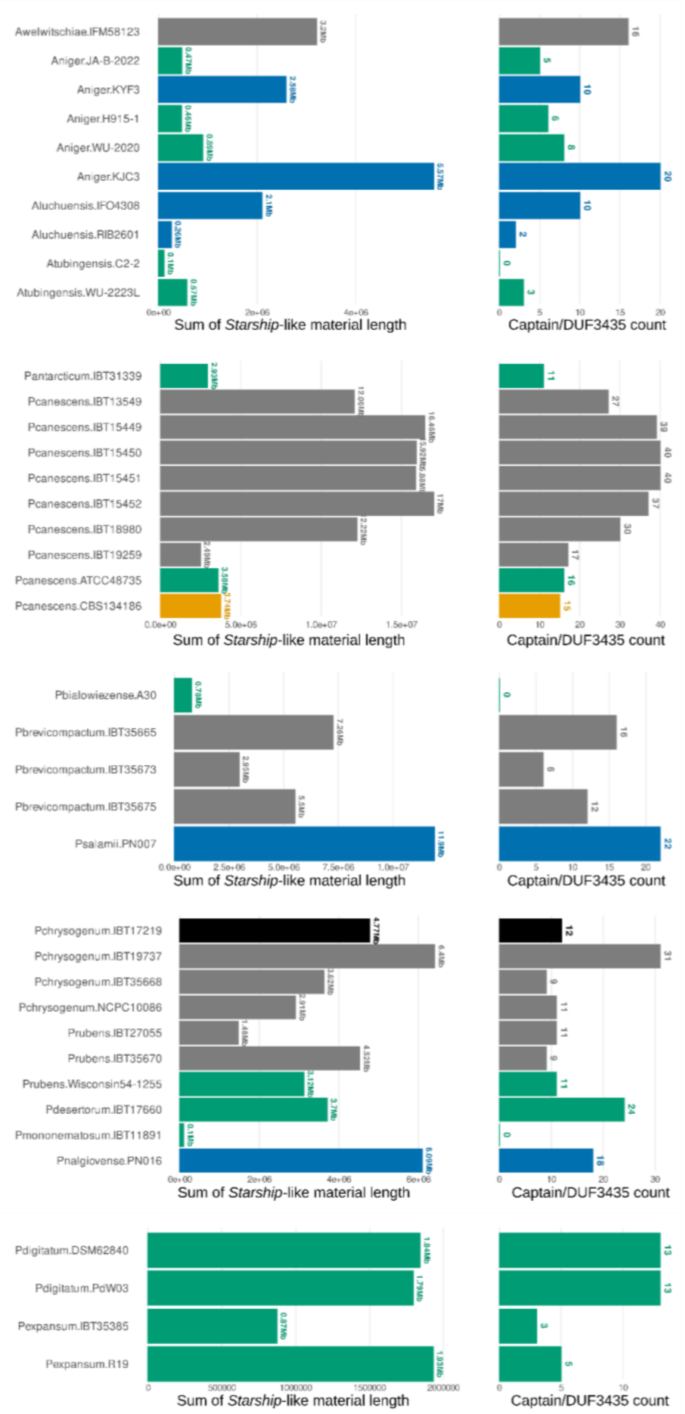


**fig. S29. The sum of *Starship*-related material and Captain count for each genome split by genome-graph**. Each set of genomes represents a genome-graph constructed with the assemblies for each strain present. The darker, black outlined column represents unique Starship-related material and Captains within them.

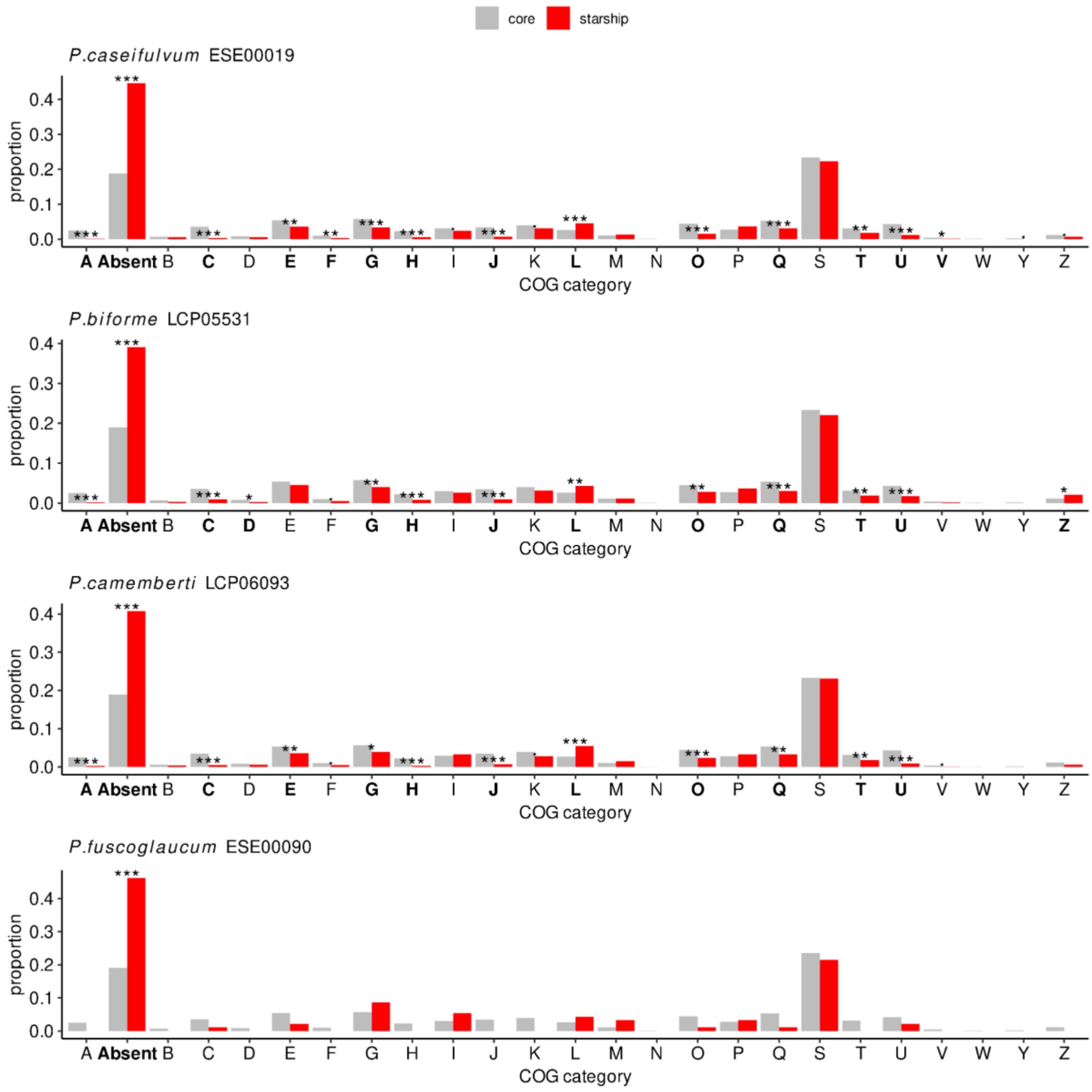


**fig. S30**. **The proportion of different COG annotations in cargo (red) and non-cargo (grey) subsets of four different genome assemblies. Those labelled as absent contained no COG annotation**. COG categories: A - RNA processing and modification; B - Chromatin structure and dynamics; C - Energy production and conversion; D - Cell cycle control, cell division, chromosome partitioning; E - Amino acid transport and metabolism; F - Nucleotide transport and metabolism; G - Carbohydrate transport and metabolism; H - Coenzyme transport and metabolism; I - Lipid transport and metabolism; J - Translation, ribosomal structure and biogenesis; K - Transcription; L - Replication, recombination and repair; M - Cell wall/membrane/envelope biogenesis; N - Cell motility; O - Posttranslational modification, protein turnover, chaperones; P - Inorganic ion transport and metabolism; Q - Secondary metabolites biosynthesis, transport and catabolism; R - General function prediction only; S - Function unknown; T - Signal transduction mechanisms; U - Intracellular trafficking, secretion, and vesicular transport; V - Defense mechanisms; W - Extracellular structures; Y - Nuclear structure; Z - Cytoskeleton. All p-values were calculated from hypergeometric tests and adjusted for multiple comparisons using Benjamini–Hochberg (· = p <= 0.1; * = p <= 0.05; ** = p <= 0.01; *** = p <= 0.001).

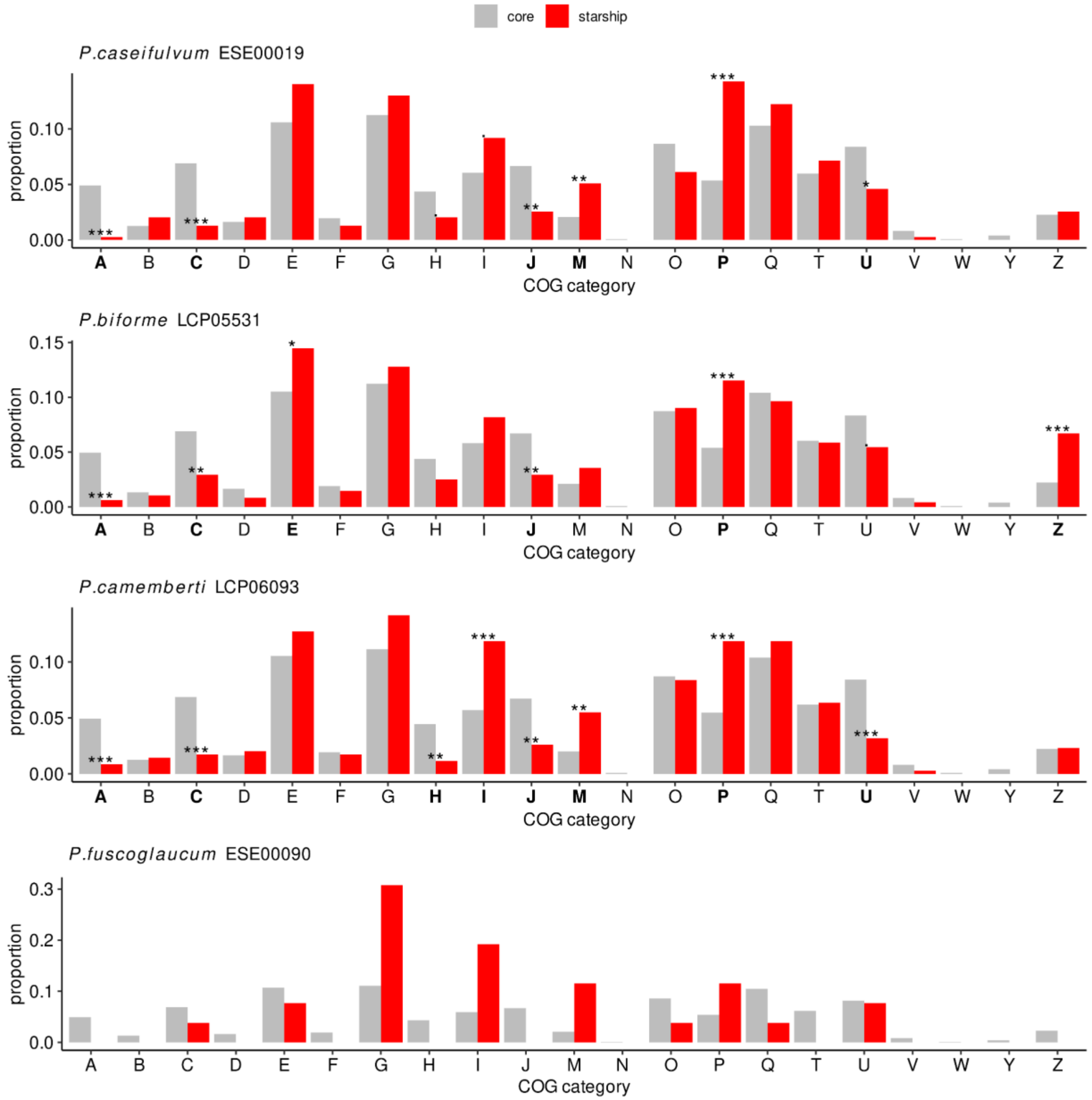


**fig. S31**. **The proportion of different COG annotations in cargo (red) and non-cargo (grey) subsets of four different genome assemblies after filtering for genes with no COG annotation and the COGs S, K and L**. COG categories: A - RNA processing and modification; B - Chromatin structure and dynamics; C - Energy production and conversion; D - Cell cycle control, cell division, chromosome partitioning; E - Amino acid transport and metabolism; F - Nucleotide transport and metabolism; G - Carbohydrate transport and metabolism; H - Coenzyme transport and metabolism; I - Lipid transport and metabolism; J - Translation, ribosomal structure and biogenesis; M - Cell wall/membrane/envelope biogenesis; N - Cell motility; O - Posttranslational modification, protein turnover, chaperones; P - Inorganic ion transport and metabolism; Q - Secondary metabolites biosynthesis, transport and catabolism; R - General function prediction only; T - Signal transduction mechanisms; U - Intracellular trafficking, secretion, and vesicular transport; V - Defense mechanisms; W - Extracellular structures; Y - Nuclear structure; Z - Cytoskeleton. All p-values were calculated from hypergeometric tests and adjusted for multiple comparisons using Benjamini–Hochberg (· = p <= 0.1; * = p <= 0.05; ** = p <= 0.01; *** = p <= 0.001).

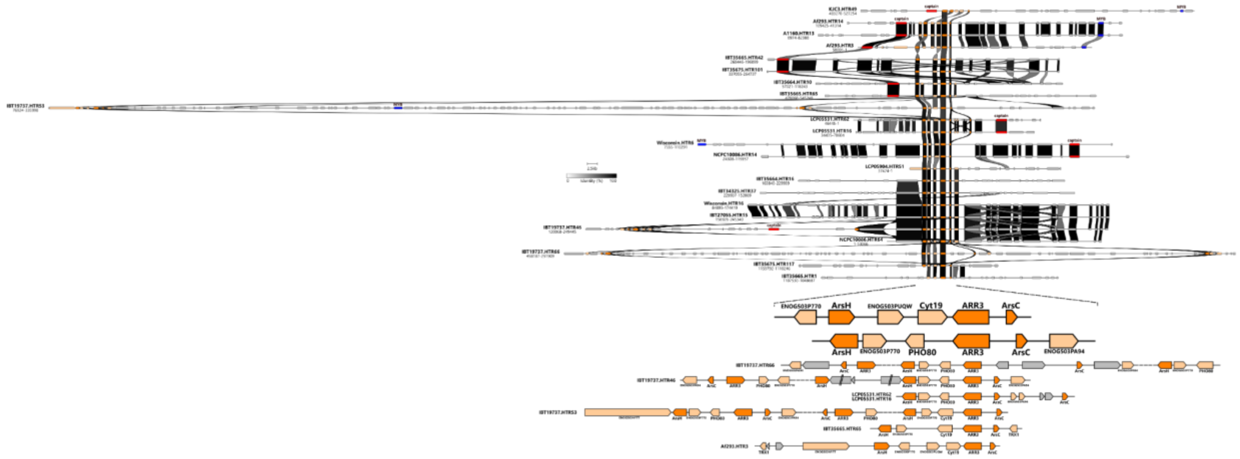


**fig. S32**. **Gene alignments in *Starship-like* regions centred around an Arsenic resistance cargo-cluster.** Genes within a subset of *Starship-like* regions were aligned and centred upon genes within the Arsenic resistance gene cluster. Core- and accessory-cluster genes are in dark and light orange respectively. Captains and MYB/SANT genes are labelled and coloured red and blue respectively. Colour of links between genes indicate pairwise identity as indicated in the scale. Examples at the bottom highlight several of the configurations of core and accessory genes.

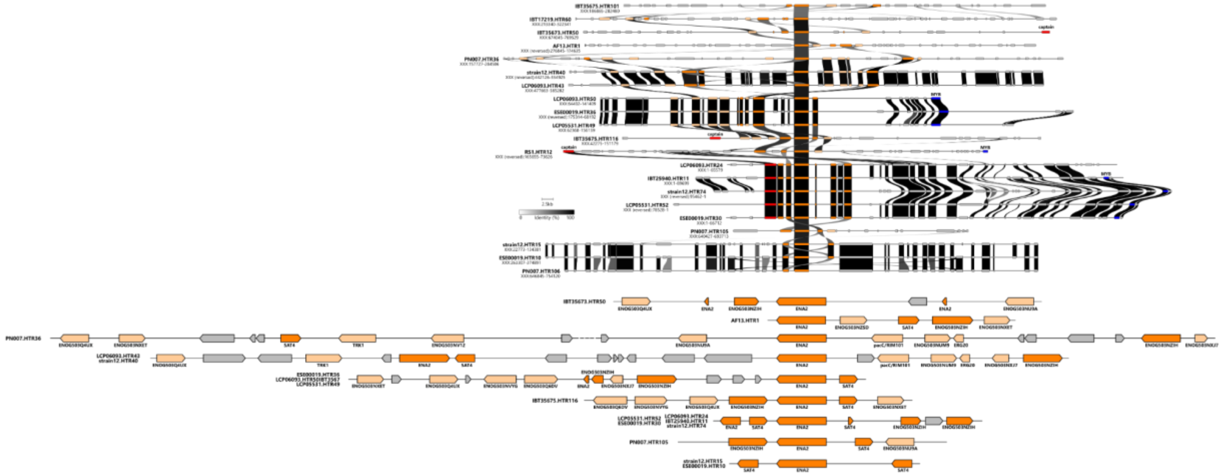

**fig. S33. Gene alignments in *Starship-like* regions centred around a salt-tolerance cargo-cluster.** Genes within a subset of *Starship-like* regions were aligned and centred upon those within the Salt-tolerance gene cluster. Core- and accessory-cluster genes are in dark and light orange respectively. Captains and MYB/SANT genes are labelled and coloured red and blue respectively. Colour of links between genes indicate pairwise identity as indicated in the scale. Examples at the bottom highlight several of the variable configurations of core and accessory genes.

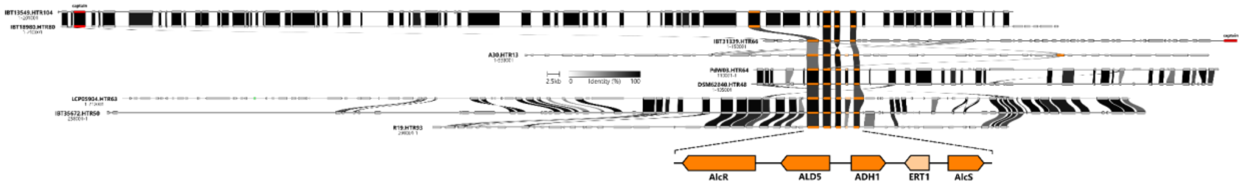


**fig. S34**. **Gene alignments in Starship-like regions centred around an ethanol utilisation cargo-cluster.** Genes within a subset of Starship-like regions were aligned and centred upon genes within the ethanol utilisation gene cluster. Core- and accessory-cluster genes are in dark and light orange respectively. Captains are labelled and coloured red. Colour of links between genes indicate pairwise identity as indicated in the scale.

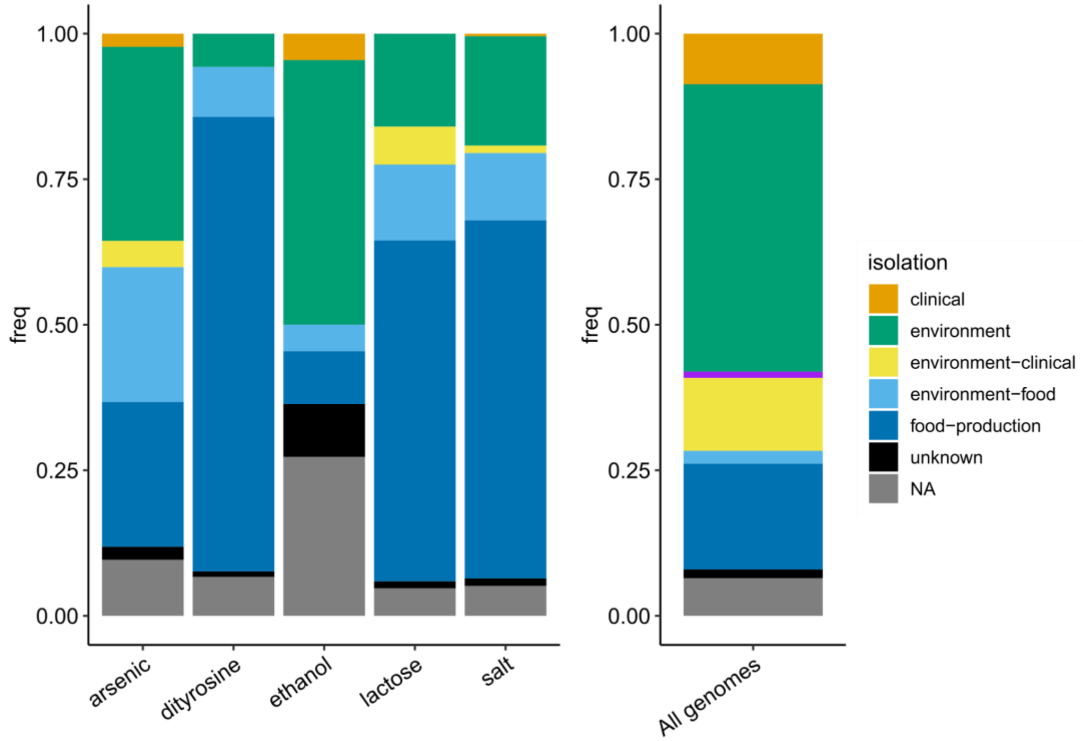


**fig. S35**. **Proportion of strains in total and with each cargo-cluster split by their isolation classification.**

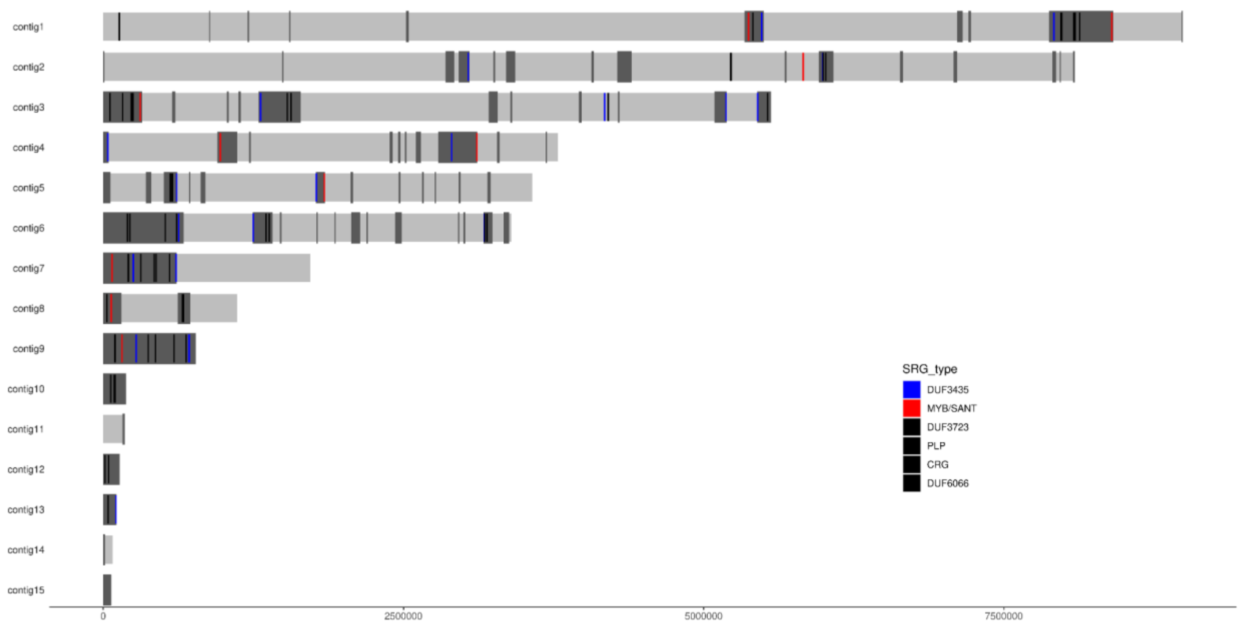


**fig. S36. Plotting the genome assembly of *P. caseifulvum* strain ESE00019 after pairwise comparison with the assembly of *P. fuscoglaucum* strain ESE00090 to identify regions absent from the latter.** Light grey boxes display contigs and dark grey boxes indicate regions missing from ESE00090. Starship-related genes are also plotted as to-scale rectangles. The rectangles for the DUF3435/Captain (red) and MYB/SANT TF (blue) are coloured while all other Starship-related genes are black.

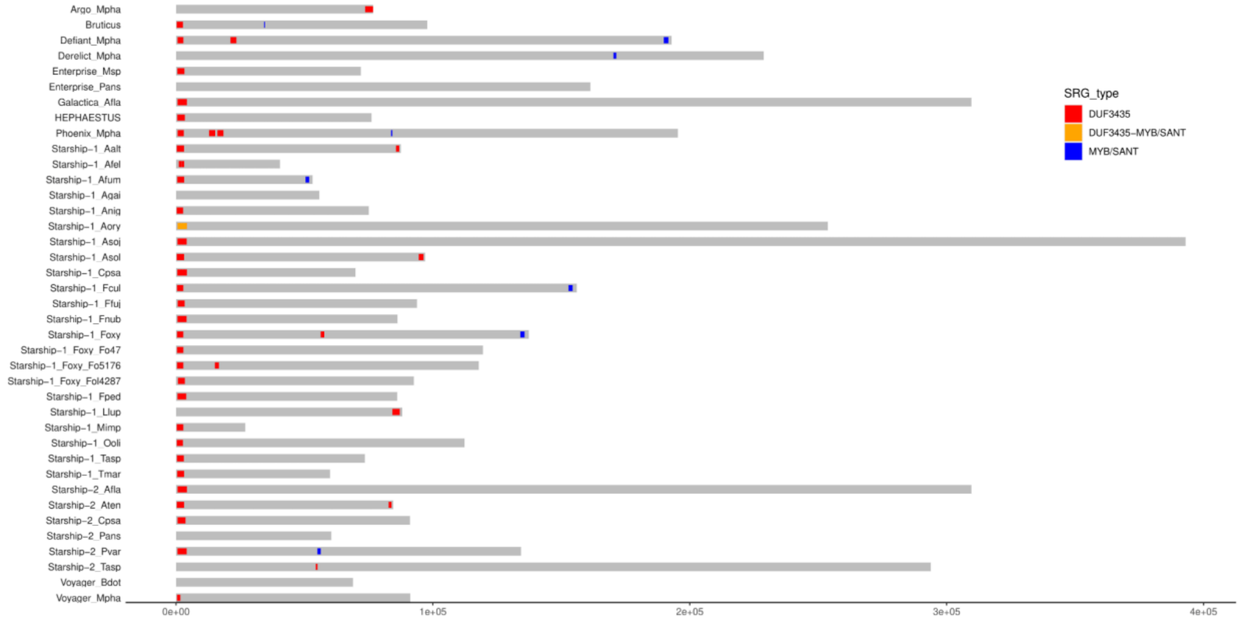


**fig. S37**. **Plotting Starships identified in Gluck-Thaler et al 2023 and identifying genes with protein domains related to DUF3435 and MYB/Sant genes**. Grey bars represent the Starships' named on the y-axis as given in Gluck-Thaler et al. Coloured blocks represent gene positions= MYB/Sant genes (blue), DUF3435 (red) and genes with both DUF3435 and MYB/SANT domains (orange).

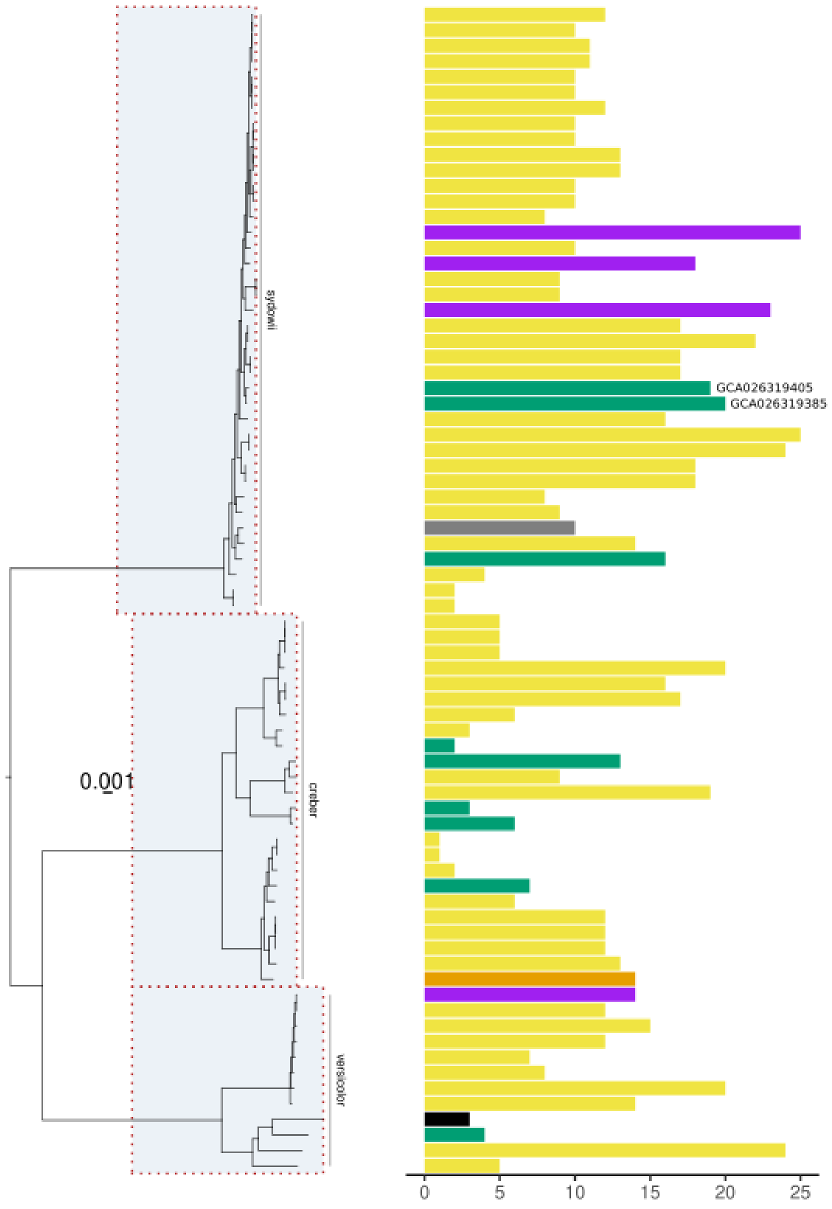


**fig. S38. A subset of assemblies (*A. sydowii*, *A. creber* and *A. versicolor*) phylogenetically placed alongside their captain count**. Bars are coloured according to isolation origins (environment = green; environment (saline) = purple; environment-clinical = yellow; clinical = orange; unknown=grey; NA= black). Two environmental genomes are labelled with their genome accession. Both were isolated from heavy metal contaminated soils.

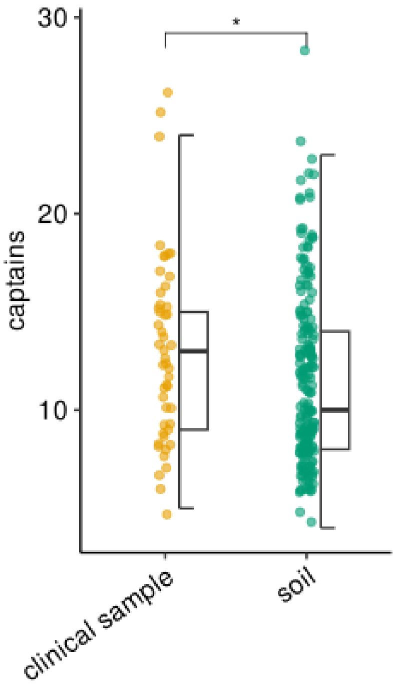


**fig. S39. A subset of *Aspergillus fumigatus* strains from Barber et al. 2020, 2021 studies tested for differences in the number of Captains separated by isolation origins**. P-values for the differences between groups were calculated using a Wilcoxon rank-sum test (* = 0.05 <= p >= 0.01).
